# Assessment of infant outgrowth of cow’s milk allergy in relation to the faecal microbiome and metaproteome

**DOI:** 10.1101/2022.11.03.514997

**Authors:** Diana M. Hendrickx, Ran An, Sjef Boeren, Sumanth K Mutte, PRESTO study team, Jolanda M Lambert, Clara Belzer

**Affiliations:** Laboratory of Microbiology, Wageningen University, Wageningen, The Netherlands; Laboratory of Biochemistry, Wageningen University, Wageningen, The Netherlands; Danone Nutricia Research, Utrecht, The Netherlands

**Author notes:** current address: Department of Food Science and Technology, School of Agriculture and Biology, Shanghai Jiao Tong University, Shanghai 200240, China (Ran An). A list of authors and their affiliations appears at the end of the paper.

## Abstract

Previous studies provide evidence for an association between modifications of the gut microbiota in early life and the development of food allergies. We studied the faecal microbiota composition (16S rRNA gene amplicon sequencing) and faecal microbiome functionality (metaproteomics) in a cohort of 40 infants diagnosed with cow’s milk allergy (CMA) when entering the study. Some of the infants showed outgrowth of CMA after 12 months, while others did not. Faecal microbiota composition of infants was analysed directly after CMA diagnosis (baseline) as well as 6 and 12 months after entering the study. The aim was to gain insight on gut microbiome parameters in relation to outgrowth of CMA.

The results of this study show that microbiome differences related to outgrowth of CMA can be mainly identified at the taxonomic level of the 16S rRNA gene, and to a lesser extent at the protein-based microbial taxonomy and functional protein level. At the 16S rRNA gene level outgrowth of CMA is characterized by lower relative abundance of *Lachnospiraceae* at baseline and lower *Bacteroidaceae* at visit 12 months.

## Introduction

The first three years of life are a key period for the development of the gut microbiome ^1^. In this period, two transition stages can be distinguished ^2^. During the first stage, immediately after birth, the gut microbiome is dominated by Proteobacteria and Actinobacteria ^3^. During the second stage, when infants start to eat solid foods until about three years of age, an adult-like microbiome, with Bacteroidetes and Firmicutes as most abundant phyla, is established ^4^.

Gut microbiome development during the first year of life is strongly influenced by various factors such as mode of delivery (vaginal/Caesarean) ^5^, antibiotics use ^6^, mode of feeding (breast fed/formula fed) ^7^ and lack of siblings ^8^.

Several studies provide evidence for an association between gut dysbiosis, induced by the factors mentioned above, and the development of food allergies (FA) ^9–11^. Previous research has shown that children with allergy have lower faecal DNA levels of *Bacteroides* ^12^, *Lactobacillus* ^12^ and *Bifidobacterium* ^13^ compared to healthy children. Furthermore, increased faecal DNA levels of *Clostridium difficile* ^14^, *Staphylococcus* ^12^ and *Escherichia coli* ^14^ have been associated with an increased risk of developing allergies. In children with cow’s milk allergy (CMA), lower faecal DNA levels of Bacteroidetes and higher levels of Proteobacteria were observed ^15^.

Metaproteomics approaches on faecal samples can provide insights into the functional roles of the intestinal microbiome in human diseases ^16^. However, they have seldom be applied to allergic diseases. One study applied metaproteomics on the infant gut microbiome to unravel functional signatures in infants with atopic dermatitis ^17^. To the best of our knowledge, metaproteomics has not been used to study the microbiome and its function in CMA. In this study, we considered both faecal metaproteomics and 16S rRNA gene amplicon sequencing data. Separate analysis was performed on each data type, but selection of core microbiota was based on combined rules applied to the two types of data. We study the changes in the microbiome of children diagnosed with CMA some of whom outgrew their CMA after one year and others did not. In this way, we aim to identify potential microbial factors associated with outgrowth of CMA.

## Methods

### Sample collection and experimental design

This study included 120 samples of the prospective interventional multicenter study PRESTO (NTR3725), retrieved from Danone Nutricia Research, where infants diagnosed with immunoglobulin E (IgE) mediated CMA received specialized nutrition aimed to induce outgrowth of CMA.

IgE-mediated CMA was diagnosed as follows as elsewhere ^18^. First, infants were considered as sensitized to cow’s milk (CM) if their CM-specific serum IgE was higher than 0.1 kU/L and/or a CM skin prick test (SPT) resulted in a wheal size ≥ 3 mm. Next, diagnosis of IgE-mediated CMA was confirmed by an open or double-blind placebo-controlled cow’s milk challenge or a history of anaphylaxis reaction to isolated ingestion of CM confirmed by two physicians.

Outgrowth of CMA at visit 12 months was evaluated through a double-blind placebo-controlled food challenge (DBPCFC) with CM powder, followed by an oral fresh milk challenge in case the former was negative, like described earlier ^18^.

Some infants received an amino acid-based formula (AAF), while others received an AAF with a synbiotic blend (AAF-syn) (oligosaccharides (oligofructose, inulin) + *Bifidobacterium breve M-16V*). Detailed information about the dosage has been described by Chatchatee et al ^18^. After 12 months, outgrowth of CMA was not different between infants that were fed with the AAF and those that were fed with the AAF-syn, and was in line with natural outgrowth of CMA ^18^.

The PRESTO study included 169 subjects, of which 40 were selected for our study based on sample, 16S rRNA gene sequencing and immunological data availability for a study duration up to 12 months. For those 40 infants, samples were collected at three visits, resulting in 120 samples: the baseline visit (0 months) where the subjects entered the study after CMA diagnosis, 6 months after entering the study and 12 months after entering the study. Faecal microbiota characterization was performed at the three visits for these 40 infants. All infants had been breastfed and were of age 3 - 13 months of age when entering the study (baseline visit). Of these 40 infants, 24 (10 infants in AAF group and 14 infants in AAF-syn group) outgrew their CMA after 12 months, while 15 infants (6 infants in AAF group and 9 infants in AAF-syn group) were still allergic to cow’s milk. For one infant the CMA status at visit 12 months was inconclusive, and this subject was removed from the analysis. The clinical characteristics of all infants included in this study are shown in Supplementary Table S1. For both metaproteomics and 16S rRNA gene amplicon sequencing 120 samples were collected, resulting in 117 samples after removing those of the infant whose allergy status at 12 months was inconclusive.

### Ethics approval

This multicenter study was designed and conducted in accordance World Medical Association (WMA) Declaration of Helsinki and the International Conference on Harmonization guidelines for Good Clinical Practice ^18^. The following national ethics committees, institutional review boards and regulatory authorities approved the study protocol and amendments:

- United Kingdom: NRES Committee North East - Sunderland (Central Ethics Committee MREC) and the local R&Ds from the following hospitals: Great Northern Children’s Hospital / Newcastle General Hospital; Southampton General Hospital; Guys & St Thomas; Barts / Royal London Hospital; Leicester Royal Infirmary
- Germany: Ethikkommission Charité – Ethikausschuss 2 am Campus Virchow-Klinikum; Ethikkommission Ärztekammer Nordrhein Düsseldorf, Ethik-Kommission der Medizinischen Fakultät der Ruhr Universität Bochum
- Italy: Comitato Etico per la Sperimentazione Clinica della Province della Pronvincia di Padova; Comitato Etico per la Sperimentazione Clinica della Province di Verona e Rovigo
- Singapore: Singhealth Centralised Institutional Review Board (CIRB) E; National Healthcare Group (NHG) Domain Specific Review Board
- Thailand: Institutional Review Board of the Faculty of Medicine, Chulalongkorn University; Committee on Human Rights Related to Research Involving Human Subjects, Faculty of Medicine Ramathibodi Hospital, Mahidol University; Ethics Committee of the Faculty of Medicine, Prince of Songkla University
- United States of America: Institutional Review Board of the Mount Sinai School of Medicine; Institutional Review Board for Human Subject Research for Baylor College of Medicine and Affiliated Hospitals (BCM IRB); University of Arkansas for Medical Sciences (UAMS) Institutional Review Board

Written informed consent for the collection and analysis of the data was obtained from the parents of all infants included in this study ^18^.

### Faeces sample collection and storage

Faeces samples for 16S-rRNA gene amplicon sequencing and metaproteomics were collected in 30-mL stool containers (Greiner 443102, Merck) at baseline, 6 months and 12 months. Aliquots were stored in 1.5 ml eppendorf tubes. Collection of stools took place between 2013 – 2018, and as a consequence the time gap between collection and analysis differs per sample. Faeces samples collected at home were immediately stored in home freezers of the parents and then transported with ice-packs to the hospitals within three months, where they were stored at −80 °C. Thereafter, faeces samples were transported from the hospitals to Danone Nutricia Research (the Netherlands) on dry-ice and stored at −80 °C until analysis. Transport of samples to LifeSequencing S.L. (Valencia, Spain) for 16S-rRNA gene amplicon sequencing, and to Wageningen University (the Netherlands) for metaproteomics was done in the same way. At Wageningen University, the samples for metaproteomics were stored at −80 °C until sample preparation. Peptide samples were stored at −20 °C until they were measured by nLC-MS/MS.

### DNA extraction for 16S-rRNA gene amplicon sequencing

DNA was extracted from faeces samples with QIAmp DNA Stool Mini Kit (Qiagen, Venlo, the Netherlands) according to the protocol of the manufacturer, but incorporating two extra bead-beating steps as described in Mischke et al^19^.

### 16S-rRNA gene amplicon sequencing

PCR amplification and sequencing of the V3-V4 region of the 16S rRNA gene was performed by LifeSequencing S.L. (Valencia, Spain). The 16S rRNA gene amplicons were sequenced using the 2 x 300 bp paired-end MiSeq protocol (Illumina).

The number of sequences (reads) per sample varied between 16830 and 66760. In total 713981 amplicon sequence variants were analyzed, of which 13309 were retained after applying the filtering steps described below.

### Sample preparation for metaproteomics by nLC-MS/MS

Total protein was isolated from faeces and subjected to metaproteomics preparation based on protein aggregation capture procedure for mass spectrometry (MS)^20^. Specifically, 10-30 mg faeces was mixed with 125 µl 100 mM Tris pH 8 (Duchefa Biochemie) and sonicated for 15 seconds with an MSE Soniprep 150 Needle sonicator (Beun de Ronde B.V.), at an amplitude of 22 µm. Twenty-five microliter of the slurry was centrifuged and the supernatant was subjected to protein concentration determination using Pierce TM BCA protein assay Kit (Thermo Fisher Scientific). The protein slurry was diluted to 1 ng/µl with 100 mM Tris pH 8. To 60 µl of the 1 ng/µl protein slurry, 150 mM dithiothreitol (Sigma Life Science) was added and incubated at 37 °C gently shaking for 45 min, followed by mixing with 198 µl 8 M urea (Sigma-Aldrich) and 27 µl 200 mM acrylamide (Sigma Life Science), and 30 min room temperature incubation. Subsequently, 4 µl of 10% trifluoroacetic acid (TFA, Alfa Aesar Chemicals), 8 µl of SpeedBeads™ magnetic carboxylate modified particles (50% GE Healthcare 45152105050250 and 50% Thermo Scientific 65152105050250 after two times washing with milli-Q water) and 750 µl acetonitrile (Biosolve B.V.) were added to the mixture, followed by 20 min room temperature incubation. The liquid was removed and beads with protein were washed twice with 1 ml 70% ethanol and with 1 ml 100% acetonitrile. Afterwards, beads with protein were subjected to overnight digestion by adding 100 µl 5 ng/µl sequencing grade trypsin (Roche Diagnostic GmbH) in ammonium bicarbonate (Sigma-Aldrich). Digestion was performed at room temperature while gently shaking overnight. The digestion was stopped by adding 4 µl 10% TFA to the mixture. The liquid was removed from the beads and transferred to a C18 µcolumn (description below). The remaining pellet was washed with 100 µl 1ml/L formic acid (Biosolve B.V.), which was also transferred to the µcolumn. After eluting all liquid through the µcolumn (while keeping the membrane not dry), 15 µl (1:1) formic acid & acetonitrile solution was added to the µcolumn and eluted the liquids. The filtered solution was then concentrated (Eppendorf Concentrator Plus) to a final volume of 10-15 µl, which volume was adjusted to exactly 50 µl with 1ml/L formic acid before storage in the −20°C freezer until analysis. The C18 µcolumn was made by adding two 1 mm pieces of C18 disk (Affinisep AttractSPE™ Disk Bio C18), 200 µl methanol (Hipersolv Chromanorm) and 4 µl 50% µcolumn material (Lichroprep RP-18) in methanol into a 200 µl pipette tip, which was eluted and washed with 100 µl methanol and equilibrated with 100 µl 1ml/L formic acid before usage.

### Metaproteomics nLC-MS/MS

Five microliter of peptide sample (defrozen and centrifuged at 12.000 * g for 30 min) was loaded directly onto a 0.10 * 250 mm ReproSil-Pur 120 C18-AQ 1.9 µm beads analytical column (prepared in-house) at a constant pressure of 825 bar (flow rate of circa 650 nL/min) with 1 ml/l HCOOH in water and eluted at a flow of 0.5 µl/min with a 50 min linear gradient from 9% to 34% acetonitril in water with 1 ml/l formic acid with a Thermo EASY nanoLC1000.

An electrospray potential of 3.5 kV was applied directly to the eluent via a stainless steel needle fitted into the waste line of a micro cross that was connected between the nLC and the analytical column.

A Field Asymmetric Ion Mobility Spectrometry (FAIMS) setup was used at a set compensation voltage of −45 V to increase the number of MSMS spectra obtained from doubly charged peptides. Full scan positive mode Fourier transform mass spectrometry (FTMS) spectra were measured between m/z 380 and 1400 on a Exploris 480 (Thermo electron, San Jose, CA, USA) at resolution (60000). MS and MSMS AGC targets were set to 300%, 100% respectively or maximum ion injection times of 50 ms (MS) and 30 ms (MSMS) were used. Higher-energy Collision dissociation (HCD) fragmented (Isolation width 1.2 m/z, 28% normalized collision energy) MSMS scans of the 25 most abundant 2-5+ charged peaks in the MS scan were recorded in data dependent mode (Resolution 15000, threshold 2e4, 15 s exclusion duration for the selected m/z +/- 10 ppm).

### 16S rRNA amplicon sequencing analysis

After demultiplexing the read pairs, low-quality sequences were removed by trimming the reads using a quality score (Q-score) threshold of 20. The trimmed reads were merged using PEAR ^21^, and merged reads with a minimal length of 300 were retained if they had a Q-score larger than 25 over a window of 15 bases and no ambiguous bases were present. Dereplication and counting of the merged reads was performed using mothur ^22^, and reads with < 2 reads over all samples were removed. Next, VSEARCH ^23^ together with the RDP gold database ^24^ were used to eliminate chimeras. Filtering of reads including PhiX and Adapter sequences (as defined in Deblur ^25^) was performed using QIIME2 ^26^. For each amplicon sequence variant (ASV), taxonomy was assigned at the genus level using the Ribosomal Database Project (RDP) ^27^ classifier against the SILVA 138 reference database ^28^, resulting in a classification of the ASVs into 173 genera.

### Database construction

ASVs were identified at the species level with NCBI nucleotide BLAST ^29^, using the 16S ribosomal RNA sequences (Bacteria and Archaea) database. The maximum number of aligned sequences was set to 10. For all other parameters the default settings were used.

For each ASV, the NCBI sequence with the highest percent identity was selected, without using any threshold. In case multiple NCBI sequences fulfilled this condition, they were all retained. The selected NCBI sequences were aggregated at the species level. For the resulting 327 species the non-redundant proteomes were obtained from UniProtKB ^30^. In case a species had no proteomes in UniProtKB, the NCBI proteome was obtained. For taxa with more than 2 proteomes in UniProtKB, two representative proteomes were selected. In case there was a reference proteome, this proteome was included and the second proteome was selected based on the Complete Proteome Detector (CPD) ^30^ and Benchmarking Universal Single-Copy Ortholog (BUSCO) ^31^ data completeness and quality scores. The resulting list of proteomes is provided in Supplementary Table S2. Including too many rare taxa found by 16S rRNA sequencing in a proteomics database can lead to a lower number of protein identifications. Therefore, a threshold for removing spurious taxa had to be defined. Genera were ordered from high to low relative abundance and multiple databases were constructed including the top genera that covered 50%, 66%, 75%, 80%, 90%, 92% and 100% of the total relative abundance in the 16S rRNA sequencing data. The database constructed using a threshold of 92% for the total relative abundance resulted in the highest number of protein identifications. To keep the database size limited yet complete, only the identified proteins (instead of complete proteomes) from the remaining 8% of the species were added to the database generated in the previous step. Since many proteins can be identical/redundant across multiple species/proteomes, we reduced the redundancy by clustering proteins that were identical using CD-HIT ^32^ with settings ‘-c 1.0 -d 500 -M 0’. The resulting non-redundant microbial proteomes database contained 416437 proteins.

### Metaproteomics data analysis

The obtained nLC-MS/MS spectra were analysed with MaxQuant version 2.0.3.0 ^33^ using the “Specific Trypsin/P” Digestion mode with maximally 2 missed cleavages and further default settings for the Andromeda search engine ^34^ (first search 20 ppm peptide tolerance, main search 4.5 ppm tolerance, MSMS fragment match tolerance of 20 ppm). Propionamide (C) was set as a fixed modification, while variable modifications were set for Protein N-terminal Acetylation and M oxidation with the maximum number of modifications per peptide set to 5. Next to the metaproteomics database described above under "database construction", a Human database downloaded from Uniprot (UP000005640 with 20400 sequences) and a contaminants database with 66 sequences of common contaminants were used. The “label-free quantification” as well as the “match between runs” options were enabled. De-amidated peptides were allowed to be used for protein quantification and all other quantification settings were kept default.

Proteins identified by only one or no unique peptide, reverse hits and hits only identified by site were filtered out. In this way we obtained 2705 protein groups, of which 2481 were from bacteria.

### Statistical analysis of 16S rRNA gene sequencing and metaproteomics data

All analyses (except LEfSe, see below) were performed in R version 3.6.1 ^35^. Clinical factors associated with outgrowth of cow’s milk allergy were determined by a two-sided Mann-Whitney U-test for numeric variables and a Fisher’s exact test for binary variables. For the microbiota analysis, ASVs were converted to relative abundances at the family level using the R microbiome package ^36^ version 1.8.0. For the metaproteomics data, relative abundances were calculated from the intensity Based Absolute Quantifcation (iBAQ) intensities.

As 16S rRNA gene-based taxonomic classification at genus level included a considerable fraction of unclassified *Lachnospiraceae* (Figure S1), statistical analysis was performed at the family level. Core microbiota were determined at the family level using the R microbiome package. Core taxa were defined as taxa that have a relative abundance higher than 1% in at least 50% of the 16S rRNA sequencing samples or at least 50% of the metaproteomics samples.

Bacterial proteins were functionally annotated by assigning protein identifiers to Kyoto Encyclopedia of Genes and Genomes (KEGG) ^37^ orthology (KO) identifiers, and classifying the KO identifiers using KEGG Brite hierarchy level C.

To compare relative abundances of core taxa, top 10 microbial protein functional classes and top 10 human protein classes within allergy groups and between visits, the data were normalized by centred log ratio (CLR) transform and a linear mixed model (LMM) analysis, with outgrowth of CMA, visit and outgrowth of CMA x visit as fixed effects and subject as random effect, was performed for each core taxon or protein class. The interaction term outgrowth of CMA x visit was included as we expected that visits compare differently at the level of outgrowth of CMA. The LMM was also used to determine differences in core taxa between allergy groups within each visit. P-values for pairwise comparisons were calculated from the model, and adjusted for multiple testing using the Benjamini-Hochberg procedure^38^. To study the effect of the wide age range of the infants when entering the study (3-13 months), we conducted an additional LMM analysis adding age as fixed effect. As adding more variables to the model can lead to overfitting, we also compared the model with and without age using Akaike Information Criterion (AIC).

To relate variability in the microbiome to environmental and clinical factors, redundancy analysis (RDA) was performed. Aitchison distance ^39^ (Euclidian distance on CLR transformed data), which has been proposed as a suitable distance metric for compositional data ^40^, was used for RDA. RDA analysis was performed with the R vegan package ^41^ version 2.5.-6, using stepwise forward and backward model selection where a variable is added when its p-value is smaller than or equal to 0.05, and removed when its p-value is larger than 0.1.

Furthermore, partial RDA with outgrowth of CMA as explanatory variable and the remainder of the environmental variables as covariates was performed to determine the proportion of variance in the microbiome explained by outgrowth of CMA. A similar RDA and partial RDA analysis was performed to relate variability in the microbial proteome and human proteins to environmental factors. Furthermore, another RDA was conducted to relate variability in the proteome and protein-based microbial composition to human proteins.

Spearman correlation between 16S gene-and protein-based taxonomic classification, and between human and microbial proteins, were calculated. P-values were determined with Monte Carlo permutation (10 000 permutations), and correlations with a p-value below 0.05 were considered significant.

Discriminative microbial features (taxa, microbial protein functional classes, human protein classes) between allergy groups (outgrowth versus no outgrowth of CMA at 12 months) and between visits were determined by Linear discriminant analysis Effect Size (LEfSe) analysis^42^, using the Galaxy module (http://huttenhower.sph.harvard.edu/galaxy). An alpha value of 0.05 for the factorial Kruskal-Wallis test among classes and a threshold of 2.0 on the logarithmic linear discriminant analysis (LDA) score for discriminative features were used for the calculations. To indicate which samples belong to the same subject in the pairwise comparisons between visits, a subject variable was included in the analysis.

LEfSe applies to relative abundances and focuses on effect size for selecting discriminative features. For comparison, we also included a method that applies to clr-transformed abundances and focuses on adjusted p-values rather than effect size. This method consists of fitting a linear mixed model (LMM) to all families or protein classes (instead of only core taxa or top 10 classes), using outgrowth of CMA, visit and outgrowth of CMA x visit as fixed effects and subject as random effect.

## Results

### Clinical factors associated with outgrowth of CMA

To determine environmental and clinical factors associated with outgrowth of CMA in our data set, we performed statistical tests (two-sided Mann-Whitney U-test for numeric variables, Fisher’s exact test for binary variables) to the metadata of the subjects used in this study (clinical characteristics, Supplementary Table S1).

Among infants with persistent CMA, a significantly higher proportion of infants with parental allergy is observed compared to infants that show outgrowth of CMA (Table S3 and Figure 1A). This suggests that persistent CMA is more common in case of parental allergy.

**Figure 1.**
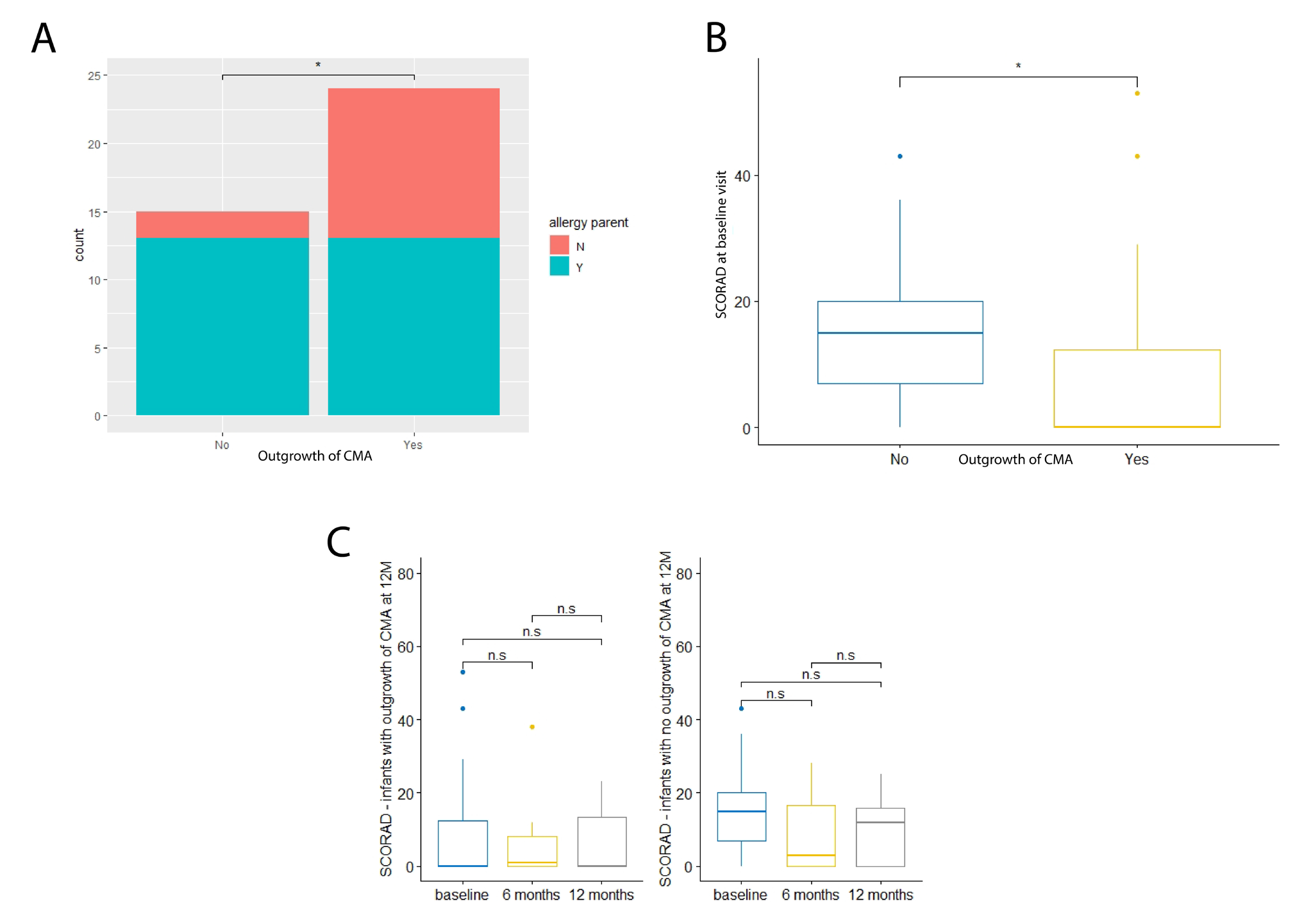
(A) Stacked bar plot for parental allergy. No = no outgrowth of CMA at visit 12 months; Yes = outgrowth of CMA at visit 12 months. (B) Boxplot for SCORAD at baseline visit. No = no outgrowth of CMA at visit 12 months; Yes = outgrowth of CMA at visit 12 months. (C) Boxplots for SCORAD over time. Left: outgrowth of CMA at 12M; right: no outgrowth of CMA at 12M. n.s.: not significant.

At baseline visit, a significantly lower score for atopic dermatitis (SCORAD) was observed in infants that show outgrowth of CMA compared to infants that do not (Table S3 and Figure 1B). The SCORAD did not change significantly over time in any of the two allergy groups (Figure 1C).

### The faecal metaproteome is dominated by bacterial proteins from Bifidobacteriaceae and Lachnospiraceae

Analysis of metaproteomics data was done by grouping the subjects into those that outgrew their CMA or those that did not at visit 12 months. The different visits were also analysed separately. For all groups, more than 40% of the total proteins are bacterial, the remaining proteins are human proteins (Figure 2A). The microbiota composition at the family level by metaproteomics showed high levels of *Bifidobacteriaceae* and *Lachnospiraceae* proteins (Figure 3A). Among the human proteins immunoglobulins are highly abundant (Figure 2B).

**Figure 2.**
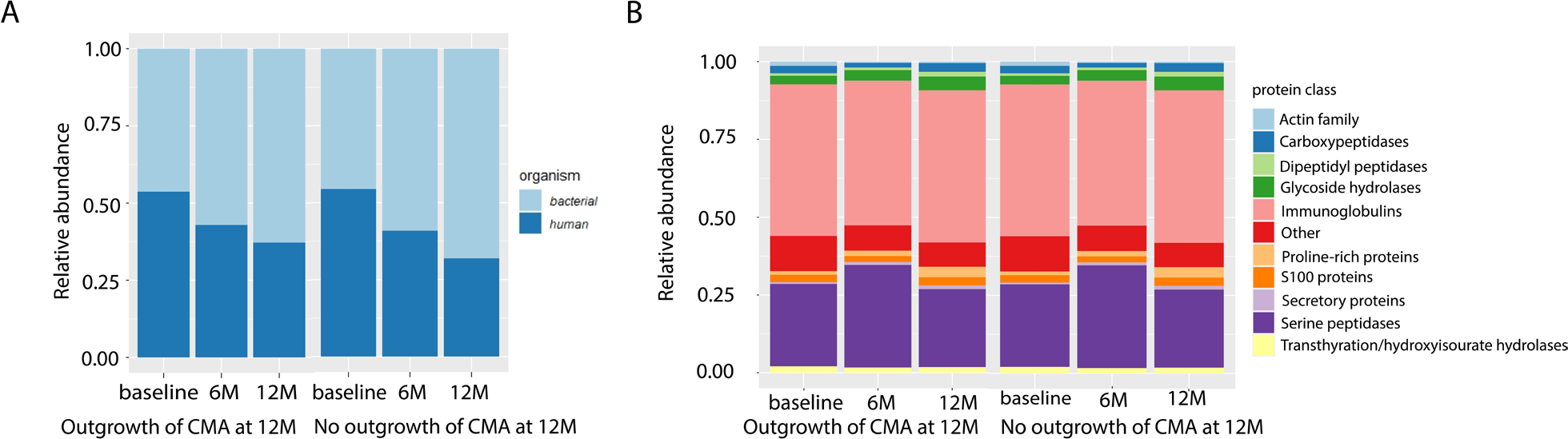
Ratio human/bacterial proteins (A) and human proteins composition (B) at each visit for the group that outgrew their CMA at visit 12 months (12M) and the group that did not.

**Figure 3.**
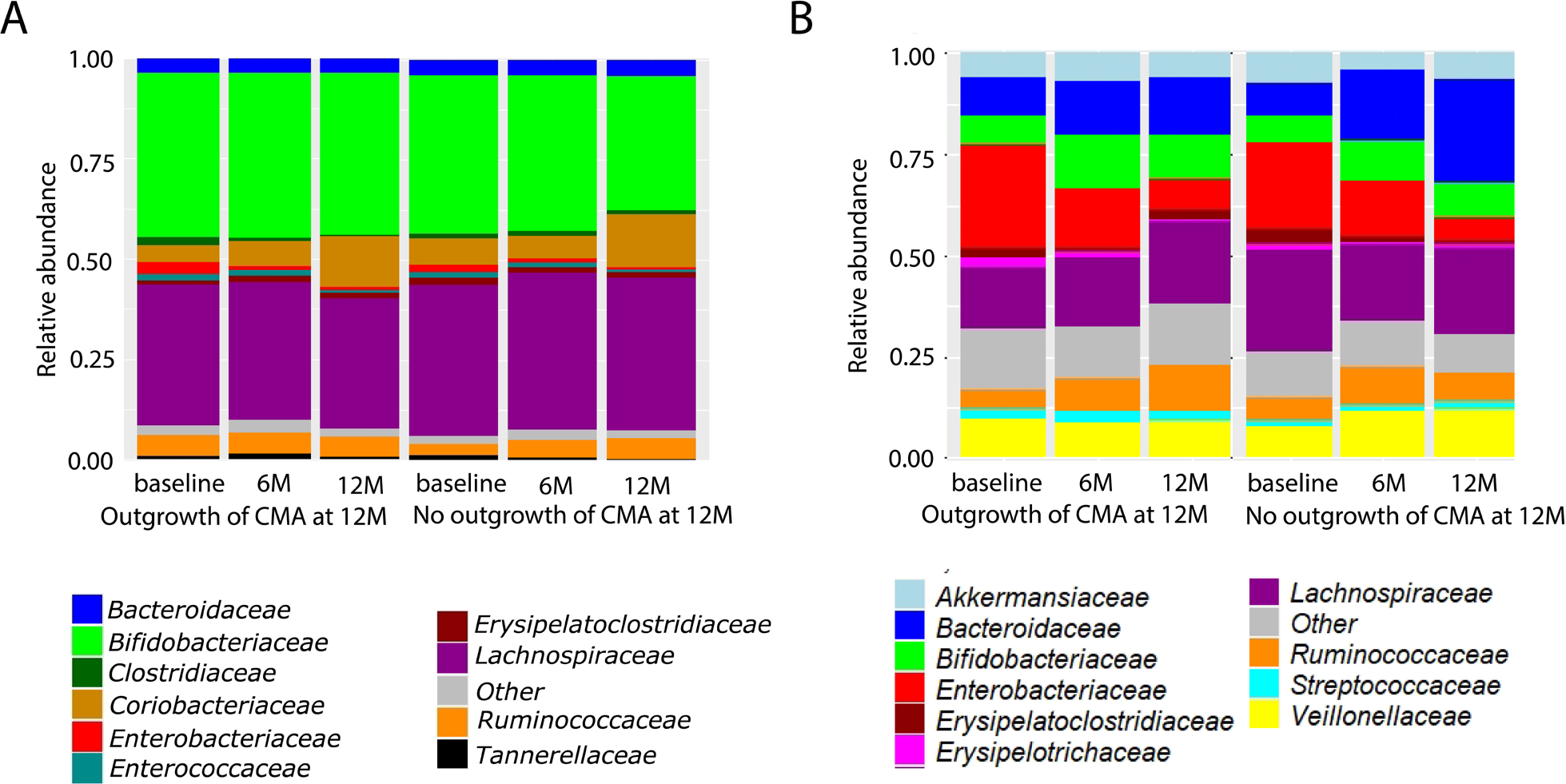
Protein-based microbial taxonomic profiles (family level) (A) and 16S rRNA gene-based taxonomic profiles (B) at each visit for the group that outgrew their CMA at visit 12 months (12M) and the group that did not.

### Differences and similarities between 16S rRNA gene-based and protein-based microbial taxonomic profiles

Metaproteomics-and 16S rRNA gene-based taxonomic profiles of the top 10 most abundant families across all samples were determined and compared between allergy groups (outgrowth of CMA versus no outgrowth of CMA at visit 12 months) and visits (Figures 3A and B).

#### Top 10 most abundant taxa in 16S rRNA gene and metaproteomics data

The top 10 most abundant families have 6 taxa in common between 16S rRNA gene-based and protein-based microbial taxonomic profiles. They are: *Bacteroidaceae*, *Bifidobacteriaceae*, *Enterobacteriaceae*, *Erysipelatoclostridiaceae*, *Lachnospiraceae* and *Ruminococcaceae* (Figure 3A and B). *Clostridiaceae*, *Coriobacteriaceae*, *Enterococcaceae* and *Tannerellaceae* were only among the top 10 taxa of protein-based microbial taxonomic profiles (Figure 3A). Taxa that were only among the top 10 in 16S rRNA gene-based profiles were *Akkermansiaceae*, *Erysipelotrichaceae* and *Streptococcaceae* and *Veillonellaceae* (Figure 3B). Both 16S rRNA-based and metaproteomics based taxonomic profiles show high inter-individual variation (Figures S2 and S3).

#### Correlation core taxa between 16S rRNA gene-based and protein-based microbial profiles

Core taxa, defined as taxa that have a relative abundance higher than 1% in at least 50% of the 16S rRNA gene sequencing or metaproteomics samples included *Bifidobacteriaceae*, *Coriobacteriaceae*, *Bacteroidaceae*, *Lachnospiraceae*, *Ruminococcaceae*, *Veillonellaceae* and *Enterobacteriaceae*. *Bifidobacteriaceae*, *Bacteroidaceae*, *Lachnospiraceae* and *Ruminococcaceae* fulfil these conditions for both 16S rRNA gene sequencing and metaproteomics samples, *Veillonellaceae* and *Enterobacteriaceae* only for 16S rRNA gene sequencing and *Coriobacteriaceae* only for metaproteomics. The relative abundance of *Bifidobacteriaceae*, *Bacteroidaceae*, *Lachnospiraceae*, *Ruminococcaceae* and *Enterobacteriaceae* correlated significantly (p-value ≤ 0.05) between 16S rRNA-based and protein-based microbial profiles (Table 1). However, apart from *Bifidobacteriaceae*, the correlations were low.

**Table 1.** Spearman correlation between 16S rRNA gene-and protein-based taxonomic classification. P-values were determined with Monte Carlo permutation (10 000 permutations). P-values below 0.05 are considered significant.

| Family | Spearman correlation | p-value |
| --- | --- | --- |
| *Bifidobacteriaceae* | 0.795 | < 1e-04 |
| *Coriobacteriaceae* | 0.142 | 0.1194 |
| *Bacteroidaceae* | 0.284 | 0.0024 |
| *Lachnospiraceae* | 0.539 | < 1e-04 |
| *Ruminococcaceae* | 0.518 | < 1e-04 |
| *Veillonellaceae* | 0.073 | 0.4268 |
| *Enterobacteriaceae* | 0.386 | < 1e-04 |

### Significant differences between allergy groups (outgrowth versus no outgrowth of CMA) in 16S rRNA gene-based relative abundance levels

To determine microbial variables most likely explaining differences between allergy groups in the 16S rRNA gene sequencing data, Linear Mixed Model (LMM) analysis on the core taxa and LEfSe and LMM analysis on the whole data set were performed. For all core taxa, the model without age as fixed effect shows the best fit to the data (lowest AIC)(Table S4). Below, differences at the family level are reported in detail, for differences at the higher taxonomy levels we refer to Figure 4.

**Figure 4.**
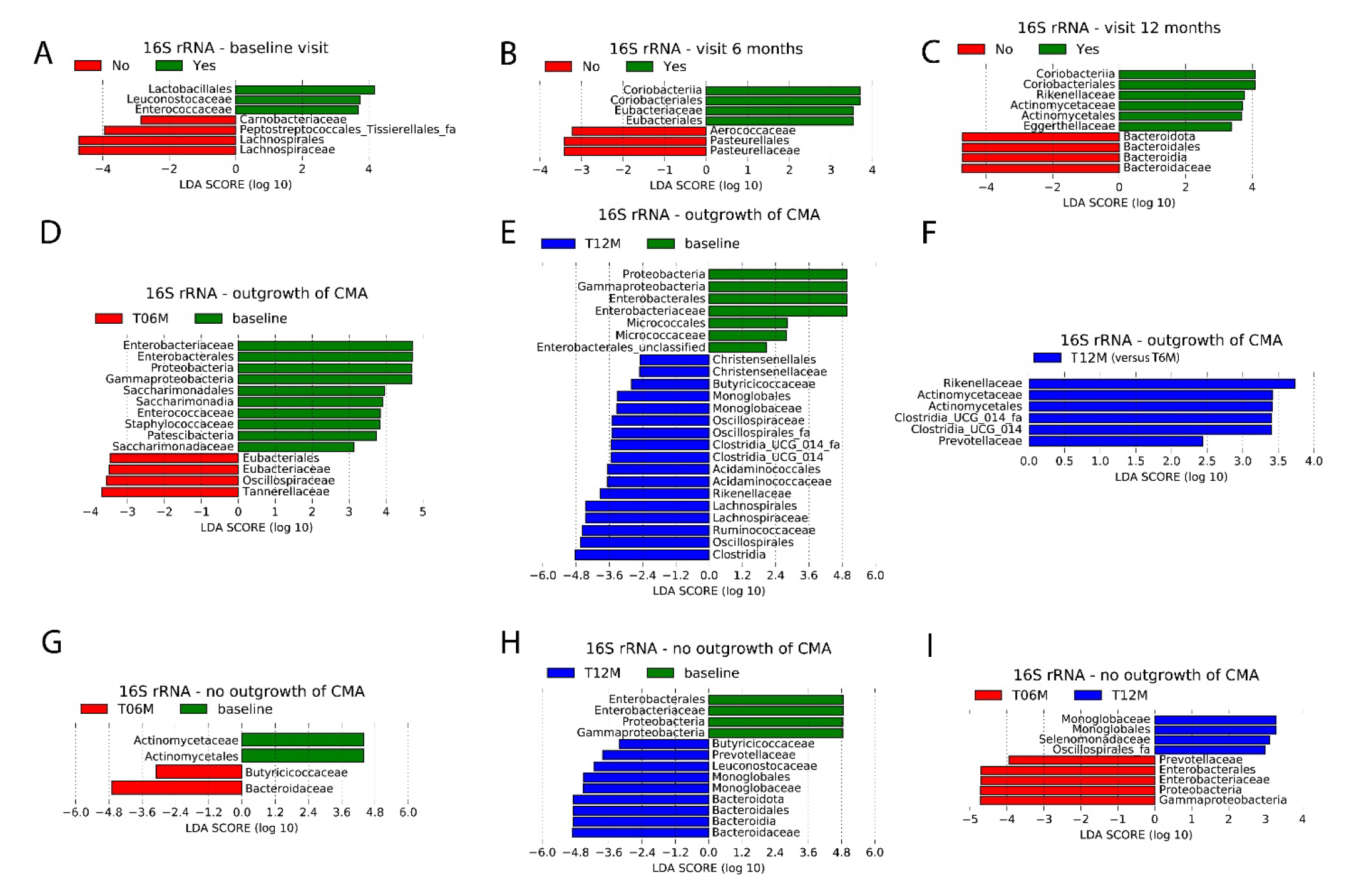
LEfSe analysis of 16S rRNA-based relative abundances at the family and all higher taxonomic levels using an alpha value of 0.05 for the factorial Kruskal-Wallis test among classes and a threshold of 2.0 on the logarithmic LDA score for discriminative features. (A-C) Using outgrowth of CMA at visit 12 months as class, Yes: outgrowth of CMA at 12 months; no: no outgrowth of CMA at 12 months. Plots of discriminative features at baseline visit (A), visit 6 months (B) and visit 12 months (C). (D-F) Pairwise comparison between visits within the group with outgrowth of CMA, using visit as class. (D) Visit 6 months (T06M) versus baseline; (E) visit 12 months (T12M) versus baseline; (F) visit 12 months (T12M) versus visit 6 months (T06M). (G-I) Pairwise comparison between visits within the group with persistent CMA, using visit as class. (G) Visit 6 months (T06M) versus baseline; (H) visit 12 months (T12M) versus baseline; (I) visit 12 months (T12M) versus visit 6 months (T06M).

#### Significant 16S rRNA gene-based differences between allergy groups within each visit

The results of both LMM analyses on the core taxa showed significantly higher relative abundances of *Lachnospiraceae* at baseline in infants with persistent CMA compared to infants who outgrew their CMA (Table S5). Furthermore, according the LEfSe analysis the families *Carnobacteriaceae*, *Lachnospiraceae* and *Peptostreptococcales-Tissierellales_fa* have significantly higher relative abundances at baseline visit in the group with persistent CMA (Figure 4A). In the group with outgrowth of CMA, higher relative abundances of *Enterococcaceae* and *Leuconostocaceae* were observed (Figure 4A). At visit 6 months, higher relative abundances of *Aerococcaceae* and *Pasteurellaceae* were observed in the group with persistent CMA, while in the group with outgrowth of CMA, higher relative abundances of the family *Eubacteriaceae* were observed (Figure 4B). At visit 12 months, higher relative abundances of *Bacteroidaceae* were observed in the infants with persistent CMA, while the infants who outgrew their CMA showed higher relative abundances of the families *Rikenellaceae*, *Actinomycetaceae* and *Eggerthellaceae* (Figure 4C).

The LMM analysis based on all taxa at the family level found no significant differences between allergy classes within visits after adjusting for multiple testing (Table S6). However, with the exception of *Eggerthellaceae* at 12 months, all discriminative features found by LEfSe were in the top 10 of LMM ordered by unadjusted p-value (Table S6). This indicates concordance between the LMM and LEfSe results, and suggests that the taxa obtained by LEfSe could have been significant when the LMM analysis could have been repeated in a larger cohort.

#### Significant 16S rRNA gene-based differences between visits within allergy groups

When comparing differences between visits within each allergy group, LMM analysis showed significantly higher relative abundances of *Ruminococcaceae* at visit 12 months compared to baseline visit in the group with outgrowth of CMA (Table S7 and Figure S4). In the group with persistent CMA, relative abundances of *Bacteroidaceae* were significantly higher at visit 12 months compared to baseline visit (Table S7 and Figure S4). Significant differences of *Enterobacteriaceae* were observed in both allergy groups. For the group with outgrowth of CMA, significantly lower relative abundances of *Enterobacteriaceae* were observed at visit 12 months compared to baseline visit, as well as at visit 6 months compared to baseline visit (Table S7 and Figure S4). In the group with persistent CMA, *Enterobacteriaceae* were only lower at visit 12 months compared to baseline (Table S7 and Figure S4). When including age as fixed effect in the LMM, we did not observe these effects.

In the group with outgrowth of CMA, the LEfSe analysis showed higher relative abundances of the families *Eubacteriaceae*, *Oscillospiraceae* and *Tannerellaceae*, and lower relative abundances of *Enterobacteriaceae*, *Enterococcaceae*, *Staphylococcaceae* and *Saccharimonadaceae* at visit 6 months compared to baseline (Figure 4D). Higher relative abundances of the families *Christensenellaceae*, *Butyricicoccaceae*, *Monoglobaceae*, *Oscillospiraceae*, *Oscillospirales_fa*, *Clostridia UCG 014* family, *Acidaminococcaceae*, *Rikenellaceae*, *Lachnospiraceae* and *Ruminococcaceae*, and lower relative abundances of *Enterobacteriaceae* and *Micrococcaceae* were observed at visit 12 months compared to baseline visit (Figure 4E). Higher relative abundances of the families *Rikenellaceae*, *Actinomycetaceae* and *Prevotellaceae* were observed at visit 12 months compared to visit 6 months (Figure 4F). The majority of families found by LEfSe in the comparisons 6 months versus baseline and 12 months versus baseline were also significant for the LMM analysis applied to all families (Table S8). For the comparison 12 months versus 6 months, the majority of the families found by LEfSe belongs to the top 10 ordered by unadjusted p-value found for the LMM analysis based on all families (Table S8). This suggests that these taxa could have been significant when the analysis could have been conducted in a larger cohort. At the family level, LEfSe analysis for the group with persistent CMA showed significantly higher relative abundances of *Butyricicoccaceae* and *Bacteroidaceae*, and significantly lower relative abundances of *Actinomycetaceae* at visit 6 months compared to baseline (Figure 4G). Higher relative abundances of the families *Butyricicoccaceae*, *Prevotellaceae*, *Leuconostocaceae*, *Monoglobaceae* and *Bacteroidaceae*, as well as lower relative abundances of *Enterobacteriaceae* were observed at visit 12 months compared to baseline (Figure 4H).

Moreover, higher relative abundance levels of the families *Monoglobaceae* and *Selenomonadaceae*, as well as lower relative abundance levels of *Prevotellaceae* and *Enterobacteriaceae* were observed at visit 12 months compared to visit 6 months (Figure 4I). The majority of the results obtained by LEfSe for this allergy group are not significant, but belong to the top 10 ordered by unadjusted p-value (Table S9).

### Significant differences between allergy groups (outgrowth versus no outgrowth of CMA) in protein-based microbial relative abundance levels

Linear Mixed Model (LMM) and LEfSe analysis were performed to determine microbial variables most likely explaining differences between allergy groups in the metaproteomics data. Similar as for the models based on 16S rRNA gene-based relative abundance levels, the models without age as fixed effect show the best goodness of fit (lowest AIC)(Table S10). Below, differences at the family level are reported in detail, for differences at the higher taxonomy levels we refer to Figure 5.

**Figure 5.**
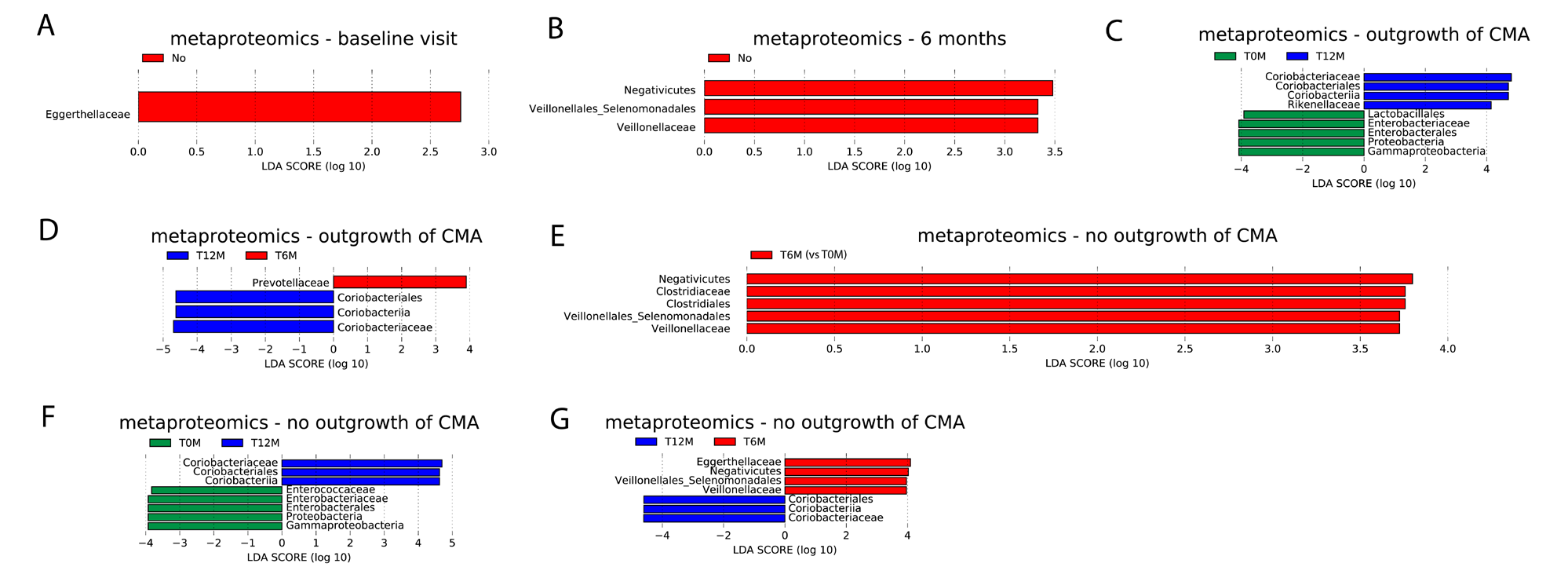
LEfSe analysis of protein-based microbial relative abundances at the family and all higher taxonomic levels using an alpha value of 0.05 for the factorial Kruskal-Wallis test among classes and a threshold of 2.0 on the logarithmic LDA score for discriminative features. (A-B) Using outgrowth of CMA at visit 12 months as class, Yes: outgrowth of CMA at 12 months; no: no outgrowth of CMA at 12 months. Plot of discriminative features at (A) baseline visit; (B) visit 6 months. (C-D) Pairwise comparison between visits within the group with outgrowth of CMA, using visit as class. (C) visit 12 months (T12M) versus baseline (T0M); (D) visit 12 months (T12M) versus visit 6 months (T6M). (E-G) Pairwise comparison between visits within the group with persistent CMA, using visit as class. (E) visit 6 months (T6M) versus baseline (T0M); (F) visit 12 months (T12M) versus baseline (T0M); (G) visit 12 months (T12M) versus visit 6 months (T6M).

#### Significant protein-based microbial differences between between allergy groups within each visit

Both LMM analyses did not identify any significant differences in core taxa between allergy groups within visits (Table S11). The results of the LEfSe analysis show higher relative abundances of *Eggerthellaceae* proteins at baseline and higher relative abundances of *Veillonelleceae* proteins at visit 6 months in infants with persistent CMA compared to infants who outgrew their CMA (Figure 5A and B). For visit 12 months, no significant differences between allergy groups were observed.

The LMM analysis based on all taxa at the family level found no significant differences between allergy classes within visits after adjusting for multiple testing (Table S12). However, *Eggerthellaceae* has the lowest uncorrected p-value for baseline visit, and *Veillonellaceae* has the second lowest uncorrected p-value for 6 months (Table S12). This suggest that these taxa could have been found significant by LMM analysis in a larger cohort with more statistical power.

#### Significant protein-based microbial differences between visits within allergy groups

In infants who outgrew their CMA, the LMM analysis comparing visits identified significantly lower relative abundance of *Enterobacteriaceae* proteins at visit 12 months compared to the baseline visit (Table S13 and Figure S5). This effect was not observed when repeating the LMM analysis including age as fixed effect. No significant differences between visits were obtained by the LMM analysis in the group with persistent CMA.

In the group with outgrowth of CMA, significantly higher relative abundances of *Coriobacteriaceae* and *Rikenellaceae* proteins, as well as significantly lower relative abundances of *Enterobacteriaceae* proteins were observed at visit 12 months compared to baseline visit (Figure 5C). Moreover, significantly higher relative abundances of *Coriobacteriaceae* and significantly lower relative abundances of *Prevotellaceae* proteins were identified at visit 12 months compared to visit 6 months (Figure 5D).

The results of the LEfSe analysis for the group who outgrew their CMA were all among the top 5 features from LMM analysis ordered by unadjusted p-value (Table S14).

The LEfSe analysis comparing visits within the group with persistent CMA identified significantly higher relative abundances of *Clostridiaceae* and *Veillonellaceae* proteins at visit 6 months compared to baseline (Figure 5E). Significantly higher relative abundances of *Coriobacteriaceae* and significantly lower relative abundances of *Enterococcaceae* and *Enterobacteriaceae* proteins were observed at visit 12 months compared to the baseline visit (Figure 5F). Moreover, significantly higher relative abundances of *Coriobacteriaceae* and significantly lower relative abundances of *Eggerthellaceae* and *Veillonellaceae* proteins were observed at visit 12 months compared to visit 6 months (Figure 5G).

The results of the LEfSe analysis for the group who did not outgrew their CMA were all among the top 10 features from LMM analysis ordered by unadjusted p-value (Table S15).

### Significant differences between allergy groups (outgrowth versus no outgrowth of CMA) in relative abundance levels of microbial protein functional classes

With the exception of the models for pentose and glucoronate interconversions, better goodness of fit (lower AIC) was obtained when not including age as fixed effect in the linear mixed models (LMM)(Table S16).

#### Significant differences of microbial protein functional classes between allergy groups within each visit

The results of LMM analysis, applied to the top 10 microbial protein functional classes, showed a significantly lower relative abundance of glycolysis / gluconeogenesis and pentose and glucuronate interconversions proteins, and a significantly higher relative abundance of RNA degradation proteins in the group with outgrowth of CMA compared to the group with persistent CMA at baseline visit (Table S17). When including age as fixed effect, only glycolysis / gluconeogenesis showed still significant differences between the two allergy groups, while the other effects were only marginally significant (adjusted p-value lower than 0.1). Lower relative abundance of pentose and glucuronate interconversions proteins in the group who outgrew their CMA than in the other group were also found by LEfSe (Figure 6A). Moreover, the results of LEfSe also showed several other protein functional classes with lower relative abundance in the group who outgrew their CMA: Nitrogen metabolism; cysteine and methionine metabolism; valine, leucine and isoleucine degradation, inositol phosphate metabolism, pyrimidine metabolism, selenocompound metabolism and streptomycin biosynthesis. Relative abundances of pyrimidine metabolism were also lower in the group who outgrew their CMA at visit 6 months (Figure 6B). Moreover, at 6 months, the group who outgrew their CMA showed higher relative abundances of proteins from amino sugar and nucleotide sugar metabolism, and from glycine, serine and threonine metabolism in the group who outgrew their CMA than in the other group (Figure 6B). At visit 12 months, relative abundances of proteins from starch and sucrose metabolism were lower in the group who outgrew their CMA than in the other group (Figure 6C).

**Figure 6.**
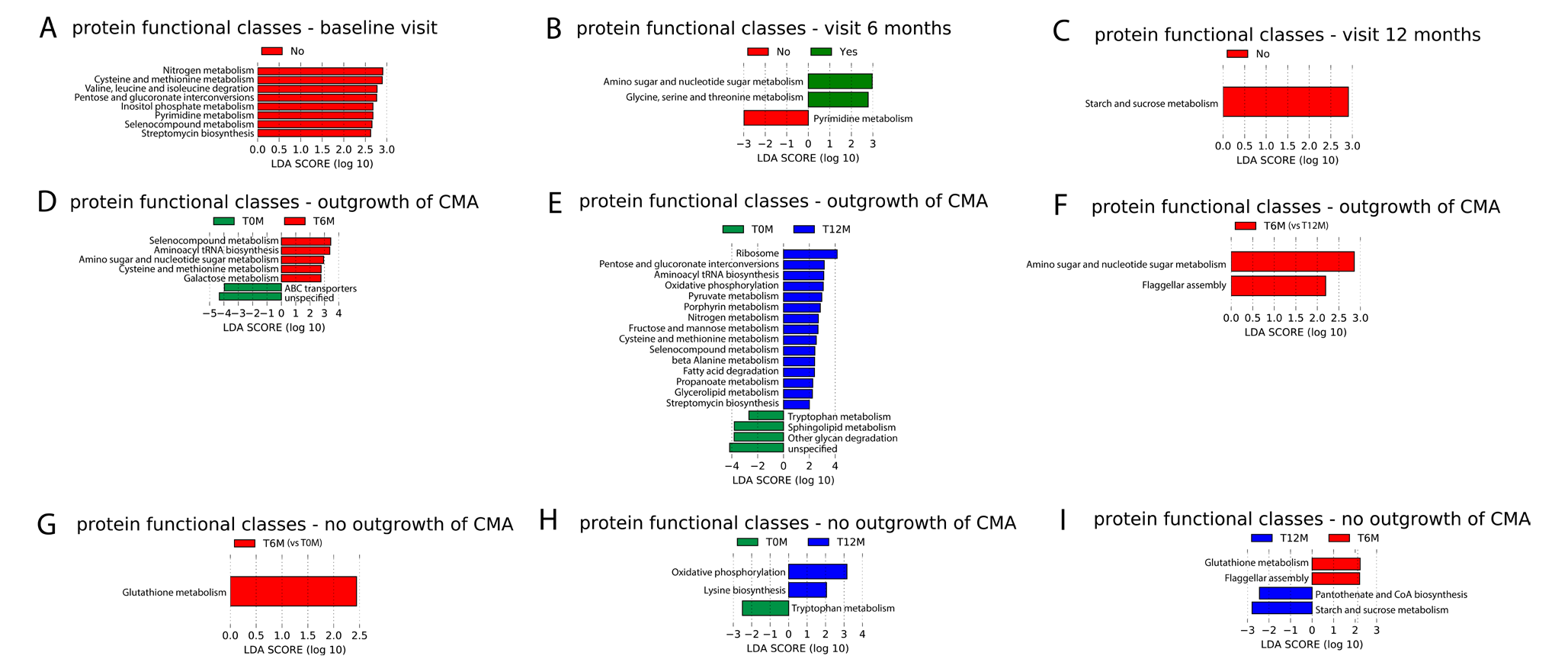
LEfSe analysis of microbial protein functional classes (KEGG Brite level c) using an alpha value of 0.05 for the factorial Kruskal-Wallis test among classes and a threshold of 2.0 on the logarithmic LDA score for discriminative features. (A-C) Using outgrowth of CMA at visit 12 months as class, Yes: outgrowth of CMA at 12 months; no: no outgrowth of CMA at 12 months. Plot of discriminative features at (A) baseline visit; (B) visit 6 months; (C) visit 12 months. (D-F) Pairwise comparison between visits within the group with outgrowth of CMA, using visit as class. (D) visit 6 months (T6M) versus baseline (T0M); (E) visit 12 months (T12M) versus baseline (T0M); (F) visit 12 months (T12M) versus visit 6 months (T6M). (G-I) Pairwise comparison between visits within the group with persistent CMA, using visit as class. (G) visit 6 months (T6M) versus baseline (T0M); (H) visit 12 months (T12M) versus baseline (T0M); (I) visit 12 months (T12M) versus visit 6 months (T6M).

Apart from a few exceptions (nitrogen metabolism at baseline, and cysteine and methionine metabolism at baseline), all results of LEfSe were also reported in the top 10 based on unadjusted p-value from the LMM analysis on all functional classes (Table S18)

#### Significant differences of microbial protein functional classes between visits within allergy groups

In the group with outgrowth of CMA, but not in the group with persistent CMA, significantly higher relative abundances of glycolysis / gluconeogenesis proteins and significantly lower relative abundances of RNA degradation proteins were found at the later visits (6M and 12M) compared to baseline (Table S19). At visit 12 months only, a significantly higher relative abundance of pentose and glucuronate interconvertions proteins was found compared to baseline in the group with outgrowth of CMA, but not in the group with persistent CMA (Table S19). After including age as fixed effect, significant differences were only observed between baseline and 12 months for glycolysis / gluconeogenesis and pentose and glucuronate interconversions. The result for pentose and glucuronate interconversions was also confirmed by LEfSe.

The results of LEfSe showed several other protein functional classes that changed over visit in the group who outgrew their CMA, but not in the other group. Relative abundance of the following protein functional classes were significantly higher at visits 6 months and 12 months than at baseline: Selenocompound metabolism, aminoacyl tRNA biosynthesis and cysteine and methionine metabolism were higher at visits 6 months and 12 months than at baseline (Figures 6D and E). Furthermore, amino sugar and nucleotide sugar metabolism, as well as galactose metabolism increased between baseline and 6 months in this group, while ABC transporters decreased (Figure 6D). Amino sugar and nucleotide sugar metabolism also increased between 6 months and 12 months (Figure 6F). Moreover, several protein functional classes increased between baseline and 12 months (Figure 6E): ribosome, pyruvate metabolism, posphyrin metabolism, nitrogen metabolism, fructose and mannose metabolism, selenocompound metabolism, beta alanine metabolism, fatty acid degradation, propanoate metabolism, glycerolipid metabolism and streptomycin biosynthesis. For sphingolipid metabolism and other glycan degradation, a decrease between baseline and 12 months was observed (Figure 6E).

For the group who outgrew their CMA, the overlap between the LEfSe and the top 10 based on uncorrected p-value from LMM is only moderate (exception: 12 months versus 6 months, were the two functional classes found by LEfSe are in top 3 from LMM based on uncorrected p-value)(Table S20).

LEfSe results also show several protein functional classes that only change over visits in the group which did not outgrew their CMA: gluthatione metabolism (increase between baseline and 6 months (Figure 6G), decrease between 6 months and 12 months(Figure 6I)), cysteine biosynthesis (increase between baseline and 12 months (Figure 6H)), pantenoate CoA biosynthesis (increase between 6 months and 12 months (Figure 6I)), starch and sucrose metabolism (increase between 6 months and 12 months (Figure 6I)).

For the group who did not outgrew their CMA, all protein functional classes found by LEfSe, except one (oxidative phosphorylation) are in the top 10 of LMM based on unadjusted p-value (Table S21).

Finally, also protein classes that changed between visits in both allergy groups were found: oxidative phosphorylation (increase between baseline and 12 months (Figure 6E and H)), tryptophan metabolism (decrease between baseline and 12 months (Figure 6E and H)) and flagellar assembly (increase between 6 months and 12 months (Figure 6F and I)).

### Significant differences between allergy groups (outgrowth versus no outgrowth of CMA) in relative abundance levels of human protein classes

#### Significant differences of human protein classes between allergy groups within each visit

To determine if human protein classes could explain differences between allergy groups, Linear Mixed Model (LMM) analysis was applied. For all human protein classes, the models without age as fixed effect show better goodness of fit (lower AIC)(Table S22). The results show no significant differences between allergy groups within visits (Table S23).

The results of LEfSe analysis showed higher relative abundances of S100 proteins and alkaline phosphatases at baseline in the group who outgrew their CMA compared to the other group (Figure 7A). In the group who outgrew their CMA, lower relative abundances of the actin family and exosomal proteins were found at visit 6 months (Figure 7B). Furthermore, at visit 12 months, higher relative abundances of serpins and lower relative abundances of P-ATPases were found in the group who outgrew their CMA (Figure 7C).

**Figure 7.**
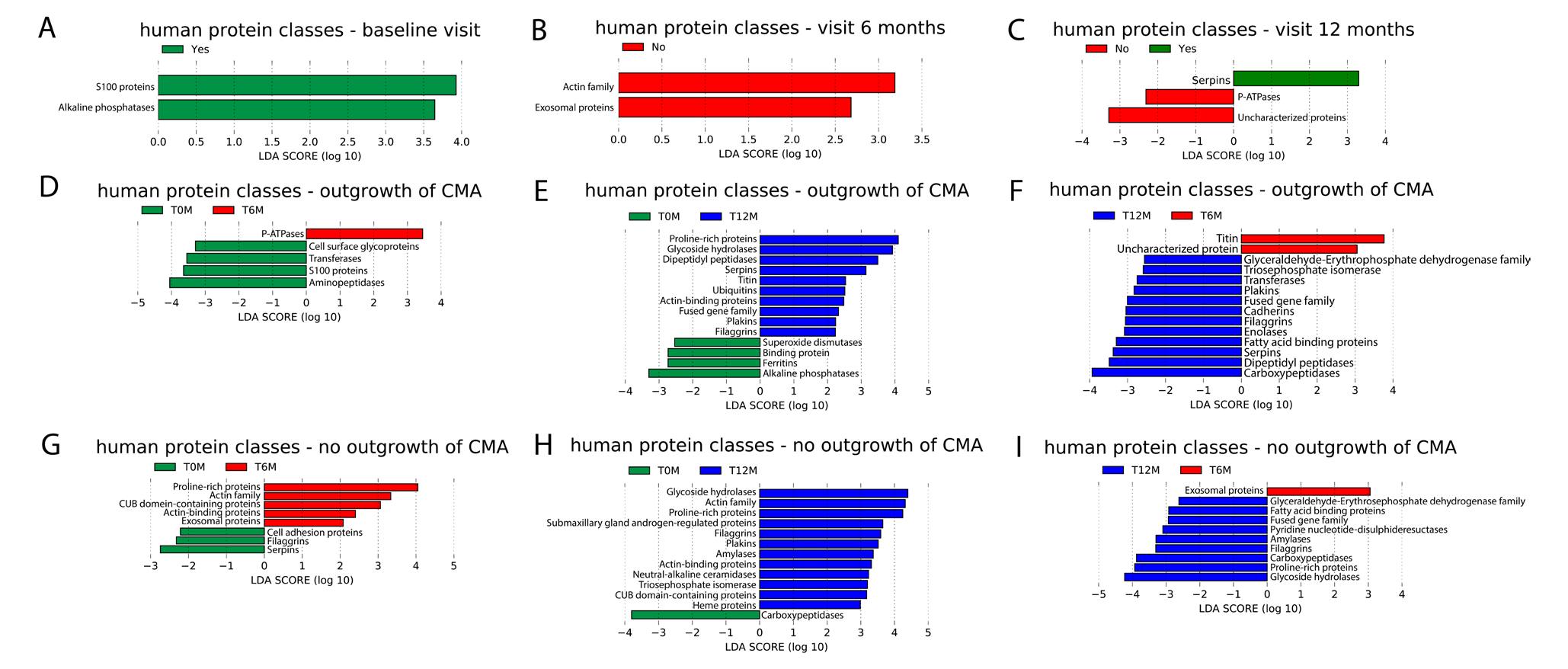
LEfSe analysis of human protein classes using an alpha value of 0.05 for the factorial Kruskal-Wallis test among classes and a threshold of 2.0 on the logarithmic LDA score for discriminative features. (A-C) Using outgrowth of CMA at visit 12 months as class, Yes: outgrowth of CMA at 12 months; no: no outgrowth of at 12 months. Plot of discriminative features at (A) baseline visit; (B) visit 6 months; (C) visit 12 months. (D-F) Pairwise comparison between visits within the group with outgrowth of CMA, using visit as class. (D) visit 6 months (T6M) versus baseline (T0M); (E) visit 12 months (T12M) versus baseline (T0M); (F) visit 12 months (T12M) versus visit 6 months (T6M). (G-I) Pairwise comparison between visits within the group with persistent CMA, using visit as class. (G) visit 6 months (T6M) versus baseline (T0M); (H) visit 12 months (T12M) versus baseline (T0M); (I) visit 12 months (T12M) versus visit 6 months (T6M).

The results of the LEfSe analysis were not significant for the LMM analysis on all human proteins, but all belonged to the top 10 based on unadjusted p-value (Table S24).

#### Significant differences of human protein classes between visits within allergy groups

According to the LMM results on the top 10 human protein classes, relative abundances of proline-rich proteins (4 protein groups) are significantly higher at visit 12 months compared to the baseline visit in the group with outgrowth of CMA, but not in the group with persistent CMA (Table S25). In the group with outgrowth of CMA only, the relative abundance of S100 proteins (4 protein groups) are significantly lower at visit 6 months than at baseline (Table S25). Moreover, relative abundances of immunoglobulins (37 protein groups) are significantly lower at visit 12 months than baseline in infants with persistent CMA, but not in infants who outgrew their CMA (Table S25). These significant differences between visits were not observed when including age as fixed effect.

LEfSe results show higher relative abundances of S100 proteins and alkaline phosphatases in the group who outgrew their CMA at baseline than in the other group (Figure 7A). At visit 6 months, lower relative abundances of exosomal proteins and proteins from the actin family were obtained for the group who outgrew their CMA compared to the other group (Figure 7B). At visit 12 months, the group who outgrew their CMA had higher serpins and lower P-ATpases than the other group (Figure 7C).

Results of LEfSe analysis showed human protein classes that changed over visit in the group who outgrew their CMA, but not in the other group: P-ATPases (increase between baseline and 6 months, Figure 7D); cell surface glycoproteins, S100 proteins and aminopeptidases, transferases (decrease between baseline and 6 months, Figure 7D); dipeptidyl peptidases and serpins (increase between baseline and 12 months and between 6 months and 12 months, Figures 7E and F); titin (increase between baseline and 12 months and decrease between 6 months and 12 months, Figures 7E and F); ubiquitins and fused gene family (increase between baseline and 12 months, Figure 7E); carboxypeptidases, superoxide dismutases, binding proteins, ferritins and alkaline phosphatases (decrease between 6 months and 12 months, Figure 7E); triose phosphate isomerases, plakins, cadherins and enolases (increase between 6 months and 12 months, Figure 7E).

For 6 months versus baseline, and 12 months versus baseline, the majority of the LEfSe results are in the top 10 from the LMM analysis based on unadjusted p-value (Table S26). For 12 months versus 6 months, there is only a moderate overlap between LEfSe resuls and the top 10 list from LMM based on unadjusted p-value (Table S26).

LEfSe results also show human protein classes that only change over visits in the group which did not outgrow their CMA: actin family, CUB domain containing proteins (increase between baseline and 6 months and between baseline and 12 months, Figure 7G and H); exosomal proteins (increase between baseline and 6 months, decrease between 6 months and 12 months, Figure 7G and I); triose phosphate isomerases, cell adhesion proteins and actin-binding proteins (increase between baseline and 6 months, Figure 7G); submaxillary gland androgen-regulated proteins, neutral alkaline-ceramides and heme proteins (increase between baseline and 12 months, Figure 7H); amylases (increase between baseline and 12 months and between 6 months and 12 months, Figure 7H and I); pyridine nucleotide disulphate reductases (increase between 6 months and 12 months, Figure 7I); proline-rich proteins (increase between baseline and 6 months and between 6 months and 12 months, Figures 7 G and I); serpins and filaggrins (decrease between baseline and 6 months (Figure 7G).

For the comparison 6 months versus baseline, the results for LEfSe in the group who did not outgrow their CMA showed large overlap with the top 10 LMM results ordered by unadjusted p-values (Table S27). However for the other comparisons, the overlap was only moderate (Table S27).

Finally, also human protein classes that changed between visits in both allergy groups were found:plakins, proline-rich proteins, glycoside hydrolases and actin-binding proteins (increase between baseline and 12 months, see Figures 7B and H); fatty acid binding proteins, carboxypeptidases, fused gene family and glyceraldehyde-erythrosephosphate dehydrogenase family (increase between 6 months and 12 months, see Figures 7C and I); filaggrins (increase between baseline and 12 months and between 6 months and 12 months, Figures 7B, C, H, I).

### Correlation between human and microbial proteins

To determine correlation between human and microbial proteins, Spearman correlations between human proteins of the top 10 protein classes and microbial proteins of the core taxa were calculated. Several significant correlations, of which the majority are weak, were found (Table S28). Significant moderate correlations (> 0.4) were observed between carboxypeptidases and *Coriobacteriaceae* (correlation =0.4598, p-value < 1e-04), and Transthyretin / hydroxyisourate hydrolases and *Coriobacteriaceae* (correlation =0.4603, p-value < 1e-04).

### Outgrowth of CMA has a low but significant contribution to the variation in 16S-rRNA gene based microbial signatures at visit 6 months

To determine the contribution of environmental and clinical factors to the variation in the gut microbiota, microbial proteome and human proteins, redundancy analysis (RDA) was performed. Outgrowth of CMA significantly explains variation in 16S rRNA gene based taxonomic profiles at visit 6 months (Table S29). However, compared to other significant features, the contribution of outgrowth of CMA is low (shorter edge in RDA plot, see Figure S6). Therefore, ordination plots coloured by outgrowth of CMA do not show a clear separation between the subjects with outgrowth of CMA and those with persistent CMA.

To determine the contribution of outgrowth of CMA to the variation in the gut microbiota and microbial proteome after adjusting for the other environmental variables, partial RDA was performed. The results show that the contribution is low and only significant for the 16S rRNA gene-based taxonomy (% variance explained by outgrowth of CMA = 6.37%, adjusted p-value = 0.043). The partial RDA plot shows that after removing the effect of other environmental factors, a more clear separation between the allergy classes (outgrowth vs persistent CMA) could be observed (Figure S7).

For the other visits, the microbial proteome profiles, the protein-based taxonomic profiles, the microbial protein functional profiles and the human protein profiles, the contribution of outgrowth of CMA to the variation in the data was not significant (Tables S29 and S30).

### Contribution of human proteins on variation in the microbial proteome

To determine human proteins that significantly explain variation in microbial proteome and protein-based taxonomy profiles, another RDA analysis was performed. Table S31 shows that a considerable fraction of the variance in the microbial proteome and protein-based taxonomy profiles can be explained by human proteins at all visits. No clear grouping of the samples by allergy group was observed in any of the RDA plots (Figure S8).

## Discussion

In this study, 16S rRNA gene amplicon sequencing and metaproteomics were applied on faecal samples in a cohort of infants with CMA at baseline visit, of which some showed outgrowth of CMA after 12 months and others did not. The aim was to get insight in outgrowth of CMA.

Apart from analysing the microbiome, environmental and clinical factors associated with outgrowth of CMA were determined. Our results show that among infants with that did not outgrow their CMA, a significantly higher proportion of infants with parental allergy was observed compared to infants that show outgrowth of CMA. This is consistent with previous findings of Xinias et al., who showed that the risk of not acquiring outgrowth of CMA at 1 year of age significantly increased in infants with a family history of atopy ^43^. Furthermore, a significantly lower SCORAD at baseline was observed in infants that show outgrowth of CMA compared to infants that do not. However, the SCORAD of the majority of the infants in this study is lower than 25, corresponding to mild severity of atopic dermatitis ^44^. This suggests that in this cohort, CMA does not manifest mainly through atopic symptoms.

The faecal metaproteome was dominated by bacterial proteins from *Bifidobacteriaceae* and *Lachnospiraceae*. Proteins for these two families were also observed in a metaproteomics study of Kingkaw et al. ^17^ on atopic dermatitis, but the fraction of these taxa was lower than in our study. There are two possible explanations for the differences between our study and Kingkaw et al. First, the infants in our study have another CMA phenotype than atopic dermatitis. Second, the protein identification and quantification of Kingkaw et al. was based on a protein database derived from earlier studies of the gut microbiome which included only 10 bacterial families. In our study, we constructed an in-house proteomics database based on the 16S rRNA gene amplicon sequencing data consisting of 19 bacterial families. Third, the cohort in the study of Kingkaw et al. included both healthy and allergic children, while our study includes only children with CMA at baseline. The gut microbiome of infants with food allergy in their first year of life is known to be dominated by *Lachnospiraceae*, while this family is less abundant in the healthy gut microbiome ^45^.

Six common taxa were found in the top 10 of both 16S rRNA gene sequencing and metaproteomics: *Bacteroidaceae*, *Bifidobacteriaceae*, *Enterobacteriaceae*, *Erysipelatoclostridiaceae*, *Lachnospiraceae* and *Ruminococcaceae*. With the exception of *Erysipelatoclostridiaceae*, these are all common constituents of the healthy gut microbiome ^46, 47^. *Clostridiaceae*, *Coriobacteriaceae*, *Enterococcaceae* and *Tannerellaceae* were only among the top 10 taxa of protein-based microbial taxonomic profiles. The first two are common taxa in the gut microbiome ^46^. The presence of increased levels of *Tannerellaceae* in the gut microbiome has been associated with the administration of synbiotics ^48^. However, protein-based relative abundances of *Tannerellaceae* in the infants that received the synbiotic were lower than in the other infants (synbiotics: median = 0.003, interquartile range: 0.001-0.006; no synbiotics: median = 0.005, interquartile range: 0.003-0.011). This means that its appearance in the top 10 taxa in this study cannot be explained by the inclusion of 23 infants (59% of the infants) which received a formula with synbiotics. Taxa that were only among the top 10 in 16S rRNA gene-based profiles were *Akkermansiaceae*, *Erysipelotrichaceae*, *Streptococcaceae* and *Veillonellaceae*. The latter three are common constituents of the gut microbiome ^46^, while increased levels of *Akkermansiaceae* have been related with synbiotics interventions ^48^. However, in this study, 16S rRNA gene-based relative abundances of *Akkermansiaceae* were lower in the infants that received the synbiotic than in those that did not (synbiotics: median = 0.00019, interquartile range: 0.00005-0.05522; no synbiotics: median = 0.00266, interquartile range: 0.00016-0.60490), so the occurrence of this family in the top 10 is not due to the synbiotic treatment of 23 infants. Differences between 16S rRNA gene-based and protein-based microbial composition have been observed in a previous study ^49^, for which two possible explanations were given by the authors. First, it could be that the 16S rRNA gene-based microbial composition is not representative for microbial protein abundance. Second, we measure a lot less proteins than DNA sequences. Third, protein expression of bacteria can be different. Some bacteria will be more active in producing proteins. Forth, in the pool of proteins that we extracted, we are also depending on the stability of the proteins.

Linear mixed model (LMM) analysis of core taxa and LEfSe analysis on the whole data were applied on the 16S rRNA gene sequencing and metaproteomics data to compare infants who outgrew their CMA with infants that did not.

Our study showed significantly higher 16S rRNA gene-based relative abundances of *Lachnospiraceae* at baseline in infants that did not outgrow their CMA compared to infants who outgrew their CMA. In contrast, a previous study reported higher levels of *Lachnospira pectinoschiza* in infants that outgrew their allergy ^50^. However, this study focuses on children with a mean age of 57.8 months at baseline, while the infants in our study were 15–25 months old at visit 12 months (3-13 months old at baseline). Moreover, higher relative abundance of a family does not mean that the relative abundance of each specific species belonging to that family had to be higher. Furthermore, two families that, to the best of our knowledge, have not been related to CMA (*Carnobacteriaceae* and *Peptostreptococcales-Tissierellales_fa*) showed higher relative abundances at baseline in the group that did not outgrew their CMA.

In infants who outgrew their CMA, higher relative abundances of *Enterococcaceae and Leuconostocaceae* at baseline visit were observed. Higher levels of the genus *Enterococcus* have been reported in non-allergic infants when compared to allergic infants ^51^. In our study, the genus Enterococcus has higher relative abundances at baseline in the group that outgrew their CMA compared to the other group (outgrowth of CMA: median = 0.00142, IQR: 0.00029-0.00544; no outgrowth of CMA: median = 0.00006, IQR: 0.00000-0.00310).

The higher relative abundances of *Aerococcaceae* and *Pasteurellaceae* at visit 6 months in the group who did not outgrew their CMA are in contradiction with Kourosh et al.^52^, who observed lower Pasteurellales in the allergic group compared to the control. However, this study was not restricted to early life but ranged from birth to the age of 18 years old.

Infants who outgrew their CMA showed higher relative abundances of *Eubacteriaceae* at visit 6 months than those who did not. Bunyavanich et al.^53^ reported an association between milk allergy resolution and higher levels of the genus *Eubacterium*. In our study, higher relative abundances of *Eubacterium* were also found at visit 6 months in the infants who outgrew their CMA compared to the other group of infants (outgrowth of CMA: median = 0.00083, IQR: 0.00000-0.00229; no outgrowth of CMA: median = 0.00000, IQR: 0.00000-0.00027).

At visit 12 months, higher *Bacteroidaceae* were observed in infants who did not outgrew their CMA. Higher levels of the phylum Bacteroidetes were reported earlier in infants with persistent CMA ^53^. For the group who outgrew their of CMA, higher relative abundances were found at visit 12 months for three families that, to the best of our knowledge, have not been associated earlier with outgrowth of CMA: *Rikenellaceae*, *Actinomycetaceae* and *Eggerthellaceae*.

When comparing visits within allergy groups, we found higher *Ruminococcaceae* at 12 months compared to baseline in the group with outgrowth of CMA, while no significant changes in this family were observed in infants with persistent CMA. Bunyavanich et al. ^53^ reported increased levels of the genus *Ruminococcus* in infants with resolved CMA compared with persistent CMA. In our study, the median relative abundance increased from 0.00000 to 0.00036 between baseline and 12 months in the group who outgrew their CMA, while in the other group the medians at baseline and 12 months were equal (both equal to 0.00003).

Furthermore, our analyses identified higher *Bacteroidaceae* at 12 months compared to baseline in the group who did not outgrow their CMA, while no significant changes were found in the group who outgrew their CMA. Higher levels of Bacteroidetes were associated earlier with persistent CMA ^53^.

In both allergy groups, lower *Enterobacteriaceae* were observed at 12 months compared to baseline visit. As *Enterobacteriaceae* are generally more abundant in young infants and decrease with age^2^, this finding could be the result of that. Decreased levels of *Enterobacteriaceae* have been previously reported in infants with CMA that received an amino acid-based formula during a period of 6 months ^15^.

In both allergy groups, the results showed higher relative abundances of the families *Butyricicoccaceae* and *Monoglobaceae* at visit 12 months compared to baseline, which suggests that these families change when infants growing older.

In our study, relative abundances of *Eubacteriaceae*, *Oscillospiraceae* and *Tannerellaceae* increased between baseline and 6 months in the group who outgrew their CMA, but not in the other group. To the best of our knowledge, these three families have not been associated with milk allergy resolution earlier.

Furthermore, we found lower *Enterococcaceae*, *Staphylococcaceae* and *Saccharimonadeceae* at visit 6 months compared to baseline in the group who outgrew their CMA, but not in the other group. Significantly lower *Enterococcaceae* were reported earlier in healthy infants compared to infants with food sensitization ^54^. A systematic review ^55^ reported contradictory results in the literature for *Staphylococcaceae* in relation to food sensitization (i.e. some studies relate food sensitization with higher *Staphylococcaceae*, while others found lower *Staphylococcaceae*).

Several families only increase between baseline and 12 months in the group who outgrew their CMA, but not in the other group: *Christensenellaceae*, *Oscillospiraceae*, *Oscillospirales_fa*, *Clostridia UCG 014* family, *Acidaminococcaceae*, *Rikenellaceae* and *Lachnospiraceae*. Higher relative abundances of *Rikenellaceae* were also reported in healthy infants compared to infants with food sensitization by Chen et al. ^54^. The same study also reported higher levels of *Acidaminococcaceae*. However, the results were not significant.

A decrease in *Micrococcaceae* between baseline and visit 12 months was found in the group who outgrew their CMA, but not in the other group. To the best of our knowledge, this family has not been previously related to CMA resolution.

In infants who outgrew their CMA, but not in the other group, higher relative abundances of *Rikenellaceae*, *Actinomycetaceae* and *Prevotellaceae* at visit 12 months compared to visit 6 months were identified. Higher *Prevotellaceae* were related earlier to CMA resolution ^53^.

Remarkably, when comparing visit 12 months to baseline, higher *Prevotellaceae* were found in the group who did not outgrew their CMA, which contradicts with previous findings ^53^.

In the group who did not outgrew their CMA, but not in the other group, higher *Leuconostocaceae* at 12 months compared to baseline, and higher *Selemonadaceae* at 12 months compared to 6 months were observed. As far as we are aware, these two families have not been related to persistent CMA earlier.

A much lower amount of significant results were identified by the metaproteomics analysis at the taxonomy level compared to the analysis of the 16S rRNA gene-based taxonomy. This suggests that changes related to outgrowth of allergy mainly occur at the level of the 16S rRNA gene-based taxonomic classification and less at the protein-based taxonomic classification level. When comparing allergy groups within visits, the only significant result at the family level were higher *Eggerthellaceae* proteins at baseline and higher *Veillonellaceae* proteins at visit 6 months in infants who did not outgrow their CMA compared to the group who outgrew their CMA. To the best of our knowledge, these results have not been reported earlier.

Similar as for 16S rRNA gene sequencing, the results for metaproteomics showed an decrease in relative abundance of *Enterobacteriaceae* proteins between baseline and 12 months in both allergy groups.

The results of the metaproteomics analysis reveal microbial metabolic processes which are expected to play a crucial role in the mechanisms underlying outgrowth of CMA. For most of the processes, multiple bacterial families contribute to their increase over visits in the group who outgrew their CMA (Table S32). In infants who outgrew their CMA, increased relative abundances of *Rikenellaceae* and *Coriobacteriaceae* proteins between baseline and 12 months can be related to an increase in relative abundance of proteins involved in pyruvate metabolism and fructose and mannose metabolism respectively (Table S32).

In general, the analysis on microbial protein functional classes did identify fewer significant differences between allergy groups and visits compared to the analysis of the 16S rRNA gene-based taxonomy, suggesting that changes related to outgrowth of CMA occur at the 16S rRNA gene-based taxonomic level and less at the protein functional level.

In this study, we also determined differences in human protein classes between allergy groups and visits. For children who did not outgrew their at 12 months, significantly lower levels of human immunoglobulins were observed at visit 12 months compared to the baseline visit. Furthermore, higher relative abundances of proline-rich proteins at 12 months compared to baseline, and lower relative abundances of S100 proteins at 6 months compared to baseline were observed in the group who outgrew their allergy, but not in the other group.

The results of the RDA analysis confirm that there is a small contribution of the microbiome to outgrowth of CMA, and that this contribution mainly occurs at the 16S rRNA gene-based taxonomy level.

Our study has several limitations. First, the study has a wide age range at baseline visit (3- 13 months). Given the large changes that the infant gut microbiome undergoes between 3 and 13 months, among others related to weaning, this wide range could affect the results.

Large differences in baseline microbiota and in its development over time are expected between infants of 3 and 13 months old. Moreover, infants of similar age may be included at baseline and 6 months, and at 6 and 12 months. Age-related variability within visits may have confounded the comparisons between visits, potentially obscuring changes related to age and outgrowth of CMA. Overall, adding age as fixed effect to the LMM resulted in worse goodness of fit (higher AIC) and a lower number of significant differences, in particular between visits. Second, the size of the cohort is small, and as a consequence the results have to be interpreted with caution. Validating the results in a similar but larger cohort is a direction for future research. Third, the small size of the cohort made it difficult to detect significant features from a complete list of taxa or protein classes using methods correcting for multiple testing, like LMM analysis. Methods based on effect size rather than statistical significance after multiple testing correction, like LEfSe, are more sensitive to detect discriminative features in a small cohort. Their drawback is the risk for detecting false positives. Therefore, we conducted a LMM analysis on the complete list of families (in addition to the analysis of the core taxa) and the complete lists of microbial functional and human protein classes (in addition of the analysis of the top 10). As expected, the number of significant features found by this LMM analysis was low, but we noticed that the majority of the results found by LEfSe were in the top ranked list of LMM-based unadjusted p-values. This indicates concordance between the LMM and LEfSe results, and suggests that the majority of differential features obtained by LEfSe could have also been detected by LMM in case of availability of a larger cohort.

In summary, we can conclude microbiome differences related to outgrowth of CMA can be mainly identified at the level of the 16S rRNA gene taxonomic level, and to a lesser extent at the protein-based microbial taxonomy and functional level. The overall contribution of the microbiome to outgrowth of CMA is low.

## Data availability

Clinical data are available from Danone Nutricia research upon reasonable request (contact: Harm Wopereis, Danone Nutricia Research, Utrecht, The Netherlands).

The 16S rRNA-gene sequencing for this study have been deposited in the European Nucleotide Archive (ENA) at EMBL-EBI under accession number PRJEB56782 (https://www.ebi.ac.uk/ena/browser/view/PRJEB56782).

The mass spectrometry proteomics data have been deposited to the ProteomeXchange Consortium via the PRIDE ^56^ partner repository with the dataset identifier PXD037190.

The R code for the statistical calculations made in this study has been deposited to a public repository on GitLab (https://git.wur.nl/afsg-microbiology/publication-supplementary-materials/2022-hendrickx-et-al-earlyfit_presto_allergy_study).

## Supporting information

Supplementary results

## Acknowledgements

Guus Roeselers (Danone Nutricia Research) is gratefully acknowledged for his input in the study design. We thank Heleen de Weerd (Danone Nutricia Research) for pre-processing the 16S rRNA gene amplicon sequencing data.

## Author contributions

DMH performed the statistical analysis and wrote the majority of the manuscript, with input of all co-authors. RA and SB performed the metaproteomics experiments, and wrote the corresponding parts of the manuscript. DMH and SKM developed the proteomics database, SKM wrote the corresponding part of the manuscript. The PRESTO study team generated the clinical data. JML provided the 16S rRNA gene sequencing data and was responsible for the project management. CB designed the study, coordinated the project and participated in all phases of the research and article preparation. All authors contributed to writing the manuscript. All authors have approved the final article.

## Additional information

Supplementary material for this paper is available at XXX.

**PRESTO study team**

**Pantipa Chatchatee^4^, Anna Nowak-Wegrzyn^5, 6^, Lars Lange^7^, Suwat Benjaponpitak^8^, Kok Wee Chong^9^, Pasuree Sangsupawanich^10^, Marleen T. J. van Ampting^3^, Manon M. Oude Nijhuis^3^, Lucien F. Harthoorn^3^, Jane E. Langford^3^, Jan Knol^1, 3^, Karen Knipping^3^, Johan Garssen^3, 11^, Valerie Trendelenburg^12^, Robert Pesek^13^, Carla M. Davis^14^, Antonella Muraro^15^, Mich Erlewyn-Lajeunesse^16^, Adam T. Fox^17^, Louise J. Michaelis^18^, Kirsten Beyer^19^, Lee Noimark^20^, Gary Stiefel^21^, Uwe Schauer^22^, Eckard Hamelmann^22^, Diego Peroni^23^, and Attilio Boner^23^**

^4^ Pediatric Allergy and Clinical Immunology Research Unit, Division of Allergy and Immunology, Department of Pediatrics, Faculty of Medicine, Chulalongkorn University, King Chulalongkorn Memorial Hospital, Thai Red Cross Society, Bangkok, Thailand

^5^ Allergy and Immunology, Department of Pediatrics, New York University Langone Health, New York, US

^6^ Department of Pediatrics, Gastroenterology and Nutrition, Collegium Medicum, University of Warmia and Mazury, Olsztyn, Poland

^7^St Marien Hospital,Bonn, Germany

^8^Pediatric Allergy and Immunology Division, Department of Pediatrics, Faculty of Medicine Ramathibodi Hospital, Mahidol University, Bangkok, Thailand

^9^Allergy Service, Department of Paediatric Medicine, KK Women’s & Children’s Hospital, Singapore

^10^Department of Pediatrics, Faculty of Medicine, Prince of Songkla University, Hat Yai, Thailand

^11^Utrecht Institute for Pharmaceutical Sciences, Utrecht University, Utrecht, The Netherlands

^12^ Department of Pediatric Pneumology, Immunology, and Intensive Care Medicine, Charité Universitätsmedizin Berlin, Berlin, Germany

^13^Arkansas Children’s Hospital, Little Rock, US

^14^Texas Children’s Hospital, Baylor College of Medicine, Houston, US

^15^Food Allergy Referral Centre, Padua University Hospital, Padua, Italy

^16^University Hospitals Southampton, Southampton, United Kingdom

^17^Guy’s and St Thomas’ NHS Foundation Trust, London, United Kingdom

^18^Great North Children’s Hospital, Newcastle Upon Tyne Hospitals NHS Foundation Trust, Newcastle Upon Tyne, United Kingdom

^19^Department of Pediatric Pneumology, Immunology, and Intensive Care Medicine, Charité Universitätsmedizin Berlin, Berlin, Germany

^20^Barts/Royal London Hospital, London, England, United Kingdom

^21^Leicester Royal Infirmary, Leicester, England, United Kingdom ^22^Ruhr-Universitat Bochem im St Josef-Hospital, Bochum, Germany

^23^University Hospital Verona, Verona, Italy

