## Supplementary results for "Assessment of infant outgrowth of cow’s milk allergy in relation to the faecal microbiome and metaproteome"

Table S1. Clinical characteristics. Numeric variables are presented as mean ± standard deviation; categorical variable are presented as number (%). SCORAD: severity of atopic dermatitis. AAF-syn: amino acid-based formula + synbiotics; AAF: standard amino acid-based formula. Vomiting, spitting, stool frequency, stool consistency, stool colour, gas/wind: parent reported outcomes - mean score of 3 reported values. Vomiting: 0 = none; 1 = 1-2 times/day; 2 = 3-4 times/day. Spitting: 0 = none; 1 = spitting up after some feeds; 2 = spitting up after all feeds. Stool frequency: 0 = none; 1 = few; 2 = several; 3 = a lot. Stool consistency: 0 = Severe diarrhoea; 1 = Diarrhoea; 2 = Normal; 3 = Constipation. Stool colour: 0 = green; 1 = yellow; 2 = brown; 3 = Dark brown / Blackish. Gas / wind: 0 = none; 1 = mild; 2 = moderate; 3 = severe.

| **characteristic** | **outgrowth of CMA (n=24)** | **no outgrowth of CMA (n=15)** | **total (n=39)** |
|---|---|---|---|
| sex: female<br>male | 8 (33%)<br>16 (67%) | 3 (20%)<br>12 (80%) | 11 (28%)<br>28 (72%) |
| age: baseline<br>6 months<br>12 months | 8.56 ± 3.04<br>14.62 ± 3.02<br>20.84 ± 3.05 | 9.68 ± 2.63<br>15.59 ± 2.54<br>21.88 ± 3.01 | 9.00 ± 2.90<br>14.99 ± 2.85<br>21.24 ± 3.03 |
| race: Causasian/white<br>Asian<br>Combination of the above/other | 5 (21%)<br>18 (75%)<br>1 (4%) | 4 (27%)<br>10 (67%)<br>1 (7%) | 9 (23%)<br>28 (72%)<br>2 (5%) |
| mode of delivery:<br>vaginal<br>ceasarian | <br>6 (25%)<br>18 (75%) | <br>7 (47%)<br>8 (53%) | <br>13 (33%)<br>26 (67%) |
| allergy mother: yes<br>no | 9 (38%)<br>15 (63%) | 10 (67%)<br>5 (33%) | 19 (49%)<br>20 (51%) |
| allergy father: yes<br>no | 6 (25%)<br>18 (75%) | 9 (60%)<br>6 (40%) | 15 (38%)<br>24 (62%) |
| sibling: yes<br>no | 18 (75%)<br>6 (25%) | 10 (67%)<br>5 (33%) | 28 (72%)<br>11 (28%) |
| SCORAD: baseline<br>6 months<br>12 months | 8.98 ± 14.41<br>5.46 ± 8.32<br>6.77 ± 8.25 | 16.27 ± 13.24<br>8.13 ± 9.67<br>10.37 ± 8.77 | 11.78 ± 14.25<br>6.49 ± 8.84<br>8.15 ± 8.52 |
| egg allergy: yes<br>no | 9 (38%)<br>15 (63%) | 5 (33%)<br>10 (67%) | 14 (36%)<br>25 (64%) |
| other food allergy than CMA and egg allergy:<br>yes<br>no | <br>3 (13%)<br>21 (88%) | <br>6 (40%)<br>9 (60%) | <br>9 (23%)<br>30 (77%) |
| treatment: AAF-syn<br>AAF | 14 (58%)<br>10 (42%) | 9 (60%)<br>6 (40%) | 23 (59%)<br>16 (41%) |
| vomiting:<br>6 months<br>12 months | <br>0.13 ± 0.29<br>0.04 ± 0.20 | <br>0.16 ± 0.52<br>0.20 ± 0.43 | <br>0.14 ± 0.39<br>0.10 ± 0.32 |
| spitting:<br>6 months<br>12 months | <br>0.13 ± 0.27<br>0.04 ± 0.15 | <br>0.11 ± 0.43<br>0.00 ± 0.00 | <br>0.12 ± 0.34<br>0.03 ± 0.12 |
| stool frequency:<br>6 months<br>12 months | <br>1.03 ± 0.24<br>1.11 ± 0.60 | <br>1.02 ± 0.96<br>0.48 ± 0.17 | <br>1.03 ± 0.35<br>1.05 ± 0.49 |
| stool consistency:<br>6 months<br>12 months | <br>1.60 ± 0.61<br>1.92 ± 0.62* | <br>1.89 ± 0.84*<br>1.97 ± 0.77* | <br>1.69 ± 0.70*<br>1.94 ± 0.67** |
| stool colour:<br>6 months<br>12 months | <br>1.44 ± 0.61<br>1.30 ± 0.80* | <br>1.06 ± 0.96*<br>1.89 ± 0.78* | <br>1.32 ± 0.92*<br>1.52 ± 0.83** |
| gas / wind:<br>6 months<br>12 months | <br>0.79 ± 0.61<br>0.86 ± 1.61 | <br>0.56 ± 0.63<br>0.56 ± 0.63 | <br>0.70 ± 0.73<br>0.74 ± 0.62 |
| number of antibiotics until visit:<br>6 months<br>12 months | <br>0.63 ± 2.61<br>1.25 ± 3.61 | <br>0.53 ± 1.41<br>2.40 ± 2.64 | <br>0.59 ± 1.21<br>1.69 ± 2.12 |
| number of infections until visit:<br>6 months<br>12 months | <br>1.25 ± 4.61<br>1.96 ± 5.61 | <br>1.67 ± 1.45<br>3.00 ± 2.30 | <br>1.41 ± 1.58<br>2.36 ± 2.31 |

*: 3 missings; **: 6 missings

Table S1. Clinical characteristics (continued)

| **characteristic** | **outgrowth of CMA (n=24)** | **no outgrowth of CMA (n=15)** | **total (n=39)** |
|---|---|---|---|
| study centre: | | | |
| Southampton General Hospital (UK) | 1 (4%) | 0 (0%) | 1 (3%) |
| Charité Hospital Berlin (Germany) | 1 (4%) | 1 (7%) | 2 (5%) |
| St.-Marien Hospital (Germany) | 3 (13%) | 1 (7%) | 4 (10%) |
| Ruhr-Universität Bochum im St. Josef-Hospital (Germany) | 1 (4%) | 0 (0%) | 1 (3%) |
| KK Women's & Children's Hospital (Singapore) | 2 (8%) | 5 (33%) | 7 (18%) |
| King Chulalongkorn Memorial Hospital (Thailand) | 6 (25%) | 4 (27%) | 10 (26%) |
| Ramathibodi Hospital (Thailand) | 5 (21%) | 2 (13%) | 7 (18%) |
| Prince of Songkla Hospital (Thailand) | 4 (17%) | 1 (7%) | 5 (13%) |
| Texas Children's Hospital (USA) | 1 (4%) | 1 (7%) | 2 (5%) |
| CM-specific IgE | | | |
| baseline | 1.70 ± 2.80 | 23.28 ± 50.99 | 10.00 ± 32.80 |
| 12 months | 1.11 ± 1.74 | 16.88 ± 20.91 | 7.18 ± 14.94 |
| total IgE | | | |
| baseline | 144.64 ± 221.27 | 564.78 ± 831.55 | 306.23 ± 572.07 |
| 12 months | 190.87 ± 199.50 | 562.64 ± 536.20 | 333.86 ± 404.46 |

Table S2. List of proteomes used for the proteomics database construction. The last two columns show the SILVA 138 genus and the subtotal of the relative abundances at genus level. For the taxa indicated in red, only the identified proteins (instead of the complete proteomes) were included.

See Supplementary_Table_S2.xlsx

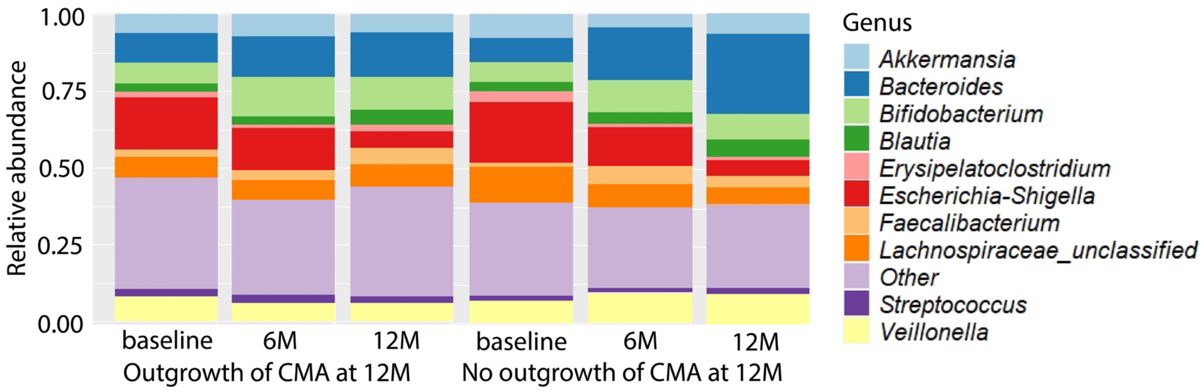

Figure S1. 16S rRNA gene-based taxonomic profiles at the genus level at each visit for the group that outgrew their CMA at visit 12 months (12M) and the group that did not.

Table S3. Clinical factors associated with outgrowth of cow's milk allergy as determined by a two-sided Mann-Whitney U-test for numeric variables and a Fisher's exact test for binary variables. Numeric variables are presented as mean ± standard deviation; categorical variable are presented as number (%). *: 3 missings

| **variable** | **outgrowth of CMA (n=24)** | **no outgrowth of CMA (n=15)** | **p-value** |
|---|---|---|---|
| age at baseline | 8.56 ± 3.04 | 9.68 ± 2.63 | 0.254 |
| allergy of at least one of the parents | yes: 13 (54%)<br>no: 11 (46%) | yes: 13 (87%)<br>no: 2 (13%) | **0.045** |
| mode of delivery | vaginal: 6 (25%)<br>ceasarian: 18 (75%) | vaginal: 7 (47%)<br>ceasarian: 8 (53%) | 0.185 |
| egg allergy | yes: 9 (38%)<br>no: 15 (63%) | yes: 5 (33%)<br>no: 10 (67%) | 1.000 |
| gaswind 6 months | 0.79 ± 0.61 | 0.56 ± 0.63 | 0.504 |
| gaswind 12 months | 0.86 ± 1.61 | 0.56 ± 0.63 | 0.103 |
| number of antibiotics until visit 6 months | 0.63 ± 2.61 | 0.53 ± 1.41 | 0.389 |
| number of antibiotics until visit12 months | 1.25 ± 3.61 | 2.40 ± 2.64 | 0.212 |
| number of infections until visit 6 months | 1.25 ± 4.61 | 1.67 ± 1.45 | 0.176 |
| number of infections until visit 12 months | 1.96 ± 5.61 | 3.00 ± 2.30 | 0.096 |
| other food allergy | yes: 3 (13%)<br>no: 21 (88%) | yes: 6 (40%)<br>no: 9 (60%) | 0.063 |
| SCORAD 0 months | 8.98 ± 14.41 | 16.27 ± 13.24 | **0.036** |
| SCORAD 6 months | 5.46 ± 8.32 | 8.13 ± 9.67 | 0.338 |
| SCORAD 12 months | 6.77 ± 8.25 | 10.37 ± 8.77 | 0.218 |
| sex | female: 8 (33%)<br>male: 16 (67%) | female: 3 (20%)<br>male: 12 (80%) | 0.477 |
| sibling | yes: 18 (75%)<br>no: 6 (25%) | yes: 10 (67%)<br>no: 5 (33%) | 0.718 |
| spitting 6 months | 0.13 ± 0.27 | 0.11 ± 0.43 | 0.441 |
| spitting 12 months | 0.04 ± 0.15 | 0.00 ± 0.00 | 0.274 |
| stool colour 6 months | 1.44 ± 0.61 | 1.06 ± 0.96* | 0.096 |
| stool colour 12 months | 1.30 ± 0.80* | 1.89 ± 0.78* | 0.144 |
| stool consistency 6 months | 1.60 ± 0.61 | 1.89 ± 0.84* | 0.171 |
| stool consistency 12 months | 1.92 ± 0.62* | 1.97 ± 0.77* | 0.825 |
| stool frequency 6 months | 1.03 ± 0.24 | 1.02 ± 0.96 | 0.130 |
| stool frequency 12 months | 1.11 ± 0.60 | 0.48 ± 0.17 | 0.636 |
| treatment (synbiotics vs non-synbiotics) | AAF-syn: 14 (58%)<br>AAF: 10 (42%) | AAF-syn: 9 (60%)<br>AAF: 6 (40%) | 1.000 |
| vomiting 6 months | 0.13 ± 0.29 | 0.16 ± 0.52 | 0.854 |
| vomiting 12 months | 0.04 ± 0.20 | 0.20 ± 0.43 | 0.125 |
| study centre (western vs non-western country) | western: 7 (29%)<br>non-western: 17 (71%) | western: 3 (20%)<br>non-western: 12 (80%) | 0.711 |

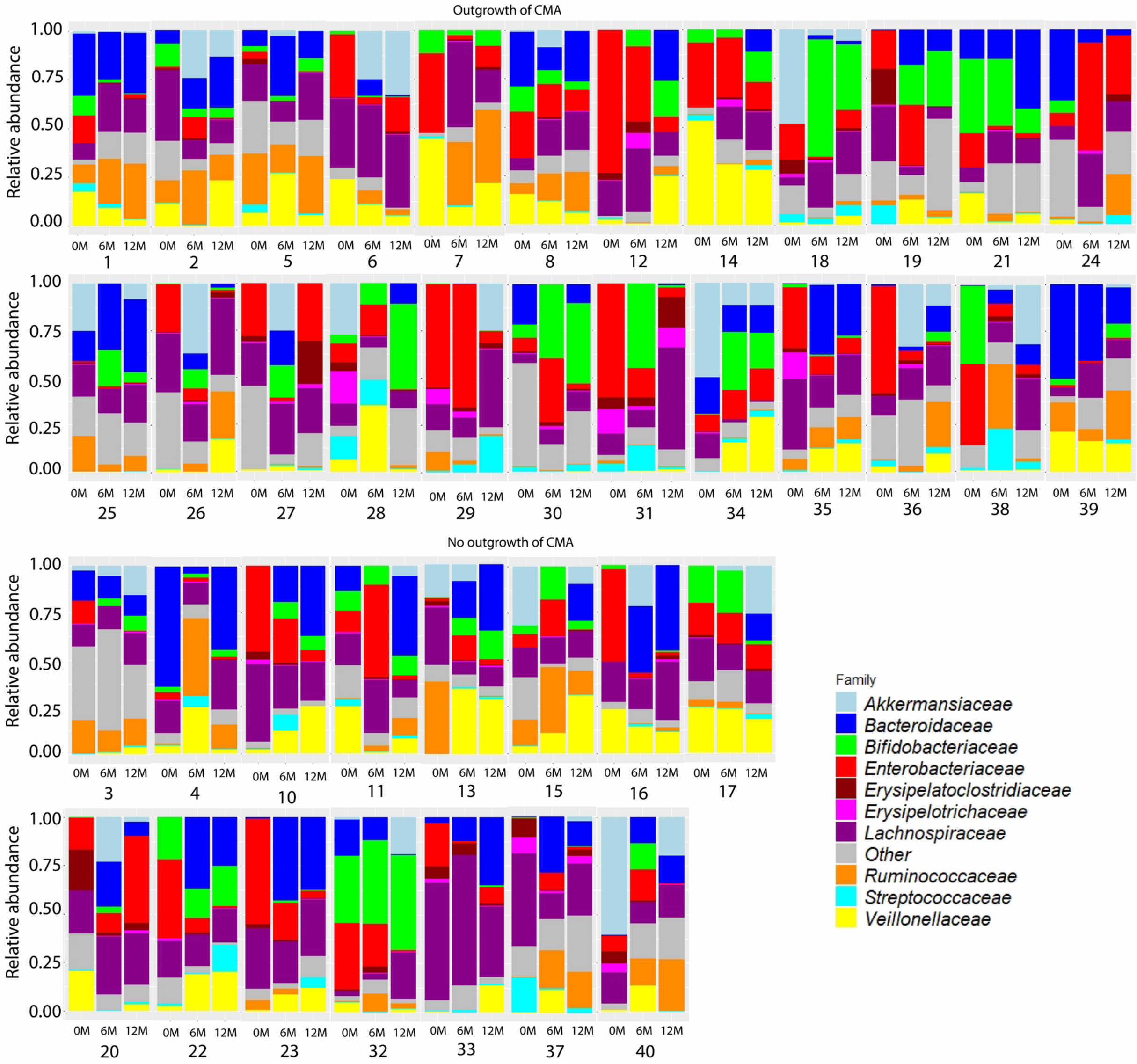

Figure S2. Microbiota composition profiles for each visit (0 months, 6 months, 12 months) for each infant. Subject numbers correspond with Supplementary Table S1. Taxonomy at family level based on 16S rRNA gene sequencing data.

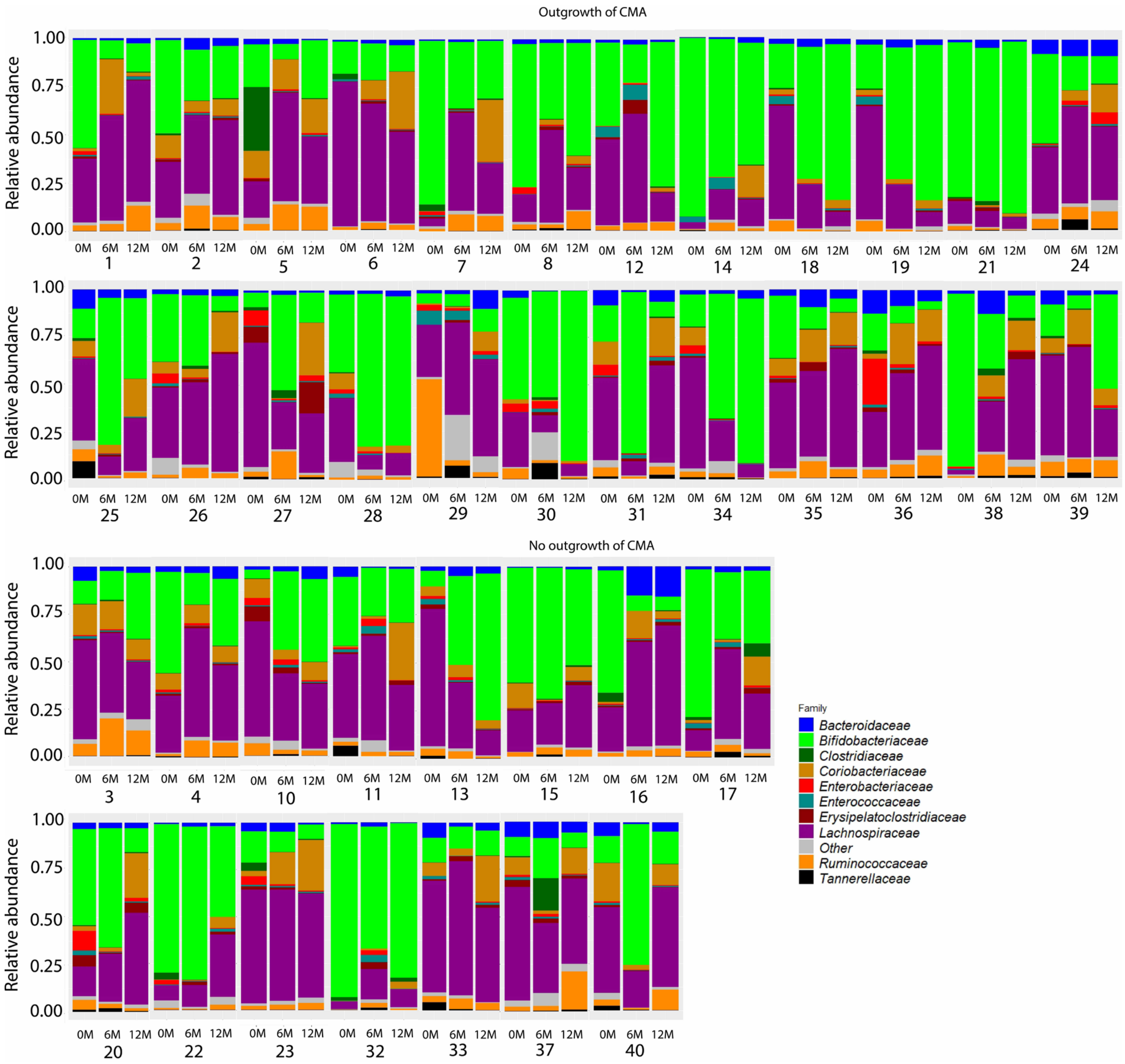

Figure S3. Microbiota composition profiles for each visit (0 months, 6 months, 12 months) for each infant. Subject numbers correspond with Supplementary Table S1. Taxonomy at family level based on metaproteomics data.

Table S4. Akaike Information Criterion (AIC) values for Linear Mixed Models (LMM) fitted to 16S rRNA gene based relative abundance of core families. Model A: model without age as fixed effect, model B: model including age as fixed effect. For each core family, the best model (lowest AIC) is indicated in bold.

| | AIC – model A | AIC – model B |
|---|---|---|
| ***Bifidobacteriaceae*** | **575.2536** | 579.3477 |
| ***Bacteroidaceae*** | **548.7505** | 549.9798 |
| ***Lachnospiraceae*** | **373.4082** | 378.1709 |
| ***Ruminococcaceae*** | **512.6130** | 516.0835 |
| ***Coriobacteriaceae*** | **588.2699** | 592.4549 |
| ***Veillonellaceae*** | **535.6380** | 539.9659 |
| ***Enterobacteriaceae*** | **456.1456** | 457.4884 |

Table S5. Median 16S rRNA gene based relative abundance and interquartile range (IQR) of core families in faeces samples at baseline and follow-up visits (6 months, 12 months) in infants that outgrew their CMA after 12 months (12M) and children that did not. adj. p-value: p-value determined by Linear Mixed Model (LMM) analysis with Benjamini-Hochberg correction for multiple testing. Core taxa were defined as taxa that have a relative abundance higher than 1% in at least 50% of the 16S rRNA gene or metaproteomics samples. A: model without age as fixed effect, B: model including age as fixed effect.

| | **Baseline visit** | | |
|---|---|---|---|
| | **Outgrowth of CMA** | **No outgrowth of CMA** | **adj. p-value** |
| ***Bifidobacteriaceae*** | 0.025 (0.001-0.079) | 0.004 (0.001-0.076) | A: 0.9594; B: 0.9026 |
| ***Bacteroidaceae*** | 0.004 (0.000-0.166) | 0.005 (0.000-0.081) | A: 0.9594; B: 0.9026 |
| ***Lachnospiraceae*** | **0.116 (0.058-0.198)** | **0.212 (0.165-0.306)** | **A: 0.0280; B: 0.0457** |
| ***Ruminococcaceae*** | 0.004 (0.000-0.070) | 0.004 (0.000-0.044) | A: 0.9704; B: 0.9026 |
| ***Coriobacteriaceae*** | 0.00003 (0.00000-0.00406) | 0.00000 (0.00000-0.00022) | A: 0.9594; B: 0.9026 |
| ***Veillonellaceae*** | 0.023 (0.002-0.154) | 0.026 (0.006-0.128) | A: 0.9704; B: 0.9026 |
| ***Enterobacteriaceae*** | 0.218 (0.084-0.350) | 0.165 (0.074-0.373) | A: 0.9594; B: 0.9026 |
| | **Visit 6 months** | | |
| | **Outgrowth of CMA** | **No outgrowth of CMA** | **adj. p-value** |
| ***Bifidobacteriaceae*** | 0.059 (0.011-0.195) | 0.087 (0.018-0.143) | A: 0.7222; B: 0.6813 |
| ***Bacteroidaceae*** | 0.084 (0.003-0.041) | 0.137 (0.078-0.257) | A: 0.6884; B: 0.6813 |
| ***Lachnospiraceae*** | 0.167 (0.101-0.237) | 0.143 (0.118-0.219) | A: 0.6884; B: 0.6813 |
| ***Ruminococcaceae*** | 0.032 (0.004-0.116) | 0.029 (0.002-0.125) | A: 0.8870; B: 0.9970 |
| ***Coriobacteriaceae*** | 0.00003 (0.00000-0.00358) | 0.00000 (0.00000-0.00006) | A: 0.6884; B: 0.6813 |
| ***Veillonellaceae*** | 0.017 (0.003-0.122) | 0.119 (0.010-0.165) | A: 0.6884; B: 0.6813 |
| ***Enterobacteriaceae*** | 0.080 (0.016-0.202) | 0.129 (0.050-0.192) | A: 0.6884; B: 0.6813 |
| | **Visit 12 months** | | |
| | **Outgrowth of CMA** | **No outgrowth of CMA** | **adj. p-value** |
| ***Bifidobacteriaceae*** | 0.052 (0.018-0.143) | 0.037 (0.007-0.093) | A: 0.9059; B: 0.8378 |
| ***Bacteroidaceae*** | 0.114 (0.019-0.258) | 0.252 (0.133-0.375) | A: 0.4883; B: 0.5029 |
| ***Lachnospiraceae*** | 0.194 (0.120-0.242) | 0.206 (0.169-0.265) | A: 0.4883; B: 0.5558 |
| ***Ruminococcaceae*** | 0.040 (0.014-0.217) | 0.022 (0.002-0.131) | A: 0.4883; B: 0.5029 |
| ***Coriobacteriaceae*** | 0.00005 (0.00002-0.02495) | 0.00003 (0.00000-0.00596) | A: 0.4883; B: 0.5558 |
| ***Veillonellaceae*** | 0.051 (0.012-0.159) | 0.114 (0.026-0.191) | A: 0.4883; B: 0.5558 |
| ***Enterobacteriaceae*** | 0.050 (0.012-0.108) | 0.016 (0.008-0.046) | A: 0.6889; B: 0.8378 |

Table S6. Difference in 16S rRNA gene based taxa at family level between allergy groups within visits as determined by Linear Mixed Model (LMM) analysis with outgrowth of CMA, visit and outgrowth of CMA x visit as fixed effects and subject as random effect. Abbreviation: p-adj.: adjusted p-value (p-value corrected for multiple testing using the Benjamini-Hochberg correction). Top 10 features ordered by unadjusted p-value.

| **Baseline visit – outgrowth vs no outgrowth of CMA** | | | |
|---|---|---|---|
| **Family** | **p-value LMM** | **p-adj. LMM** | **Found by LEfSe** |
| *Lachnospiraceae* | 0.004 | 0.250 | yes |
| *Carnobacteriaceae* | 0.025 | 0.487 | yes |
| *Atopobiaceae* | 0.035 | 0.487 | |
| *Enterococcaceae* | 0.037 | 0.487 | yes |
| *Saccharimonadaceae* | 0.040 | 0.487 | |
| *Enterobacterales_unclassified* | 0.046 | 0.487 | |
| *Leuconostocaceae* | 0.058 | 0.526 | yes |
| *Staphylococcaceae* | 0.071 | 0.563 | |
| *Streptococcaceae* | 0.091 | 0.640 | |
| *Clostridia_UCG.014_fa* | 0.128 | 0.806 | |
| **Visit 6 months – outgrowth vs no outgrowth of CMA** | | | |
| **Family** | **p-value LMM** | **p-adj. LMM** | **Found by LEfSe** |
| *Aerococcaceae* | 0.004 | 0.254 | yes |
| *Pasteurellaceae* | 0.008 | 0.257 | yes |
| *Eubacteriaceae* | 0.058 | 0.688 | yes |
| *Prevotellaceae* | 0.063 | 0.688 | |
| *Monoglobaceae* | 0.104 | 0.688 | |
| *Tannerellaceae* | 0.138 | 0.688 | |
| *Oscillospiraceae* | 0.152 | 0.688 | |
| *Oscillospirales_fa* | 0.155 | 0.688 | |
| *Bacteria_unclassified* | 0.163 | 0.688 | |
| *Muribaculaceae* | 0.163 | 0.688 | |
| **Visit 12 months – outgrowth vs no outgrowth of CMA** | | | |
| **Family** | **p-value LMM** | **p-adj. LMM** | **Found by LEfSe** |
| *Rikenellaceae* | 0.005 | 0.213 | yes |
| *Prevotellaceae* | 0.007 | 0.213 | |
| *Comamonadaceae* | 0.022 | 0.453 | |
| *Actinomycetaceae* | 0.036 | 0.564 | yes |
| *Gastranaerophilales_fa* | 0.079 | 0.582 | |
| *Bacteroidaceae* | 0.082 | 0.582 | yes |
| *Bacteria_unclassified* | 0.102 | 0.582 | |
| *Muribaculaceae* | 0.102 | 0.582 | |
| *RF39_fa* | 0.102 | 0.582 | |
| *Defluviitaleaceae* | 0.102 | 0.582 | |

Table S7. Significance of difference in 16S rRNA gene based core taxa between visits within each allergy group as determined by Linear Mixed Model (LMM) analysis with Benjamini-Hochberg correction. Upper part of the table: infants that outgrew their CMA at 12 months; lower part of the table: infants that did not outgrew their CMA at 12 months. Abbreviations: Med.(IQR): median relative abundance and interquartile range; 0M: 0 months (baseline); 6M: 6 months; 12M: 12 months; p-adj.: adjusted p-value (p-value corrected for multiple testing using the Benjamini-Hochberg correction). A: model without age as fixed effect, B: model including age as fixed effect.

| **Infants that outgrew their CMA at visit 12 months** | | | | | | |
|---|---|---|---|---|---|---|
| | **Med.(IQR) 0M** | **Med.(IQR) 6M** | **Med.(IQR) 12M** | **p-adj. 0M vs 6M** | **p-adj. 0M vs 12M** | **p-adj. 6M vs 12M** |
| ***Bifidobacteriaceae*** | 0.025 (0.001-0.079) | 0.059 (0.011-0.195) | 0.052 (0.018-0.143) | A:0.8512 B:0.9911 | A:0.9395 B:0.9882 | A:0.9450 B:0.9943 |
| ***Bacteroidaceae*** | 0.004 (0.000-0.166) | 0.084 (0.003-0.041) | 0.114 (0.019-0.258) | A:0.6587 B:0.9911 | A:0.1561 B:0.9882 | A:0.9146 B:0.9943 |
| ***Lachnospiraceae*** | 0.116 (0.058-0.198) | 0.167 (0.101-0.237) | 0.194 (0.120-0.242) | A:0.6587 B:0.9911 | A:0.6758 B:0.9882 | A:0.9450 B:0.9943 |
| ***Ruminococcaceae*** | 0.004 (0.000-0.070) | 0.032 (0.004-0.116) | 0.040 (0.014-0.217) | A:0.3301 B:0.9911 | **A:0.0182** B:0.9882 | A:0.9146 B:0.9943 |
| ***Coriobacteriaceae*** | 0.00003 (0.00000-0.00406) | 0.00003 (0.00000-0.00358) | 0.00005 (0.00002-0.02495) | A:0.9095 B:0.9911 | A:0.6758 B:0.9882 | A:0.7840 B:0.9943 |
| ***Veillonellaceae*** | 0.023 (0.002-0.154) | 0.017 (0.003-0.122) | 0.051 (0.012-0.159) | A:0.8818 B:0.9911 | A:0.9847 B:0.9882 | A:0.9146 B:0.9943 |
| ***Enterobacteriaceae*** | 0.218 (0.084-0.350) | 0.080 (0.016-0.202) | 0.050 (0.012-0.108) | **A:0.0406** B:0.9911 | **A:0.00003** B:0.9882 | A:0.7840 B:0.9943 |
| **Infants that did not outgrew their CMA at visit 12 months** | | | | | | |
| | **Med.(IQR) 0M** | **Med.(IQR) 6M** | **Med.(IQR) 12M** | **p-adj. 0M vs 6M** | **p-adj. 0M vs 12M** | **p-adj. 6M vs 12M** |
| ***Bifidobacteriaceae*** | 0.004 (0.001-0.076) | 0.087 (0.018-0.143) | 0.037 (0.007-0.093) | A:0.4933 B:0.9414 | A:0.7403 B:0.9965 | A:0.9859 B:0.9997 |
| ***Bacteroidaceae*** | 0.005 (0.000-0.081) | 0.137 (0.078-0.257) | 0.252 (0.133-0.375) | A:0.1638 B:0.9414 | **A:0.0007** B:0.9965 | A:0.8307 B:0.9997 |
| ***Lachnospiraceae*** | 0.212 (0.165-0.306) | 0.143 (0.118-0.219) | 0.206 (0.169-0.265) | A:0.6960 B:0.9414 | A:0.7403 B:0.9965 | A:0.9859 B:0.9997 |
| ***Ruminococcaceae*** | 0.004 (0.000-0.044) | 0.029 (0.002-0.125) | 0.022 (0.002-0.131) | A:0.3861 B:0.9414 | A:0.7403 B:0.9965 | A:0.9859 B:0.9997 |
| ***Coriobacteriaceae*** | 0.00000 (0.00000-0.00022) | 0.00000 (0.00000-0.00006) | 0.00003 (0.00000-0.00596) | A:0.7983 B:0.9414 | A:0.7403 B:0.9965 | A:0.8307 B:0.9997 |
| ***Veillonellaceae*** | 0.026 (0.006-0.128) | 0.119 (0.010-0.165) | 0.114 (0.026-0.191) | A:0.6960 B:0.9414 | A:0.7403 B:0.9965 | A:0.9859 B:0.9997 |
| ***Enterobacteriaceae*** | 0.165 (0.074-0.373) | 0.129 (0.050-0.192) | 0.016 (0.008-0.046) | A:0.6960 B:0.9414 | **A:0.0007** B:0.9965 | A:0.0602 B:0.9997 |

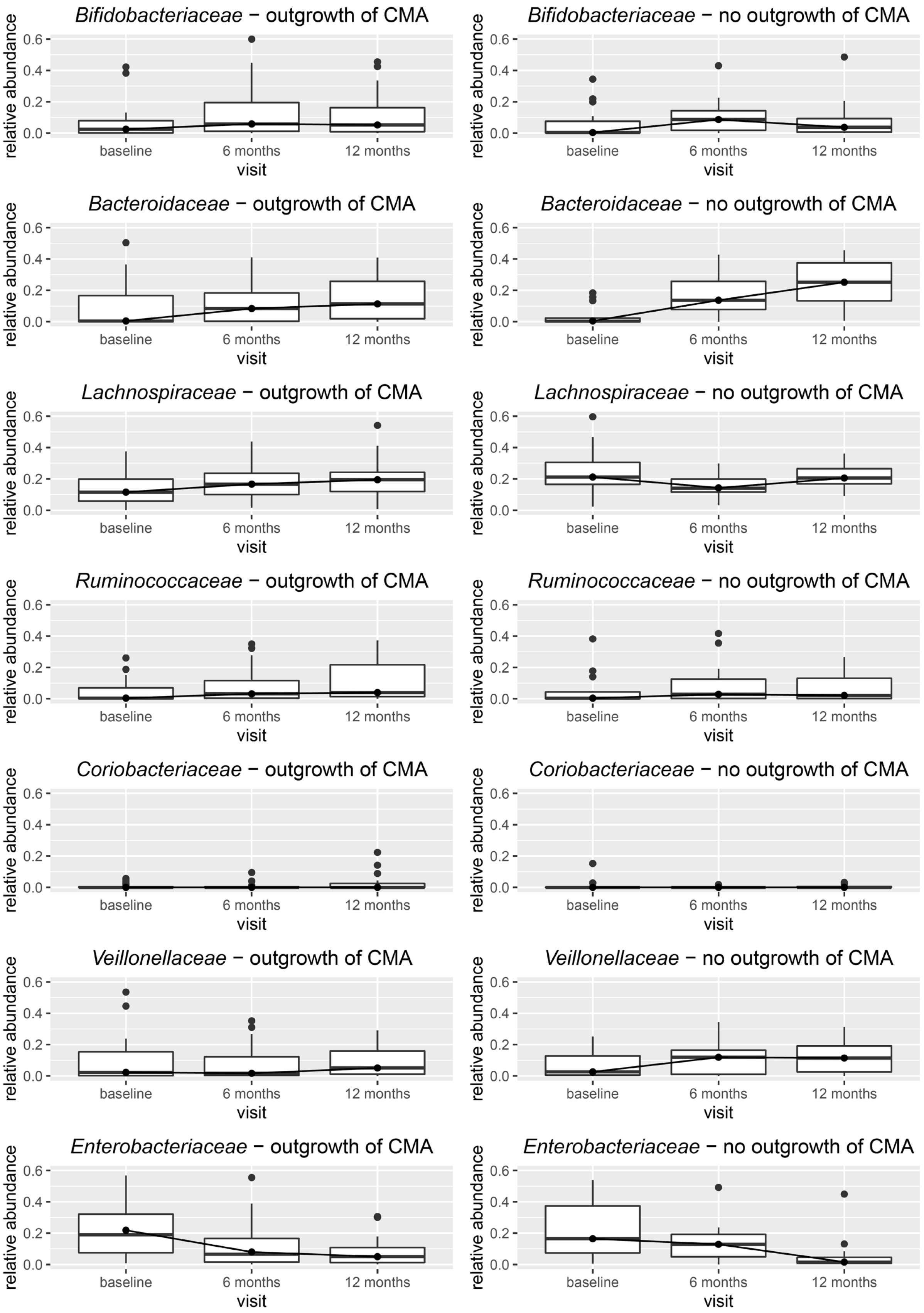

Figure S4. Boxplots of 16S rRNA gene-based relative abundances of core taxa over time.

Table S8. Difference in 16S rRNA gene based taxa at family level between visits within the group which outgrew their CMA as determined by Linear Mixed Model (LMM) analysis with outgrowth of CMA, visit and outgrowth of CMA x visit as fixed effects and subject as random effect. Abbreviation: p-adj.: adjusted p-value (p-value corrected for multiple testing using the Benjamini-Hochberg correction). Significant features ordered by adjusted p-value. In case of less than 10 significant features, the top 10 ordered by unadjusted p-value is presented. Bold = significant after multiple testing correction.

| **Outgrowth of CMA – visit 6 months vs baseline** | | | |
|---|---|---|---|
| **Family** | **p-value LMM** | **p-adj. LMM** | **Found by LEfSe** |
| Muribaculaceae | 0.0001 | **0.0018** | |
| RF39_fa | 0.0001 | **0.0018** | |
| Defluviitaleaceae | 0.0001 | **0.0018** | |
| Atopobiaceae | 0.0004 | **0.0045** | |
| Clostridia_unclassified | 0.0005 | **0.0045** | |
| Staphylococcaceae | 0.0017 | **0.0130** | yes |
| Micrococcaceae | 0.0022 | **0.0156** | |
| Oscillospiraceae | 0.0034 | **0.0215** | yes |
| Enterobacterales_unclassified | 0.0053 | **0.0301** | |
| Enterobacteriaceae | 0.0058 | **0.0301** | yes |
| Eubacteriaceae | 0.0062 | **0.0301** | yes |
| **Outgrowth of CMA – visit 12 months vs baseline** | | | |
| **Family** | **p-value LMM** | **p-adj. LMM** | **Found by LEfSe** |
| Muribaculaceae | 2.57E-10 | **3.24E-09** | |
| RF39_fa | 2.57E-10 | **3.24E-09** | |
| Defluviitaleaceae | 2.57E-10 | **3.24E-09** | |
| Clostridia_unclassified | 3.73E-07 | **3.92E-06** | |
| Comamonadaceae | 6.11E-07 | **5.50E-06** | |
| Enterobacteriaceae | 4.79E-06 | **3.65E-05** | yes |
| Atopobiaceae | 5.22E-06 | **3.65E-05** | |
| Micrococcaceae | 9.60E-06 | **6.04E-05** | yes |
| Enterobacterales_unclassified | 1.14E-05 | **6.52E-05** | yes |
| Rikenellaceae | 0.0001 | **0.0008** | yes |
| Neisseriaceae | 0.0002 | **0.0010** | |
| Staphylococcaceae | 0.0013 | **0.0056** | |
| Ruminococcaceae | 0.0052 | **0.0219** | yes |
| Enterococcaceae | 0.0068 | **0.0269** | |
| Butyricicoccaceae | 0.0096 | **0.0354** | yes |
| Oscillospiraceae | 0.0110 | **0.0385** | yes |
| Oscillospirales_fa | 0.0120 | **0.0397** | yes |
| **Outgrowth of CMA – visit 12 months vs 6 months** | | | |
| **Family** | **p-value LMM** | **p-adj. LMM** | **Found by LEfSe** |
| Comamonadaceae | 0.0032 | 0.0642 | |
| Muribaculaceae | 0.0061 | 0.0642 | |
| RF39_fa | 0.0061 | 0.0642 | |
| Defluviitaleaceae | 0.0061 | 0.0642 | |
| Aerococcaceae | 0.0079 | 0.0714 | |
| Clostridia_UCG.014_fa | 0.0249 | 0.1887 | yes |
| Rikenellaceae | 0.0270 | 0.1887 | yes |
| Neisseriaceae | 0.0463 | 0.2916 | |
| Enterobacteriaceae | 0.1138 | 0.6518 | |
| Clostridia_unclassified | 0.1482 | 0.7779 | |

Table S9. Difference in 16S rRNA gene based taxa at family level between visits within the group which did not outgrew their CMA as determined by Linear Mixed Model (LMM) analysis with outgrowth of CMA, visit and outgrowth of CMA x visit as fixed effects and subject as random effect. Abbreviation: p-adj.: adjusted p-value (p-value corrected for multiple testing using the Benjamini-Hochberg correction). Significant features ordered by adjusted p-value. In case of less than 10 significant features, the top 10 ordered by unadjusted p-value is presented. Bold = significant after multiple testing correction.

| **No outgrowth of CMA – visit 6 months vs baseline** | | | |
|---|---|---|---|
| **Family** | **p-value LMM** | **p-adj. LMM** | **Found by LEfSe** |
| *Butyricicoccaceae* | 0.0104 | 0.6522 | yes |
| *Bacteroidaceae* | 0.0234 | 0.6946 | yes |
| *Actinomycetaceae* | 0.0331 | 0.6946 | yes |
| *Acidaminococcaceae* | 0.0557 | 0.8766 | |
| *Ruminococcaceae* | 0.1103 | 0.9997 | |
| *Clostridiaceae* | 0.1333 | 0.9997 | |
| *Pasteurellaceae* | 0.1434 | 0.9997 | |
| *Morganellaceae* | 0.1500 | 0.9997 | |
| *Peptostreptococcales.Tissierellales_fa* | 0.1662 | 0.9997 | |
| *Bifidobacteriaceae* | 0.2114 | 0.9997 | |
| **No outgrowth of CMA – visit 12 months vs baseline** | | | |
| **Family** | **p-value LMM** | **p-adj. LMM** | **Found by LEfSe** |
| *Enterobacteriaceae* | 0.0002 | **0.0066** | yes |
| *Bacteroidaceae* | 0.0002 | **0.0066** | yes |
| *Muribaculaceae* | 0.0091 | 0.0816 | |
| *RF39_fa* | 0.0091 | 0.0816 | |
| *Defluviitaleaceae* | 0.0091 | 0.0816 | |
| *Butyricicoccaceae* | 0.0122 | 0.0959 | yes |
| *Prevotellaceae* | 0.0180 | 0.1258 | yes |
| *Actinomycetaceae* | 0.0233 | 0.1465 | |
| *Micrococcaceae* | 0.0405 | 0.2318 | |
| *Carnobacteriaceae* | 0.0488 | 0.2563 | |
| **No outgrowth of CMA – visit 12 months vs 6 months** | | | |
| **Family** | **p-value LMM** | **p-adj. LMM** | **Found by LEfSe** |
| *Enterobacteriaceae* | 0.0086 | 0.5393 | yes |
| *Monoglobaceae* | 0.0707 | 0.6486 | yes |
| *Prevotellaceae* | 0.0790 | 0.6486 | yes |
| *Muribaculaceae* | 0.0824 | 0.6486 | |
| *RF39_fa* | 0.0824 | 0.6486 | |
| *Defluviitaleaceae* | 0.0824 | 0.6486 | |
| *Selenomonadaceae* | 0.1039 | 0.7273 | yes |
| *Oscillospirales_fa* | 0.2099 | 0.9996 | yes |
| *Fusobacteriaceae* | 0.2561 | 0.9996 | |
| *Bacteroidaceae* | 0.2906 | 0.9996 | |

Table S10. Akaike Information Criterion (AIC) values for Linear Mixed Models (LMM) fitted to protein-based based relative abundance of core families. Model A: model without age as fixed effect, model B: model including age as fixed effect. For each core family, the best model (lowest AIC) is indicated in bold.

| | **AIC – model A** | **AIC – model B** |
|---|---|---|
| ***Bifidobacteriaceae*** | **424.9814** | 429.6942 |
| ***Bacteroidaceae*** | **325.5471** | 332.1521 |
| ***Lachnospiraceae*** | **307.7449** | 314.8004 |
| ***Ruminococcaceae*** | **329.2437** | 366.3991 |
| ***Coriobacteriaceae*** | **503.6151** | 508.5317 |
| ***Veillonellaceae*** | **524.2340** | 529.1699 |
| ***Enterobacteriaceae*** | **424.9663** | 430.4048 |

Table S11. Median protein-based relative abundance and interquartile range (IQR) of core families in faeces samples at baseline and follow-up visits (6 months, 12 months) in infants that outgrew their CMA after 12 months (12M) and children that did not. adj. p-value: p-value determined by Linear Mixed Model (LMM) analysis with Benjamini-Hochberg correction for multiple testing. Core taxa were defined as taxa that have a relative abundance higher than 1% in at least 50% of the 16S rRNA gene or metaproteomics samples. A: model without age as fixed effect, B: model including age as fixed effect.

| | **Baseline visit** | | |
|---|---|---|---|
| | **Outgrowth of CMA** | **No outgrowth of CMA** | **adj. p-value** |
| ***Bifidobacteriaceae*** | 0.346 (0.186-0.543) | 0.363 (0.127-0.624) | A:0.4304; B:0.4623 |
| ***Bacteroidaceae*** | 0.024 (0.014-0.047) | 0.028 (0.016-0.064) | A:0.4558; B:0.5069 |
| ***Lachnospiraceae*** | 0.342 (0.256-0.450) | 0.441 (0.188-0.597) | A:0.4304; B:0.4623 |
| ***Ruminococcaceae*** | 0.029 (0.022-0.049) | 0.022 (0.020-0.035) | A:0.2125; B:0.2497 |
| ***Coriobacteriaceae*** | 0.025 (0.001-0.084) | 0.048 (0.013-0.096) | A:0.2839; B:0.3398 |
| ***Veillonellaceae*** | 0.002 (0.001-0.005) | 0.001 (0.000-0.003) | A:0.2832; B:0.2497 |
| ***Enterobacteriaceae*** | 0.012 (0.005-0.038) | 0.011 (0.005-0.021) | A:0.2125; B:0.2497 |
| | **Visit 6 months** | | |
| | **Outgrowth of CMA** | **No outgrowth of CMA** | **adj. p-value** |
| ***Bifidobacteriaceae*** | 0.359 (0.162-0.692) | 0.349 (0.159-0.639) | A:0.5676; B:0.6646 |
| ***Bacteroidaceae*** | 0.024 (0.017-0.043) | 0.028 (0.021-0.041) | A:0.4572; B:0.5095 |
| ***Lachnospiraceae*** | 0.421 (0.140-0.508) | 0.369 (0.236-0.550) | A:0.7562; B:0.7844 |
| ***Ruminococcaceae*** | 0.037 (0.019-0.076) | 0.027 (0.023-0.038) | A:0.3156; B:0.3371 |
| ***Coriobacteriaceae*** | 0.026 (0.004-0.103) | 0.025 (0.013-0.079) | A:0.4311; B:0.4937 |
| ***Veillonellaceae*** | 0.001 (0.000-0.003) | 0.003 (0.002-0.007) | A:0.3156; B:0.3371 |
| ***Enterobacteriaceae*** | 0.007 (0.004-0.010) | 0.005 (0.002-0.014) | A:0.4132; B:0.4870 |
| | **Visit 12 months** | | |
| | **Outgrowth of CMA** | **No outgrowth of CMA** | **adj. p-value** |
| ***Bifidobacteriaceae*** | 0.301 (0.133-0.658) | 0.347 (0.130-0.460) | A:0.6735; B:0.7277 |
| ***Bacteroidaceae*** | 0.029 (0.018-0.047) | 0.032 (0.016-0.054) | A:0.6735; B:0.7277 |
| ***Lachnospiraceae*** | 0.298 (0.131-0.517) | 0.347 (0.315-0.493) | A:0.6735; B:0.7277 |
| ***Ruminococcaceae*** | 0.042 (0.013-0.088) | 0.035 (0.018-0.056) | A:0.6735; B:0.7277 |
| ***Coriobacteriaceae*** | 0.124 (0.041-0.181) | 0.105 (0.065-0.194) | A:0.6735; B:0.7277 |
| ***Veillonellaceae*** | 0.002 (0.001-0.004) | 0.002 (0.001-0.003) | A:0.6735; B:0.7277 |
| ***Enterobacteriaceae*** | 0.004 (0.002-0.009) | 0.003 (0.003-0.006) | A:0.6735; B:0.7277 |

Table S12. Difference in protein-based taxa at family level between allergy groups within visits as determined by Linear Mixed Model (LMM) analysis with outgrowth of CMA, visit and outgrowth of CMA x visit as fixed effects and subject as random effect. Abbreviation: p-adj.: adjusted p-value (p-value corrected for multiple testing using the Benjamini-Hochberg correction). Top 10 features ordered by unadjusted p-value.

| **Baseline visit – outgrowth vs no outgrowth of CMA** | | | |
|---|---|---|---|
| **Family** | **p-value LMM** | **p-adj. LMM** | **Found by LEfSe** |
| *Eggerthellaceae* | 0.009 | 0.178 | yes |
| *Ruminococcaceae* | 0.049 | 0.288 | |
| *Prevotellaceae* | 0.056 | 0.288 | |
| *Enterobacteriaceae* | 0.061 | 0.288 | |
| *Veillonellaceae* | 0.121 | 0.393 | |
| *Clostridiaceae* | 0.124 | 0.393 | |
| *Coriobacteriaceae* | 0.162 | 0.440 | |
| *Erysipelatoclostridiaceae* | 0.194 | 0.460 | |
| *Bifidobacteriaceae* | 0.308 | 0.651 | |
| *Lachnospiraceae* | 0.369 | 0.701 | |
| **Visit 6 months – outgrowth vs no outgrowth of CMA** | | | |
| **Family** | **p-value LMM** | **p-adj. LMM** | **Found by LEfSe** |
| *Prevotellaceae* | 0.011 | 0.209 | |
| *Veillonellaceae* | 0.071 | 0.571 | yes |
| *Ruminococcaceae* | 0.090 | 0.571 | |
| *Enterobacteriaceae* | 0.177 | 0.715 | |
| *Tannerellaceae* | 0.243 | 0.715 | |
| *Coriobacteriaceae* | 0.246 | 0.715 | |
| *Bacteroidaceae* | 0.327 | 0.715 | |
| *Rikenellaceae* | 0.359 | 0.715 | |
| *Erysipelatoclostridiaceae* | 0.370 | 0.715 | |
| *Eggerthellaceae* | 0.417 | 0.715 | |
| **Visit 12 months – outgrowth vs no outgrowth of CMA** | | | |
| **Family** | **p-value LMM** | **p-adj. LMM** | **Found by LEfSe** |
| *Oscillospiraceae* | 0.125 | 0.853 | |
| *Lactobacillaceae* | 0.220 | 0.853 | |
| *Tannerellaceae* | 0.379 | 0.853 | |
| *Enterobacteriaceae* | 0.385 | 0.853 | |
| *Prevotellaceae* | 0.487 | 0.853 | |
| *Erysipelatoclostridiaceae* | 0.497 | 0.853 | |
| *Clostridiaceae* | 0.508 | 0.853 | |
| *Eggerthellaceae* | 0.508 | 0.853 | |
| *Coriobacteriaceae* | 0.533 | 0.853 | |
| *Bifidobacteriaceae* | 0.545 | 0.853 | |

Table S13. Significance of difference in protein-based core taxa between visits within each allergy group as determined by Linear Mixed Model (LMM) analysis with Benjamini-Hochberg correction. Upper part of the table: infants that outgrew their CMA at 12 months; lower part of the table: infants that did not outgrew their CMA at 12 months. Abbreviations: Med.(IQR): median relative abundance and interquartile range; 0M: 0 months (baseline); 6M: 6 months; 12M: 12 months; p-adj.: adjusted p-value (p-value corrected for multiple testing using the Benjamini-Hochberg correction). A: model without age as fixed effect, B: model including age as fixed effect.

| **Infants that outgrew their CMA at visit 12 months** | | | | | | |
|---|---|---|---|---|---|---|
| | **Med.(IQR) 0M** | **Med.(IQR) 6M** | **Med.(IQR) 12M** | **p-adj. 0M vs 6M** | **p-adj. 0M vs 12M** | **p-adj. 6M vs 12M** |
| ***Bifidobacteriaceae*** | 0.346 (0.186-0.543) | 0.359 (0.162-0.692) | 0.301 (0.133-0.658) | A:0.9649 B:0.9936 | A:0.3228 B:0.9786 | A:0.7998 B:0.9977 |
| ***Bacteroidaceae*** | 0.024 (0.014-0.047) | 0.024 (0.017-0.043) | 0.029 (0.018-0.047) | A:0.9649 B:0.9936 | A:0.3228 B:0.9786 | A:0.8224 B:0.9977 |
| ***Lachnospiraceae*** | 0.342 (0.256-0.450) | 0.421 (0.140-0.508) | 0.298 (0.131-0.517) | A:0.9649 B:0.9936 | A:0.1488 B:0.9786 | A:0.6659 B:0.9977 |
| ***Ruminococcaceae*** | 0.029 (0.022-0.049) | 0.037 (0.019-0.076) | 0.042 (0.013-0.088) | A:0.9649 B:0.9936 | A:0.3228 B:0.9786 | A:0.6659 B:0.9977 |
| ***Coriobacteriaceae*** | 0.025 (0.001-0.084) | 0.026 (0.004-0.103) | 0.124 (0.041-0.181) | A:0.9649 B:0.9936 | A:0.1232 B:0.9786 | A:0.1184 B:0.9977 |
| ***Veillonellaceae*** | 0.002 (0.001-0.005) | 0.001 (0.000-0.003) | 0.002 (0.001-0.004) | A:0.9649 B:0.9936 | A:0.8621 B:0.9786 | A:0.6659 B:0.9977 |
| ***Enterobacteriaceae*** | 0.012 (0.005-0.038) | 0.007 (0.004-0.010) | 0.004 (0.002-0.009) | A:0.9649 B:0.9936 | **A:0.0141** B:0.9786 | A:0.6659 B:0.9977 |
| **Infants that did not outgrew their CMA at visit 12 months** | | | | | | |
| | **Med.(IQR) 0M** | **Med.(IQR) 6M** | **Med.(IQR) 12M** | **p-adj. 0M vs 6M** | **p-adj. 0M vs 12M** | **p-adj. 6M vs 12M** |
| ***Bifidobacteriaceae*** | 0.363 (0.127-0.624) | 0.349 (0.159-0.639) | 0.347 (0.130-0.460) | A:0.9985 B:0.9987 | A:0.8975 B:0.9980 | A:0.9999 B:0.9994 |
| ***Bacteroidaceae*** | 0.028 (0.016-0.064) | 0.028 (0.021-0.041) | 0.032 (0.016-0.054) | A:0.9985 B:0.9987 | A:0.8975 B:0.9980 | A:0.9999 B:0.9994 |
| ***Lachnospiraceae*** | 0.441 (0.188-0.597) | 0.369 (0.236-0.550) | 0.347 (0.315-0.493) | A:0.9985 B:0.9987 | A:0.9991 B:0.9980 | A:0.9999 B:0.9994 |
| ***Ruminococcaceae*** | 0.022 (0.020-0.035) | 0.027 (0.023-0.038) | 0.035 (0.018-0.056) | A:0.9985 B:0.9987 | A:0.9991 B:0.9980 | A:0.9999 B:0.9994 |
| ***Coriobacteriaceae*** | 0.048 (0.013-0.096) | 0.025 (0.013-0.079) | 0.105 (0.065-0.194) | A:0.9985 B:0.9987 | A:0.8975 B:0.9980 | A:0.9999 B:0.9994 |
| ***Veillonellaceae*** | 0.001 (0.000-0.003) | 0.003 (0.002-0.007) | 0.002 (0.001-0.003) | A:0.2456 B:0.9812 | A:0.8975 B:0.9980 | A:0.9999 B:0.9994 |
| ***Enterobacteriaceae*** | 0.011 (0.005-0.021) | 0.005 (0.002-0.014) | 0.003 (0.003-0.006) | A:0.9985 B:0.9987 | A:0.8975 B:0.9980 | A:0.9999 B:0.9994 |

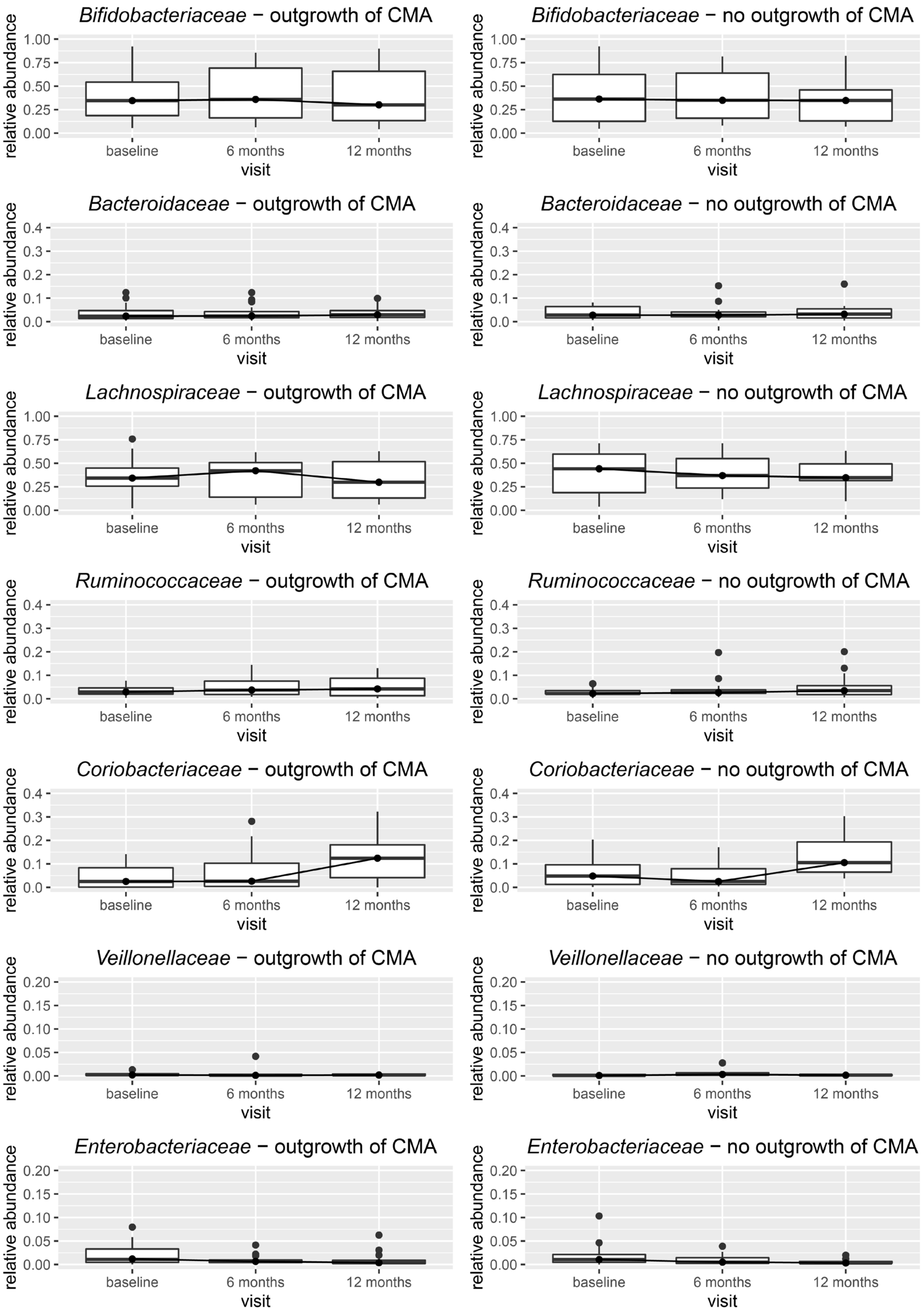

Figure S5. Boxplots of protein-based relative abundances of core taxa over time.

Table S14. Difference in protein-based taxa at family level between visits within the group which outgrew their CMA as determined by Linear Mixed Model (LMM) analysis with outgrowth of CMA, visit and outgrowth of CMA x visit as fixed effects and subject as random effect. Abbreviation: p-adj.: adjusted p-value (p-value corrected for multiple testing using the Benjamini-Hochberg correction). Top 10 features ordered by unadjusted p-value. Bold = significant after multiple testing correction.

| **Outgrowth of CMA – visit 6 months vs baseline** | | | |
|---|---|---|---|
| **Family** | **p-value LMM** | **p-adj. LMM** | **Found by LEfSe** |
| *Eggerthellaceae* | 0.025 | 0.479 | |
| *Enterobacteriaceae* | 0.145 | 0.992 | |
| *Prevotellaceae* | 0.277 | 0.992 | |
| *Acidaminococcaceae* | 0.319 | 0.992 | |
| *Enterococcaceae* | 0.339 | 0.992 | |
| *Rikenellaceae* | 0.516 | 0.992 | |
| *Lachnospiraceae* | 0.552 | 0.992 | |
| *Bacteroidaceae* | 0.591 | 0.992 | |
| *Erysipelatoclostridiaceae* | 0.606 | 0.992 | |
| *Bifidobacteriaceae* | 0.642 | 0.992 | |
| **Outgrowth of CMA – visit 12 months vs baseline** | | | |
| **Family** | **p-value LMM** | **p-adj. LMM** | **Found by LEfSe** |
| *Enterobacteriaceae* | 0.002 | **0.037** | yes |
| *Enterococcaceae* | 0.004 | **0.037** | |
| *Coriobacteriaceae* | 0.035 | 0.223 | yes |
| *Lachnospiraceae* | 0.064 | 0.275 | |
| *Rikenellaceae* | 0.072 | 0.275 | yes |
| *Eggerthellaceae* | 0.100 | 0.318 | |
| *Bifidobacteriaceae* | 0.201 | 0.545 | |
| *Bacteroidaceae* | 0.262 | 0.584 | |
| *Ruminococcaceae* | 0.277 | 0.584 | |
| *Tannerellaceae* | 0.629 | 0.998 | |
| **Outgrowth of CMA – visit 12 months vs 6 months** | | | |
| **Family** | **p-value LMM** | **p-adj. LMM** | **Found by LEfSe** |
| *Coriobacteriaceae* | 0.017 | 0.321 | yes |
| *Prevotellaceae* | 0.089 | 0.842 | yes |
| *Enterococcaceae* | 0.142 | 0.902 | |
| *Enterobacteriaceae* | 0.238 | 1.000 | |
| *Tannerellaceae* | 0.301 | 1.000 | |
| *Ruminococcaceae* | 0.406 | 1.000 | |
| *Lachnospiraceae* | 0.431 | 1.000 | |
| *Veillonellaceae* | 0.476 | 1.000 | |
| *Rikenellaceae* | 0.498 | 1.000 | |
| *Acidaminococcaceae* | 0.533 | 1.000 | |

Table S15. Difference in protein-based taxa at family level between visits within the group which did not outgrew their CMA as determined by Linear Mixed Model (LMM) analysis with outgrowth of CMA, visit and outgrowth of CMA x visit as fixed effects and subject as random effect. Abbreviation: p-adj.: adjusted p-value (p-value corrected for multiple testing using the Benjamini-Hochberg correction). Top 10 features ordered by unadjusted p-value.

| **No outgrowth of CMA – visit 6 months vs baseline** | | | |
|---|---|---|---|
| **Family** | **p-value LMM** | **p-adj. LMM** | **Found by LEfSe** |
| *Veillonellaceae* | 0.035 | 0.667 | yes |
| *Clostridiaceae* | 0.195 | 1.000 | yes |
| *Rikenellaceae* | 0.238 | 1.000 | |
| *Tannerellaceae* | 0.510 | 1.000 | |
| *Enterococcaceae* | 0.543 | 1.000 | |
| *Akkermansiaceae* | 0.572 | 1.000 | |
| *Bacteroidaceae* | 0.576 | 1.000 | |
| *Enterobacteriaceae* | 0.594 | 1.000 | |
| *Oscillospiraceae* | 0.626 | 1.000 | |
| *Acidaminococcaceae* | 0.651 | 1.000 | |
| **No outgrowth of CMA – visit 12 months vs baseline** | | | |
| **Family** | **p-value LMM** | **p-adj. LMM** | **Found by LEfSe** |
| *Enterococcaceae* | 0.055 | 0.796 | yes |
| *Prevotellaceae* | 0.098 | 0.796 | |
| *Tannerellaceae* | 0.129 | 0.796 | |
| *Enterobacteriaceae* | 0.181 | 0.796 | yes |
| *Rikenellaceae* | 0.210 | 0.796 | |
| *Oscillospiraceae* | 0.313 | 0.805 | |
| *Eggerthellaceae* | 0.334 | 0.805 | |
| *Veillonellaceae* | 0.413 | 0.805 | |
| *Coriobacteriaceae* | 0.416 | 0.805 | yes |
| *Lactobacillaceae* | 0.424 | 0.805 | |
| **No outgrowth of CMA – visit 12 months vs 6 months** | | | |
| **Family** | **p-value LMM** | **p-adj. LMM** | **Found by LEfSe** |
| *Eggerthellaceae* | 0.166 | 1.000 | yes |
| *Coriobacteriaceae* | 0.204 | 1.000 | yes |
| *Prevotellaceae* | 0.291 | 1.000 | |
| *Enterococcaceae* | 0.403 | 1.000 | |
| *Veillonellaceae* | 0.422 | 1.000 | yes |
| *Lactobacillaceae* | 0.437 | 1.000 | |
| *Tannerellaceae* | 0.672 | 1.000 | |
| *Clostridiaceae* | 0.684 | 1.000 | |
| *Enterobacteriaceae* | 0.697 | 1.000 | |
| *Erysipelatoclostridiaceae* | 0.770 | 1.000 | |

Table S16. Akaike Information Criterion (AIC) values for Linear Mixed Models (LMM) fitted to top 10 microbial protein functional classes (KEGG Brite hierarchy level c). Model A: model without age as fixed effect, model B: model including age as fixed effect. For each functional class, the best model (lowest AIC) is indicated in bold.

| | **AIC – model A** | **AIC – model B** |
|---|---|---|
| ***Ribosome*** | **389.7372** | 393.1967 |
| ***Oxidative phosphorylation*** | **512.4582** | 514.2544 |
| ***ABC transporters*** | **403.3784** | 409.0291 |
| ***Glycolysis / Gluconeogenesis*** | **261.7322** | 266.7850 |
| ***Purine metabolism*** | **382.7607** | 388.2610 |
| ***RNA degradation*** | **313.6898** | 314.2276 |
| ***Aminoacyl tRNA biosynthesis*** | **483.9525** | 487.8248 |
| ***Fructose and mannose metabolism*** | **460.9811** | 464.6671 |
| ***Pyruvate metabolism*** | **321.6124** | 328.6667 |
| ***Pentose and glucuronate interconversions*** | 569.1572 | **567.4152** |

Table S17. Median relative abundance and interquartile range (IQR) of top 10 microbial protein functional classes (KEGG Brite hierarchy level c) in faeces samples at baseline and follow-up visits (6 months, 12 months) in infants that outgrew their CMA after 12 months (12M) and children that did not. adj. p-value: p-value determined by Linear Mixed Model (LMM) analysis with Benjamini-Hochberg correction for multiple testing. A: model without age as fixed effect, B: model including age as fixed effect.

| | **Baseline visit** | | |
|---|---|---|---|
| | **Outgrowth of CMA** | **No outgrowth of CMA** | **adj. p-value** |
| ***Ribosome*** | 0.042 (0.020-0.115) | 0.073 (0.049-0.136) | A:0.5704; B:0.5576 |
| ***Oxidative phosphorylation*** | 0.001 (0.000-0.003) | 0.001 (0.000-0.002) | A:0.9732; B:0.8056 |
| ***ABC transporters*** | 0.003 (0.001-0.007) | 0.005 (0.002-0.009) | A:0.8191; B:0.7399 |
| ***Glycolysis / Gluconeogenesis*** | **0.006 (0.003-0.008) Min=0.001, max=0.019** | **0.006 (0.003-0.008) Min=0.001, max = 0.024** | **A:0.0154; B:0.0285** |
| ***Purine metabolism*** | 0.003 (0.002-0.006) | 0.005 (0.002-0.007) | A:0.5704; B:0.6574 |
| ***RNA degradation*** | **0.006 (0.005-0.008)** | **0.004 (0.003-0.006)** | **A:0.0439;** B:0.0867 |
| ***Aminoacyl tRNA biosynthesis*** | 0.001 (0.000-0.004) | 0.003 (0.002-0.006) | A:0.5109; B:0.3949 |
| ***Fructose and mannose metabolism*** | 0.001 (0.000-0.004) | 0.002 (0.001-0.004) | A:0.5704; B:0.7144 |
| ***Pyruvate metabolism*** | 0.002 (0.001-0.004) | 0.004 (0.001-0.007) | A:0.2425; B:0.2654 |
| ***Pentose and glucuronate interconversions*** | **0.000 (0.000-0.001)** | **0.000 (0.000-0.002)** | **A:0.0439;** B:0.0867 |
| | **Visit 6 months** | | |
| | **Outgrowth of CMA** | **No outgrowth of CMA** | **adj. p-value** |
| ***Ribosome*** | 0.087 (0.053-0.161) | 0.103 (0.061-0.156) | A:0.9492; B:0.9427 |
| ***Oxidative phosphorylation*** | 0.002 (0.001-0.004) | 0.002 (0.001-0.003) | A:0.9492; B:0.9427 |
| ***ABC transporters*** | 0.008 (0.004-0.011) | 0.008 (0.003-0.012) | A:0.9492; B:0.9427 |
| ***Glycolysis / Gluconeogenesis*** | 0.007 (0.004-0.010) | 0.007 (0.004-0.008) | A:0.9492; B:0.9427 |
| ***Purine metabolism*** | 0.005 (0.003-0.007) | 0.005 (0.003-0.009) | A:0.9492; B:0.9427 |
| ***RNA degradation*** | 0.005 (0.003-0.006) | 0.004 (0.002-0.007) | A:0.9492; B:0.9427 |
| ***Aminoacyl tRNA biosynthesis*** | 0.004 (0.001-0.010) | 0.003 (0.002-0.008) | A:0.9492; B:0.9427 |
| ***Fructose and mannose metabolism*** | 0.002 (0.000-0.005) | 0.002 (0.001-0.003) | A:0.9492; B:0.9427 |
| ***Pyruvate metabolism*** | 0.005 (0.002-0.008) | 0.004 (0.002-0.006) | A:0.9492; B:0.9427 |
| ***Pentose and glucuronate interconversions*** | 0.001 (0.000-0.004) | 0.002 (0.000-0.003) | A:0.9492; B:0.9427 |
| | **Visit 12 months** | | |
| | **Outgrowth of CMA** | **No outgrowth of CMA** | **adj. p-value** |
| ***Ribosome*** | 0.083 (0.058-0.130) | 0.096 (0.058-0.150) | A:0.9694; B:0.9862 |
| ***Oxidative phosphorylation*** | 0.003 (0.001-0.006) | 0.003 (0.001-0.007) | A:0.9694; B:0.9862 |
| ***ABC transporters*** | 0.003 (0.002-0.009) | 0.004 (0.003-0.005) | A:0.9694; B:0.9862 |
| ***Glycolysis / Gluconeogenesis*** | 0.007 (0.005-0.011) | 0.005 (0.003-0.009) | A:0.9694; B:0.9862 |
| ***Purine metabolism*** | 0.004 (0.003-0.006) | 0.005 (0.004-0.007) | A:0.9694; B:0.9862 |
| ***RNA degradation*** | 0.004 (0.002-0.005) | 0.003 (0.002-0.004) | A:0.9694; B:0.9862 |
| ***Aminoacyl tRNA biosynthesis*** | 0.005 (0.002-0.009) | 0.005 (0.003-0.006) | A:0.9694; B:0.9862 |
| ***Fructose and mannose metabolism*** | 0.003 (0.002-0.006) | 0.004 (0.002-0.008) | A:0.9694; B:0.9862 |
| ***Pyruvate metabolism*** | 0.005 (0.003-0.007) | 0.003 (0.003-0.005) | A:0.9694; B:0.9862 |
| ***Pentose and glucuronate interconversions*** | 0.002 (0.001-0.003) | 0.002 (0.001-0.002) | A:0.9694; B:0.9862 |

Table S18. Difference in protein functional classes (KEGG Brite hierarchy level c) between allergy groups within visits as determined by Linear Mixed Model (LMM) analysis with outgrowth of CMA, visit and outgrowth of CMA x visit as fixed effects and subject as random effect. Abbreviation: p-adj.: adjusted p-value (p-value corrected for multiple testing using the Benjamini-Hochberg correction). Top 10 features ordered by unadjusted p-value.

| **Baseline visit – outgrowth vs no outgrowth of CMA** | | | |
|---|---|---|---|
| **Functional class** | **p-value LMM** | **p-adj. LMM** | **Found by LEfSe** |
| Pentose and glucuronate interconversions | 0.006 | 0.334 | yes |
| Inositol phosphate metabolism | 0.029 | 0.334 | yes |
| Pyrimidine metabolism | 0.031 | 0.334 | yes |
| Glycolysis / Gluconeogenesis | 0.032 | 0.334 | |
| Nicotinate and nicotinamide metabolism | 0.033 | 0.334 | |
| Selenocompound metabolism | 0.037 | 0.334 | yes |
| Glyoxylate and dicarboxylate metabolism | 0.039 | 0.334 | |
| Valine, leucine and isoleucine degradation | 0.040 | 0.334 | yes |
| Histidine metabolism | 0.041 | 0.334 | |
| Streptomycin biosynthesis | 0.043 | 0.334 | yes |
| **Visit 6 months – outgrowth vs no outgrowth of CMA** | | | |
| **Functional class** | **p-value LMM** | **p-adj. LMM** | **Found by LEfSe** |
| Amino sugar and nucleotide sugar metabolism | 0.006 | 0.488 | yes |
| Pyrimidine metabolism | 0.013 | 0.569 | yes |
| Glycine, serine and threonine metabolism | 0.023 | 0.656 | yes |
| Biosynthesis of ansamycins | 0.055 | 0.983 | |
| Lysine biosynthesis | 0.106 | 0.983 | |
| Starch and sucrose metabolism | 0.121 | 0.983 | |
| Glutathione metabolism | 0.138 | 0.983 | |
| Phosphotransferase system | 0.214 | 0.983 | |
| Vitamin B6 metabolism | 0.215 | 0.983 | |
| Pentose and glucuronate interconversions | 0.233 | 0.983 | |
| **Visit 12 months – outgrowth vs no outgrowth of CMA** | | | |
| **Functional class** | **p-value LMM** | **p-adj. LMM** | **Found by LEfSe** |
| Sphingolipid metabolism | 0.057 | 0.998 | |
| Other glycan degradation | 0.057 | 0.998 | |
| Amino sugar and nucleotide sugar metabolism | 0.088 | 0.998 | |
| Lysine biosynthesis | 0.095 | 0.998 | |
| Valine, leucine and isoleucine biosynthesis | 0.104 | 0.998 | |
| Pantothenate and CoA biosynthesis | 0.123 | 0.998 | |
| Methane metabolism | 0.130 | 0.998 | |
| Starch and sucrose metabolism | 0.144 | 0.998 | yes |
| Glutathione metabolism | 0.147 | 0.998 | |
| Biofilm formation | 0.193 | 0.998 | |

Table S19. Significance of difference in top 10 microbial protein functional classes (KEGG Brite hierarchy level c) between visits within each allergy group as determined by Linear Mixed Model (LMM) analysis with Benjamini-Hochberg correction. Upper part of the table: infants that outgrew their CMA at 12 months; lower part of the table: infants that did not outgrew their CMA at 12 months. Abbreviations: Med.(IQR): median relative abundance and interquartile range; 0M: 0 months (baseline); 6M: 6 months; 12M: 12 months; p-adj.: adjusted p-value (p-value corrected for multiple testing using the Benjamini-Hochberg correction). A: model without age as fixed effect, B: model including age as fixed effect.

| **Infants that outgrew their CMA at visit 12 months** | | | | | | |
|---|---|---|---|---|---|---|
| | **Med.(IQR) 0M** | **Med.(IQR) 6M** | **Med.(IQR) 12M** | **p-adj. 0M vs 6M** | **p-adj. 0M vs 12M** | **p-adj. 6M vs 12M** |
| ***Ribosome*** | 0.042 (0.020-0.115) | 0.087 (0.053-0.161) | 0.083 (0.058-0.130) | A:0.9904<br>B:0.8351 | A:0.9928<br>B:0.6884 | A:0.8819<br>B:1.0000 |
| ***Oxidative phosphorylation*** | 0.001 (0.000-0.003) | 0.002 (0.001-0.004) | 0.003 (0.001-0.006) | A:0.9904<br>B:0.8351 | A:0.2481<br>B:0.9973 | A:0.5793<br>B:1.0000 |
| ***ABC transporters*** | 0.003 (0.001-0.007) | 0.008 (0.004-0.011) | 0.003 (0.002-0.009) | A:0.9904<br>B:0.9912 | A:0.2481<br>B:0.4802 | A:0.1814<br>B:0.1814 |
| ***Glycolysis / Gluconeogenesis*** | 0.006 (0.003-0.008) | 0.007 (0.004-0.010) | 0.007 (0.005-0.011) | **A:0.0413**<br>B:0.0608 | **A:0.0039**<br>**B:0.0078** | A:0.8819<br>B:1.0000 |
| ***Purine metabolism*** | 0.003 (0.002-0.006) | 0.005 (0.003-0.007) | 0.004 (0.003-0.006) | A:0.9904<br>B:0.8351 | A:0.3657<br>B: 0.9973 | A:0.4538<br>B:1.0000 |
| ***RNA degradation*** | 0.006 (0.005-0.008) | 0.005 (0.003-0.006) | 0.004 (0.002-0.005) | **A:0.0283**<br>B:0.8351 | **A:< $10^{-4}$**<br>B:0.9973 | A:0.2859<br>B:1.0000 |
| ***Aminoacyl tRNA biosynthesis*** | 0.001 (0.000-0.004) | 0.004 (0.001-0.010) | 0.005 (0.002-0.009) | A:0.0892<br>B:0.0608 | A:0.1056<br>B:0.1458 | A:0.9820<br>B:1.0000 |
| ***Fructose and mannose metabolism*** | 0.001 (0.000-0.004) | 0.002 (0.000-0.005) | 0.003 (0.002-0.006) | A:0.9904<br>B:0.8351 | A:0.4031<br>B:0.9973 | A:0.7269<br>B:1.0000 |
| ***Pyruvate metabolism*** | 0.002 (0.001-0.004) | 0.005 (0.002-0.008) | 0.005 (0.003-0.007) | A:0.9904<br>B:0.8351: | A:0.2760<br>B:0.9973 | A:0.8819<br>B:1.0000 |
| ***Pentose and glucuronate interconversions*** | 0.000 (0.000-0.001) | 0.001 (0.000-0.004) | 0.002 (0.001-0.003) | A:0.9904<br>B:0.8351 | **A:0.0166**<br>**B:0.0248** | A:0.1814<br>B:0.1814 |
| **Infants that did not outgrew their CMA at visit 12 months** | | | | | | |
| | **Med.(IQR) 0M** | **Med.(IQR) 6M** | **Med.(IQR) 12M** | **p-adj. 0M vs 6M** | **p-adj. 0M vs 12M** | **p-adj. 6M vs 12M** |
| ***Ribosome*** | 0.073 (0.049-0.136) | 0.103 (0.061-0.156) | 0.096 (0.058-0.150) | A:0.9984<br>B:0.9946 | A:0.9922<br>B:0.9998 | A:0.9736<br>B:0.9936 |
| ***Oxidative phosphorylation*** | 0.001 (0.000-0.002) | 0.002 (0.001-0.003) | 0.003 (0.001-0.007) | A:0.9984<br>B:0.9946: | A:0.9922<br>B:0.9998 | A:0.9585<br>B:0.9936 |
| ***ABC transporters*** | 0.005 (0.002-0.009) | 0.008 (0.003-0.012) | 0.004 (0.003-0.005) | A:0.9984<br>B:0.9946: | A:0.9922<br>B:0.9998: | A:0.9585<br>B:0.9936 |
| ***Glycolysis / Gluconeogenesis*** | 0.006 (0.003-0.008) | 0.007 (0.004-0.008) | 0.005 (0.003-0.009) | A:0.9984<br>B:0.9946 | A:0.9922<br>B:0.9998 | A:0.9585<br>B:0.9936 |
| ***Purine metabolism*** | 0.005 (0.002-0.007) | 0.005 (0.003-0.009) | 0.005 (0.004-0.007) | A:0.9984<br>B:0.9946 | A:0.9922<br>B:0.9998 | A:0.9736<br>B:0.9936 |
| ***RNA degradation*** | 0.004 (0.003-0.006) | 0.004 (0.002-0.007) | 0.003 (0.002-0.004) | A:0.9984<br>B:0.9946 | A:0.6915<br>B:0.9998 | A:0.9585<br>B:0.9936 |
| ***Aminoacyl tRNA biosynthesis*** | 0.003 (0.002-0.006) | 0.003 (0.002-0.008) | 0.005 (0.003-0.006) | A:0.9984<br>B:0.9946 | A:0.9922<br>B:0.9998 | A:0.9736<br>B:0.9936 |
| ***Fructose and mannose metabolism*** | 0.002 (0.001-0.004) | 0.002 (0.001-0.003) | 0.004 (0.002-0.008) | A:0.9984<br>B:0.9946 | A:0.9922<br>B:0.9998: | A:0.9736<br>B:0.9936 |
| ***Pyruvate metabolism*** | 0.004 (0.001-0.007) | 0.004 (0.002-0.006) | 0.003 (0.003-0.005) | A:0.9984<br>B:0.9946 | A:0.9922<br>B:0.9998 | A:0.9736<br>B:0.9936 |
| ***Pentose and glucuronate interconversions*** | 0.000 (0.000-0.002) | 0.002 (0.000-0.003) | 0.002 (0.001-0.002) | A:0.9984<br>B:0.9946 | A:0.9922<br>B:0.9998 | A:0.9736<br>B:0.9936 |

Table S20. Difference in protein functional classes (KEGG Brite hierarchy level c) between visits within the group which outgrew their CMA as determined by Linear Mixed Model (LMM) analysis with outgrowth of CMA, visit and outgrowth of CMA x visit as fixed effects and subject as random effect. Abbreviation: p-adj.: adjusted p-value (p-value corrected for multiple testing using the Benjamini-Hochberg correction). Top 10 features ordered by unadjusted p-value. Bold = significant after multiple testing correction.

| **Outgrowth of CMA – visit 6 months vs baseline** | | | |
|---|---|---|---|
| **Functional class** | **p-value LMM** | **p-adj. LMM** | **Found by LEfSe** |
| Nicotinate and nicotinamide metabolism | 0.0052 | 0.3578 | |
| Aminoacyl-tRNA biosynthesis | 0.0095 | 0.3578 | yes |
| Selenocompound metabolism | 0.0294 | 0.5017 | yes |
| RNA degradation | 0.0295 | 0.5017 | |
| Two component system | 0.0410 | 0.5291 | |
| Carbon fixation pathways in prokaryotes | 0.0469 | 0.5291 | |
| Cyanoamino acid metabolism | 0.0558 | 0.5291 | |
| Riboflavin metabolism | 0.0560 | 0.5291 | |
| Biofilm formation | 0.1104 | 0.8074 | |
| Galactose metabolism | 0.1314 | 0.8074 | yes |
| **Outgrowth of CMA – visit 12 months vs baseline** | | | |
| **Functional class** | **p-value LMM** | **p-adj. LMM** | **Found by LEfSe** |
| Selenocompound metabolism | 0.0001 | **0.0104** | yes |
| RNA degradation | 0.0002 | **0.0104** | |
| Tryptophan metabolism | 0.0023 | **0.0484** | yes |
| Carbon fixation pathways in prokaryotes | 0.0028 | **0.0484** | |
| Propanoate metabolism | 0.0038 | 0.0513 | yes |
| Cyanoamino acid metabolism | 0.0044 | 0.0513 | |
| Pentose and glucuronate interconversions | 0.0048 | 0.0513 | yes |
| Lysine degradation | 0.0108 | 0.0876 | |
| Terpenoid backbone biosynthesis | 0.0108 | 0.0876 | |
| Aminoacyl-tRNA biosynthesis | 0.0113 | 0.0876 | yes |
| **Outgrowth of CMA – visit 12 months vs 6 months** | | | |
| **Functional class** | **p-value LMM** | **p-adj. LMM** | **Found by LEfSe** |
| Flagellar assembly | 0.0205 | 0.8566 | yes |
| Pentose and glucuronate interconversions | 0.0449 | 0.8566 | |
| Amino sugar and nucleotide sugar metabolism | 0.0558 | 0.8566 | yes |
| Sphingolipid metabolism | 0.0583 | 0.8566 | |
| Other glycan degradation | 0.0583 | 0.8566 | |
| Tryptophan metabolism | 0.0683 | 0.8566 | |
| ABC transporters | 0.0705 | 0.8566 | |
| Fatty acid degradation | 0.1208 | 0.9412 | |
| Vitamin B6 metabolism | 0.1326 | 0.9412 | |
| Lysine degradation | 0.1455 | 0.9412 | |

Table S21. Difference in protein functional classes (KEGG Brite hierarchy level c) between visits within the group which did not outgrew their CMA as determined by Linear Mixed Model (LMM) analysis with outgrowth of CMA, visit and outgrowth of CMA x visit as fixed effects and subject as random effect. Abbreviation: p-adj.: adjusted p-value (p-value corrected for multiple testing using the Benjamini-Hochberg correction). Top 10 features ordered by unadjusted p-value.

| **No outgrowth of CMA – visit 6 months vs baseline** | | | |
|---|---|---|---|
| **Functional class** | **p-value LMM** | **p-adj. LMM** | **Found by LEfSe** |
| Carbon fixation pathways in prokaryotes | 0.0272 | 0.9997 | |
| Amino sugar and nucleotide sugar metabolism | 0.0681 | 0.9997 | |
| Glutathione metabolism | 0.0831 | 0.9997 | yes |
| Flagellar assembly | 0.0936 | 0.9997 | |
| Taurine and hypotaurine metabolism | 0.1871 | 0.9997 | |
| Lysine degradation | 0.2173 | 0.9997 | |
| Terpenoid backbone biosynthesis | 0.2173 | 0.9997 | |
| Histidine metabolism | 0.2588 | 0.9997 | |
| Starch and sucrose metabolism | 0.2655 | 0.9997 | |
| Tryptophan metabolism | 0.2860 | 0.9997 | |
| **No outgrowth of CMA – visit 12 months vs baseline** | | | |
| **Functional class** | **p-value LMM** | **p-adj. LMM** | **Found by LEfSe** |
| Tryptophan metabolism | 0.0054 | 0.4564 | yes |
| Sphingolipid metabolism | 0.0169 | 0.4798 | |
| Other glycan degradation | 0.0169 | 0.4798 | |
| Carbon fixation pathways in prokaryotes | 0.0249 | 0.5287 | |
| Lysine biosynthesis | 0.1680 | 0.9999 | yes |
| Lysine degradation | 0.1766 | 0.9999 | |
| Terpenoid backbone biosynthesis | 0.1766 | 0.9999 | |
| Cyanoamino acid metabolism | 0.2079 | 0.9999 | |
| RNA degradation | 0.2534 | 0.9999 | |
| Glycine, serine and threonine metabolism | 0.2673 | 0.9999 | |
| **No outgrowth of CMA – visit 12 months vs 6 months** | | | |
| **Functional class** | **p-value LMM** | **p-adj. LMM** | **Found by LEfSe** |
| Flagellar assembly | 0.0058 | 0.2511 | yes |
| Sphingolipid metabolism | 0.0089 | 0.2511 | |
| Other glycan degradation | 0.0089 | 0.2511 | |
| Glutathione metabolism | 0.0188 | 0.3992 | yes |
| Starch and sucrose metabolism | 0.0590 | 0.9999 | yes |
| Amino sugar and nucleotide sugar metabolism | 0.0716 | 0.9999 | |
| Oxidative phosphorylation | 0.1352 | 0.9999 | |
| Biosynthesis of various plant secondary metabolites | 0.1765 | 0.9999 | |
| Tryptophan metabolism | 0.2137 | 0.9999 | |
| Pantothenate and CoA biosynthesis | 0.2164 | 0.9999 | yes |

Table S22. Akaike Information Criterion (AIC) values for Linear Mixed Models (LMM) fitted to top 10 human protein classes. Model A: model without age as fixed effect, model B: model including age as fixed effect. For each functional class, the best model (lowest AIC) is indicated in bold.

| | **AIC – model A** | **AIC – model B** |
|---|---|---|
| ***Immunoglobulins*** | **375.8744** | 376.0734 |
| ***Glycoside hydrolases*** | **426.9341** | 433.0265 |
| ***Transthyretin/hydroxyisourate hydrolases*** | **613.9620** | 614.0648 |
| ***Proline-rich proteins*** | **605.7402** | 609.7291 |
| ***S100 proteins*** | **501.4520** | 505.8600 |
| ***Secretory proteins*** | **540.8112** | 544.3607 |
| ***Serine peptidases*** | **470.7533** | 472.6541 |
| ***Carboxypeptidases*** | **387.2018** | 393.1329 |
| ***Dipeptidyl peptidases*** | **573.0248** | 576.8840 |
| ***Actin family*** | **528.2970** | 530.2747 |

Table S23. Median relative abundance and interquartile range (IQR) of top 10 human protein classes in faeces samples at baseline and follow-up visits (6 months, 12 months) in infants that outgrew their CMA after 12 months (12M) and children that did not. adj. p-value: p-value determined by Linear Mixed Model (LMM) analysis with Benjamini-Hochberg correction for multiple testing. A: model without age as fixed effect, B: model including age as fixed effect.

| | **Baseline visit** | | |
|---|---|---|---|
| | **Outgrowth of CMA** | **No outgrowth of CMA** | **adj. p-value** |
| ***Immunoglobulins*** | 0.530 (0.324-0.657) | 0.645 (0.544-0.770) | A:0.8517; B:0.5278 |
| ***Glycoside hydrolases*** | 0.009 (0.006-0.031) | 0.010 (0.006-0.021) | A:0.9510; B:0.9512 |
| ***Transthyretin/hydroxyisourate hydrolases*** | 0.013 (0.000-0.030) | 0.006 (0.000-0.025) | A:0.9510; B:0.9512 |
| ***Proline-rich proteins*** | 0.001 (0.000-0.007) | 0.001 (0.000-0.008) | A:0.9510; B:0.9512 |
| ***S100 proteins*** | 0.012 (0.007-0.025) | 0.006 (0.002-0.009) | A:0.8517; B:0.5278 |
| ***Secretory proteins*** | 0.002 (0.002-0.011) | 0.003 (0.000-0.016) | A:0.9510; B:0.9512 |
| ***Serine peptidases*** | 0.167 (0.033-0.384) | 0.128 (0.073-0.235) | A:0.9510; B:0.9512 |
| ***Carboxypeptidases*** | 0.007 (0.001-0.015) | 0.004 (0.001-0.013) | A:0.9510; B:0.9512 |
| ***Dipeptidyl peptidases*** | 0.003 (0.001-0.007) | 0.001 (0.000-0.009) | A:0.9510; B:0.9512 |
| ***Actin family*** | 0.001 (0.000-0.003) | 0.001 (0.000-0.002) | A:0.9510; B:0.9512 |
| | **Visit 6 months** | | |
| | **Outgrowth of CMA** | **No outgrowth of CMA** | **adj. p-value** |
| ***Immunoglobulins*** | 0.497 (0.295-0.639) | 0.593 (0.461-0.746) | A:0.7298; B:0.9071 |
| ***Glycoside hydrolases*** | 0.014 (0.010-0.048) | 0.025 (0.008-0.042) | A:0.7298; B:0.7697 |
| ***Transthyretin/hydroxyisourate hydrolases*** | 0.002 (0.000-0.022) | 0.013 (0.002-0.022) | A:0.4709; B:0.6573 |
| ***Proline-rich proteins*** | 0.005 (0.001-0.018) | 0.008 (0.006-0.019) | A:0.4709; B:0.6573 |
| ***S100 proteins*** | 0.005 (0.001-0.017) | 0.002 (0.001-0.009) | A:0.7298; B:0.7697 |
| ***Secretory proteins*** | 0.004 (0.002-0.008) | 0.010 (0.002-0.021) | A:0.7298; B:0.7697 |
| ***Serine peptidases*** | 0.213 (0.152-0.424) | 0.110 (0.070-0.324) | A:0.3296; B:0.5091 |
| ***Carboxypeptidases*** | 0.003 (0.002-0.011) | 0.007 (0.003-0.011) | A:0.7298; B:0.6798 |
| ***Dipeptidyl peptidases*** | 0.002 (0.000-0.008) | 0.001 (0.000-0.003) | A:0.4709; B:0.5932 |
| ***Actin family*** | 0.001 (0.000-0.005) | 0.004 (0.001-0.010) | A:0.4709; B:0.5932 |
| | **Visit 12 months** | | |
| | **Outgrowth of CMA** | **No outgrowth of CMA** | **adj. p-value** |
| ***Immunoglobulins*** | 0.514 (0.366-0.636) | 0.570 (0.431-0.672) | A:0.9011; B:0.6762 |
| ***Glycoside hydrolases*** | 0.032 (0.015-0.045) | 0.043 (0.034-0.084) | A:0.4855; B:0.4862 |
| ***Transthyretin/hydroxyisourate hydrolases*** | 0.016 (0.007-0.024) | 0.006 (0.002-0.020) | A:0.4892; B:0.4862 |
| ***Proline-rich proteins*** | 0.010 (0.004-0.065) | 0.028 (0.015-0.046) | A:0.4855; B:0.4862 |
| ***S100 proteins*** | 0.010 (0.003-0.037) | 0.007 (0.002-0.018) | A:0.4855; B:0.4862 |
| ***Secretory proteins*** | 0.002 (0.001-0.007) | 0.003 (0.002-0.006) | A:0.8664; B:0.6762 |
| ***Serine peptidases*** | 0.160 (0.122-0.379) | 0.135 (0.091-0.196) | A:0.4855; B:0.4862 |
| ***Carboxypeptidases*** | 0.012 (0.004-0.025) | 0.017 (0.010-0.028) | A:0.4855; B:0.4862 |
| ***Dipeptidyl peptidases*** | 0.013 (0.007-0.022) | 0.005 (0.001-0.026) | A:0.4855; B:0.4862 |
| ***Actin family*** | 0.002 (0.001-0.005) | 0.002 (0.001-0.014) | A:0.4855; B:0.4862 |

Table S24. Difference in human protein classes between allergy groups within visits as determined by Linear Mixed Model (LMM) analysis with outgrowth of CMA, visit and outgrowth of CMA x visit as fixed effects and subject as random effect. Abbreviation: p-adj.: adjusted p-value (p-value corrected for multiple testing using the Benjamini-Hochberg correction). Top 10 features ordered by unadjusted p-value.

| **Baseline visit – outgrowth vs no outgrowth of CMA** | | | |
|---|---|---|---|
| **Human protein class** | **p-value LMM** | **p-adj. LMM** | **Found by LEfSe** |
| Cell adhesion proteins | 0.019 | 0.977 | |
| Mucin family | 0.048 | 0.977 | |
| Superoxide dismutases | 0.054 | 0.977 | |
| F ATPases | 0.071 | 0.977 | |
| Annexins | 0.096 | 0.977 | |
| S100 proteins | 0.100 | 0.977 | yes |
| Elongation factors | 0.122 | 0.977 | |
| Alkaline phosphatases | 0.152 | 0.977 | yes |
| Binding protein | 0.158 | 0.977 | |
| Transmembrane ATPases | 0.190 | 0.977 | |
| **Visit 6 months – outgrowth vs no outgrowth of CMA** | | | |
| **Human protein class** | **p-value LMM** | **p-adj. LMM** | **Found by LEfSe** |
| Peroxidases | 0.002 | 0.148 | |
| Exosomal proteins | 0.004 | 0.177 | yes |
| Actin family | 0.066 | 0.921 | yes |
| Carboxylic ester hydrolases | 0.082 | 0.921 | |
| Lectins | 0.098 | 0.921 | |
| F-ATPases | 0.098 | 0.921 | |
| CUB-domain containing proteins | 0.103 | 0.921 | |
| Titin | 0.103 | 0.921 | |
| Phosphoglycerate kinase | 0.110 | 0.921 | |
| Complement C3 like proteins | 0.114 | 0.921 | |
| **Visit 12 months – outgrowth vs no outgrowth of CMA** | | | |
| **Human protein class** | **p-value LMM** | **p-adj. LMM** | **Found by LEfSe** |
| Serpins | 0.004 | 0.322 | yes |
| Tumor suppressor | 0.017 | 0.803 | |
| Adenosine deaminases | 0.064 | 0.933 | |
| P-ATPases | 0.082 | 0.933 | yes |
| Cadherins | 0.086 | 0.933 | |
| Ubiquitins | 0.092 | 0.933 | |
| Bile salt activated lipase | 0.099 | 0.933 | |
| F-ATPases | 0.111 | 0.933 | |
| Submaxillary gland androgen regulated proteins | 0.140 | 0.933 | |
| CUB-domain containing proteins | 0.153 | 0.933 | |

Table S25. Significance of difference in top 10 human protein classes between visits within each allergy group as determined by Linear Mixed Model (LMM) analysis with Benjamini-Hochberg correction. Upper part of the table: infants that outgrew their CMA at 12 months; lower part of the table: infants that did not outgrew their CMA at 12 months. Abbreviations: Med.(IQR): median relative abundance and interquartile range; 0M: 0 months (baseline); 6M: 6 months; 12M: 12 months; p-adj.: adjusted p-value (p-value corrected for multiple testing using the Benjamini-Hochberg correction). A: model without age as fixed effect, B: model including age as fixed effect.

| **Infants that outgrew their CMA at visit 12 months** | | | | | | |
|---|---|---|---|---|---|---|
| | **Med.(IQR) 0M** | **Med.(IQR) 6M** | **Med.(IQR) 12M** | **p-adj. 0M vs 6M** | **p-adj. 0M vs 12M** | **p-adj. 6M vs 12M** |
| ***Immunoglobulins*** | 0.530 (0.324-0.657) | 0.497 (0.295-0.639) | 0.514 (0.366-0.636) | A:1.0000 B:0.5077 | A:0.0527 B:0.9965 | A:0.0764 B:0.9595 |
| ***Glycoside hydrolases*** | 0.009 (0.006-0.031) | 0.014 (0.010-0.048) | 0.032 (0.015-0.045) | A:1.0000 B:0.8435 | A:0.9863 B:0.9965 | A:0.9051 B:0.9595 |
| ***Transthyretin/hydroxyiso urate hydrolases*** | 0.013 (0.000-0.030) | 0.002 (0.000-0.022) | 0.016 (0.007-0.024) | A:1.0000 B:0.1935 | A:0.5965 B:0.9159 | A:0.0764 B:0.9595 |
| ***Proline-rich proteins*** | 0.001 (0.000-0.007) | 0.005 (0.001-0.018) | 0.010 (0.004-0.065) | A:0.2247 B:0.5965 | **A:0.0339** B:0.9159 | A:0.9051 B:0.9595 |
| ***S100 proteins*** | 0.012 (0.007-0.025) | 0.005 (0.001-0.017) | 0.010 (0.003-0.037) | **A:0.0429** B:0.1103 | A:0.0527 B:0.9159 | A:0.9398 B:0.9595 |
| ***Secretory proteins*** | 0.002 (0.002-0.011) | 0.004 (0.002-0.008) | 0.002 (0.001-0.007) | A:1.0000 B:0.8435 | A:0.4763 B:0.9965 | A:0.3249 B:0.9595 |
| ***Serine peptidases*** | 0.167 (0.033-0.384) | 0.213 (0.152-0.424) | 0.160 (0.122-0.379) | A:1.0000 B:0.2652 | A:0.9099 B:0.9159 | A:0.3223 B:0.9595 |
| ***Carboxypeptidases*** | 0.007 (0.001-0.015) | 0.003 (0.002-0.011) | 0.012 (0.004-0.025) | A:1.0000 B:0.8435 | A:0.9099 B:0.9159 | A:0.9398 B:0.9595 |
| ***Dipeptidyl peptidases*** | 0.003 (0.001-0.007) | 0.002 (0.000-0.008) | 0.013 (0.007-0.022) | A:1.0000 B:0.8435 | A:0.4050 B:0.9965 | A:0.3249 B:0.9595 |
| ***Actin family*** | 0.001 (0.000-0.003) | 0.001 (0.000-0.005) | 0.002 (0.001-0.005) | A:1.0000 B:0.8329 | A:0.9099 B:0.9159 | A:0.9051 B:0.9595 |
| **Infants that did not outgrew their CMA at visit 12 months** | | | | | | |
| | **Med.(IQR) 0M** | **Med.(IQR) 6M** | **Med.(IQR) 12M** | **p-adj. 0M vs 6M** | **p-adj. 0M vs 12M** | **p-adj. 6M vs 12M** |
| ***Immunoglobulins*** | 0.645 (0.544-0.770) | 0.593 (0.461-0.746) | 0.570 (0.431-0.672) | A:0.5769 B:0.9994 | **A:0.0222** B:0.9985 | A:0.9798 B:0.9994 |
| ***Glycoside hydrolases*** | 0.010 (0.006-0.021) | 0.025 (0.008-0.042) | 0.043 (0.034-0.084) | A:0.9968 B:0.9994 | A:0.9998 B:0.9985 | A:0.9798 B:0.9994 |
| ***Transthyretin/hydroxyiso urate hydrolases*** | 0.006 (0.000-0.025) | 0.013 (0.002-0.022) | 0.006 (0.002-0.020) | A:0.9968 B:0.6372 | A:0.9998 B:0.8230 | A:0.9798 B:0.9994 |
| ***Proline-rich proteins*** | 0.001 (0.000-0.008) | 0.008 (0.006-0.019) | 0.028 (0.015-0.046) | A:0.5769 B:0.5256 | A:0.0562 B:0.9706 | A:0.9798 B:0.9994 |
| ***S100 proteins*** | 0.006 (0.002-0.009) | 0.002 (0.001-0.009) | 0.007 (0.002-0.018) | A:0.5769 B:0.5256 | A:0.4603 B:0.8230 | A:0.9798 B:0.9994 |
| ***Secretory proteins*** | 0.003 (0.000-0.016) | 0.010 (0.002-0.021) | 0.003 (0.002-0.006) | A:0.5769 B:0.5256 | A:0.9998 B:0.9985 | A:0.9798 B:0.9994 |
| ***Serine peptidases*** | 0.128 (0.073-0.235) | 0.110 (0.070-0.324) | 0.135 (0.091-0.196) | A:0.5769 B:0.9994 | A:0.4298 B:0.9985 | A:0.9798 B:0.9994 |
| ***Carboxypeptidases*** | 0.004 (0.001-0.013) | 0.007 (0.003-0.011) | 0.017 (0.010-0.028) | A:0.5769 B:0.5298 | A:0.9998 B:0.9985 | A:0.9798 B:0.9994 |
| ***Dipeptidyl peptidases*** | 0.001 (0.000-0.009) | 0.001 (0.000-0.003) | 0.005 (0.001-0.026) | A:0.5769 B:0.5256 | A:0.9998 B:0.9985 | A:0.9798 B:0.9994 |
| ***Actin family*** | 0.001 (0.000-0.002) | 0.004 (0.001-0.010) | 0.002 (0.001-0.014) | A:0.5769 B:0.4352 | A:0.9896 B:0.8230 | A:0.9798 B:0.9994 |

Table S26. Difference in human protein classes between visits within the group which outgrew their CMA as determined by Linear Mixed Model (LMM) analysis with outgrowth of CMA, visit and outgrowth of CMA x visit as fixed effects and subject as random effect. Abbreviation: p-adj.: adjusted p-value (p-value corrected for multiple testing using the Benjamini-Hochberg correction). Top 10 features ordered by unadjusted p-value.

| **Outgrowth of CMA – visit 6 months vs baseline** | | | |
|---|---|---|---|
| **Human protein class** | **p-value LMM** | **p-adj. LMM** | **Found by LEfSe** |
| S100 proteins | 0.007 | 0.649 | yes |
| P-ATPases | 0.046 | 1.000 | yes |
| Proline rich proteins | 0.070 | 1.000 | |
| Cell surface glycoproteins | 0.071 | 1.000 | yes |
| Initiation factors | 0.093 | 1.000 | |
| Annexins | 0.107 | 1.000 | |
| Actin binding proteins | 0.126 | 1.000 | |
| Alkaline phosphatases | 0.138 | 1.000 | |
| Transferases | 0.153 | 1.000 | yes |
| Cadherins | 0.240 | 1.000 | |
| **Outgrowth of CMA – visit 12 months vs baseline** | | | |
| **Human protein class** | **p-value LMM** | **p-adj. LMM** | **Found by LEfSe** |
| Alkaline phosphatases | 0.001 | 0.073 | yes |
| Proline rich proteins | 0.002 | 0.073 | yes |
| Superoxide dismutases | 0.004 | 0.133 | yes |
| Ubiquitins | 0.008 | 0.159 | yes |
| Fused gene family | 0.009 | 0.159 | yes |
| Filaggrins | 0.013 | 0.202 | yes |
| Binding protein | 0.016 | 0.205 | yes |
| Ligases | 0.020 | 0.220 | |
| Ferritins | 0.022 | 0.220 | yes |
| Plakins | 0.024 | 0.220 | yes |
| **Outgrowth of CMA – visit 12 months vs 6 months** | | | |
| **Human protein class** | **p-value LMM** | **p-adj. LMM** | **Found by LEfSe** |
| Filaggrins | 0.003 | 0.163 | yes |
| Fused gene family | 0.006 | 0.163 | yes |
| Transthyretin hydroxyisourate hydrolases | 0.007 | 0.163 | |
| Selenium binding proteins | 0.007 | 0.163 | |
| Triosephosphate isomerase | 0.027 | 0.405 | yes |
| Haemoglobins | 0.032 | 0.405 | |
| Phosphorylase enzymes | 0.043 | 0.405 | |
| Initiation factors | 0.044 | 0.405 | |
| Cadherins | 0.048 | 0.405 | yes |
| Fatty acid binding proteins | 0.048 | 0.405 | yes |

Table S27. Difference in human protein classes between visits within the group which did not outgrew their CMA as determined by Linear Mixed Model (LMM) analysis with outgrowth of CMA, visit and outgrowth of CMA x visit as fixed effects and subject as random effect. Abbreviation: p-adj.: adjusted p-value (p-value corrected for multiple testing using the Benjamini-Hochberg correction). Top 10 features ordered by unadjusted p-value.

| **No outgrowth of CMA – visit 6 months vs baseline** | | | |
|---|---|---|---|
| **Human protein class** | **p-value LMM** | **p-adj. LMM** | **Found by LEfSe** |
| Exosomal proteins | 0.002 | 0.149 | yes |
| Cell adhesion proteins | 0.008 | 0.376 | yes |
| Filaggrins | 0.039 | 0.825 | yes |
| Proline rich proteins | 0.039 | 0.825 | yes |
| Actin family | 0.045 | 0.825 | yes |
| Complement C3 like proteins | 0.093 | 1.000 | |
| Serpins | 0.095 | 1.000 | yes |
| Selenium binding proteins | 0.101 | 1.000 | |
| Mucin family | 0.111 | 1.000 | |
| Actin binding proteins | 0.129 | 1.000 | yes |
| **No outgrowth of CMA – visit 12 months vs baseline** | | | |
| **Human protein class** | **p-value LMM** | **p-adj. LMM** | **Found by LEfSe** |
| Tumor suppressor | 0.003 | 0.222 | |
| Cell adhesion proteins | 0.006 | 0.222 | |
| Proline rich proteins | 0.007 | 0.222 | yes |
| Serpins | 0.035 | 0.673 | |
| Neutral alkaline ceramidases | 0.037 | 0.673 | yes |
| Immunoglobulins | 0.060 | 0.746 | |
| Macroglobulins | 0.075 | 0.746 | |
| Calreticulin family | 0.083 | 0.746 | |
| Amylases | 0.088 | 0.746 | yes |
| Triosephosphate isomerase | 0.088 | 0.746 | yes |
| **No outgrowth of CMA – visit 12 months vs 6 months** | | | |
| **Human protein class** | **p-value LMM** | **p-adj. LMM** | **Found by LEfSe** |
| Filaggrins | 0.001 | 0.070 | yes |
| Selenium binding proteins | 0.004 | 0.181 | |
| Exosomal proteins | 0.007 | 0.181 | yes |
| Tumor suppressor | 0.008 | 0.181 | |
| Fatty acid binding proteins | 0.052 | 0.947 | yes |
| Pyridine nucleotide disulphide reductases | 0.090 | 0.947 | yes |
| Immunoglobulins | 0.096 | 0.947 | |
| Ferritins | 0.108 | 0.947 | |
| Peroxidases | 0.112 | 0.947 | |
| Alkaline phosphatases | 0.113 | 0.947 | |

Table S28. Spearman correlation between human (top 10 human protein classes) and microbial (core taxa) proteins. P-values (between brackets) were determined with Monte Carlo permutation (10 000 permutations). P-values below 0.05 are considered significant.

| | ***Bacteroidaceae*** | ***Bifidobacteriaceae*** | ***Lachnospiraceae*** | ***Ruminococcaceae*** | ***Coriobacteriaceae*** | ***Enterobacteriaceae*** | ***Veillonellaceae*** |
|---|---|---|---|---|---|---|---|
| ***Immunoglobulins*** | 0.0253 (0.7871) | 0.1636 (0.0737) | -0.0546 (0.557) | -0.1155 (0.2254) | -0.0180 (0.8512) | -0.0303 (0.7390) | 0.0384 (0.6768) |
| ***Glycoside hydrolases*** | 0.1698 (0.0669) | -0.1131 (0.2216) | 0.0878 (0.3479) | 0.1419 (0.1256) | **0.2732 (0.0038)** | **-0.2042 (0.0282)** | -0.0711 (0.4448) |
| ***Transthyretin/hydroxyisourate hydrolases*** | **0.1934 (0.0373)** | **-0.1980 (0.0313)** | 0.1279 (0.1693) | 0.0191 (0.8361) | **0.4603 (< $10^{-4}$)** | 0.0484 (0.6137) | **0.2968 (0.0010)** |
| ***Proline-rich proteins*** | -0.0426 (0.6503) | -0.1585 (0.0873) | **0.1939 (0.0358)** | **0.2161 (0.0167)** | **0.3513 (0.0001)** | **-0.1953 (0.0341)** | 0.1472 (0.1134) |
| ***S100 proteins*** | **0.2828 (0.0023)** | -0.1613 (0.084) | 0.0496 (0.5894) | 0.1080 (0.2491) | 0.1783 (0.0525) | 0.1789 (0.0544) | 0.1258 (0.1714) |
| ***Secretory proteins*** | **0.2584 (0.0054)** | **-0.2712 (0.0046)** | **0.2508 (0.0082)** | 0.1477 (0.1094) | 0.1593 (0.0839) | 0.0482 (0.6052) | 0.1251 (0.1771) |
| ***Serine peptidases*** | -0.0981 (0.2916) | -0.0143 (0.8755) | -0.0412 (0.6648) | -0.0019 (0.9828) | -0.1071 (0.2498) | 0.0693 (0.4578) | -0.0509 (0.5878) |
| ***Carboxypeptidases*** | **0.1894 (0.0412)** | **-0.2943 (0.0009)** | **0.2541 (0.0068)** | **0.1895 (0.0380)** | **0.4598 (< $10^{-4}$)** | -0.0830 (0.3678) | **0.2274 (0.0148)** |
| ***Dipeptidyl peptidases*** | **0.2471 (0.0079)** | -0.1664 (0.0671) | 0.1328 (0.1544) | **0.2253 (0.0161)** | **0.2869 (0.0015)** | -0.0678 (0.4690) | -0.0237 (0.7903) |
| ***Actin family*** | 0.0685 (0.4579) | -0.1057 (0.2529) | 0.1154 (0.2175) | **0.1941 (0.0378)** | **0.1939 (0.0354)** | -0.1089 (0.2528) | 0.1551 (0.0956) |

Table S29. Significant (p-value ≤ 0.05, bold italic) and marginally significant (0.05 < p-value ≤ 0.1) results from redundancy analysis (RDA) on microbial proteome profiles, microbial proteome functional profiles (KEGG Brite level c), 16S rRNA gene-based taxonomic profiles (family level), protein-based taxonomic profiles (family level) and human protein profiles. adj. p-value: p-value adjusted for multiple testing using the Benjamini-Hochberg correction.

| **Type of profile** | **Baseline** | | | **6 months** | | | **12 months** | | |
|---|---|---|---|---|---|---|---|---|---|
| | **Variable** | **adj. p-value** | **%variance explained by selected variables** | **Variable** | **adj. p-value** | **%variance explained by selected variables** | **Variable** | **adj. p-value** | **%variance explained by selected variables** |
| **Microbial proteome** | none | n/a | n/a | delivery<br>treatment<br>age<br>egg allergy<br>outgrowth CMA<br>number of infections | ***0.003***<br>***0.012***<br>***0.008***<br>***0.003***<br>0.057<br>0.063 | 21.47% | treatment<br>delivery<br>spitting | ***0.001***<br>***0.038***<br>0.081 | 12.08% |
| **Microbial protein functional classes (KEGG Brite level c)** | allergy father<br>age<br>egg allergy | ***0.009***<br>***0.025***<br>***0.017*** | 12.58% | allergy mother<br>delivery<br>age<br>stool colour | ***0.007***<br>***0.012***<br>***0.027***<br>***0.048*** | 17.22% | treatment<br>SCORAD | ***0.003***<br>***0.030*** | 10.07% |
| **16s rRNA gene-based taxonomy** | delivery<br>age | ***0.003***<br>***0.007*** | 9.79% | treatment<br>age<br>egg allergy<br>delivery<br>**outgrowth CMA**<br>vomiting<br>stool colour | ***0.001***<br>***0.001***<br>***0.005***<br>***0.026***<br>***0.018***<br>***0.021***<br>0.051 | 32.56% | delivery<br>gaswind<br>treatment<br>stool colour | ***0.013***<br>***0.029***<br>***0.039***<br>***0.045*** | 16.78% |
| **Protein-based taxonomy** | delivery<br>sibling<br>other allergy | ***0.013***<br>***0.011***<br>***0.036*** | 15.47% | stool colour<br>age | ***0.025***<br>***0.039*** | 9.60% | none | n/a | n/a |
| **Human proteins** | none | n/a | n/a | allergy mother<br>delivery | ***0.016***<br>***0.035*** | 7.08% | spitting | ***0.027*** | 3.66% |

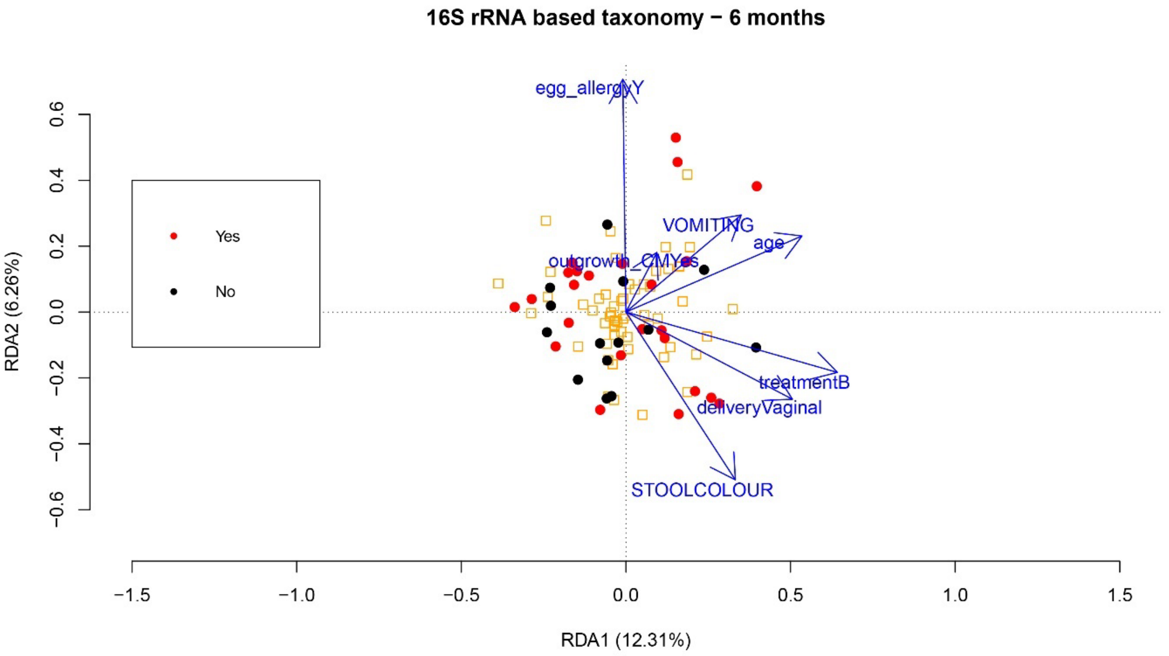

Figure S6. Redundancy analysis (RDA) of the bacterial community in faeces samples for the visit 6 months, coloured by outgrowth of CMA using 16S-rRNA gene based taxonomic profiles at the family level. Arrows indicate features with p-value ≤ 0.1. Red dots: Yes = outgrowth of CMA; black dots: No = no outgrowth of CMA; orange squares: microbial variables (each square represents a family).

Table S30. Results of partial RDA with outgrowth of CMA as explanatory variable, adjusting for other environmental factors. P-value and % variance explained by outgrowth of CMA. Significant results are indicated in bold.

| **Type of profile** | **baseline** | | **6 months** | | **12 months** | |
|---|---|---|---|---|---|---|
| | **p-value** | **%variance explained** | **p-value** | **%variance explained** | **p-value** | **%variance explained** |
| **Proteome** | 0.468 | 3.54% | 0.391 | 5.24% | 0.502 | 5.04% |
| **Microbial protein functional classes (KEGG Brite level c)** | 0.520 | 3.29% | 0.217 | 6.15% | 0.845 | 3.32% |
| **16S rRNA gene-based taxonomy** | 0.634 | 3.05% | **0.044** | **6.37%** | 0.087 | 6.23% |
| **Protein-based taxonomy** | 0.340 | 3.83% | 0.190 | 6.76% | 0.210 | 6.47% |
| **Human proteins** | 0.842 | 2.78% | 0.505 | 4.89% | 0.479 | 5.01% |

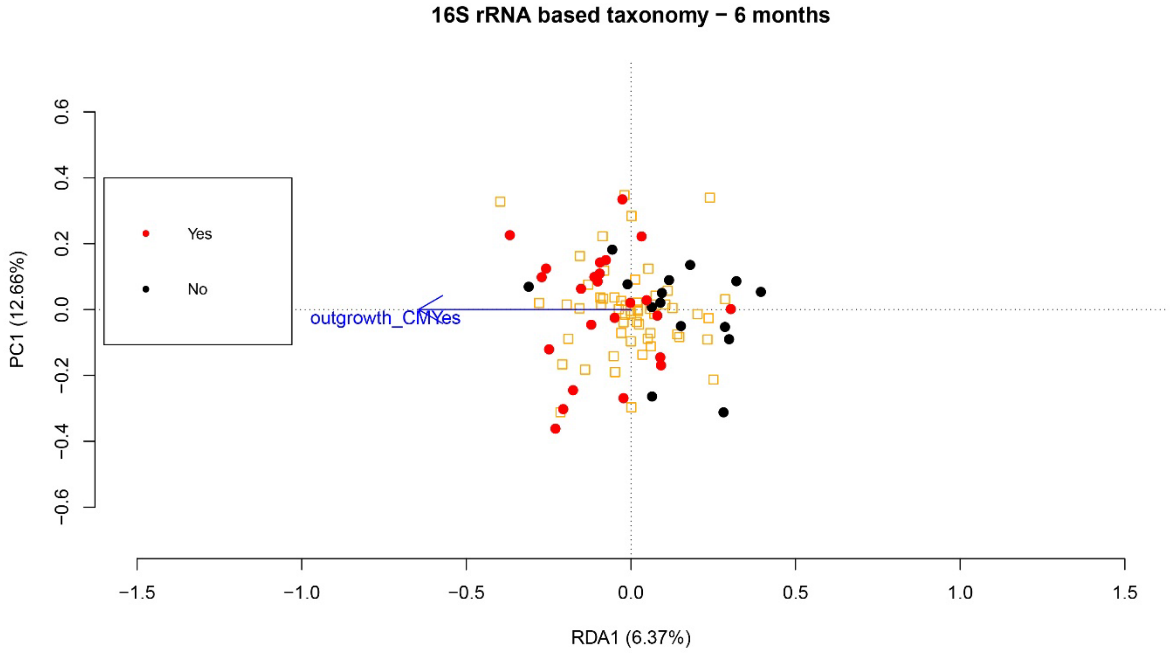

Figure S7. Partial RDA of 16S-rRNA gene based taxonomic profiles at the family level for the visit 6 months, with outgrowth of CMA as explanatory variable and adjusting for other environmental factors. Figure coloured by outgrowth of CMA. Red dots: Yes = outgrowth of CMA; black dots: No = no outgrowth of CMA; orange squares: microbial variables (each square represents a family).

Table S31. Significant (p-value ≤ 0.05, bold italic) and marginally significant (0.05 < p-value ≤ 0.1) results from redundancy analysis (RDA) on microbial proteome profiles and protein-based taxonomic profiles (family level), using human proteins as explanatory variables. adj. p-value: p-value corrected for multiple testing using Benjamini-Hochberg correction.

| **Type of profile** | **Baseline** | | |
|---|---|---|---|
| | **Variable** | **adj. p-value** | **%variance explained by selected variables** |
| **Microbial proteome** | Prolactin-inducible protein | ***0.001*** | |
| | Keratin, type II cytoskeletal 8 | ***0.004*** | 11.86% |
| | Basic salivary proline-rich protein 2 | ***0.004*** | |
| | Basic salivary proline-rich protein 3 | ***0.019*** | |
| **Protein-based taxonomy** | Superoxide dismutase [Cu-Zn] | ***0.001*** | |
| | Immunoglobulin heavy variable 3-15 | ***0.001*** | |
| | Apolipoprotein D | ***0.001*** | |
| | Dihydrolipoyl dehydrogenase, mitochondrial | ***0.002*** | |
| | Putative N-acetylated-alpha-linked acidic dipeptidase | ***0.006*** | |
| | Fatty acid-binding protein, intestinal | ***0.006*** | 50.35% |
| | Serotransferrin | ***0.010*** | |
| | ATP synthase subunit alpha, mitochondrial | ***0.005*** | |
| | Pyruvate kinase PKM | 0.055 | |
| | Heat shock protein HSP 90-alpha | 0.060 | |
| | **6 months** | | |
| | **Variable** | **adj. p-value** | **%variance explained by selected variables** |
| **Microbial proteome** | POTE ankyrin domain family member E | ***0.001*** | |
| | Elongation factor 1-alpha 1 | ***0.002*** | 9.20% |
| | Myeloperoxidase | ***0.003*** | |
| **Protein-based taxonomy** | Intelectin-2 | ***0.001*** | |
| | Immunoglobulin lambda variable 1-51 | ***0.001*** | |
| | Myosin-2 | ***0.003*** | |
| | Meprin A subunit beta | ***0.007*** | 44.67% |
| | Carboxypeptidase B | ***0.014*** | |
| | Fatty acid-binding protein, intestinal | ***0.013*** | |
| | Alpha-1-antitrypsin | ***0.011*** | |
| | Intelectin-1 | ***0.015*** | |
| | Beta-enolase | ***0.035*** | |
| | **12 months** | | |
| | **Variable** | **adj. p-value** | **%variance explained by selected variables** |
| **Microbial proteome** | Bile salt-activated lipase | ***0.001*** | |
| | Chymotrypsin-C | ***0.005*** | |
| | Basic salivary proline-rich protein 3 | ***0.032*** | 15.57% |
| | Ferritin heavy chain | ***0.026*** | |
| | Transketolase | ***0.040*** | |
| **Protein-based taxonomy** | Neutrophil gelatinase-associated lipocalin | ***0.001*** | |
| | Protein S100-A8 | ***0.001*** | |
| | Alpha-amylase 1 | ***0.001*** | |
| | Carboxypeptidase A1 | ***0.001*** | |
| | Putative uncharacterized protein MYH16 | ***0.001*** | |
| | Transketolase | ***0.001*** | |
| | Immunoglobulin heavy constant alpha 2 | ***0.001*** | |
| | Xaa-Pro aminopeptidase 2 | ***0.001*** | |
| | Serpin B6 | ***0.001*** | |
| | Trypsin-1 | ***0.001*** | |
| | Creatine kinase M-type | ***0.001*** | 88.37% |
| | Ferritin heavy chain | ***0.001*** | |
| | Phospholipase A2, membrane associated | ***0.001*** | |
| | Tropomyosin alpha-3 chain | ***0.001*** | |
| | Alpha-actinin-2 | ***0.001*** | |
| | Voltage-dependent anion-selective channel protein 1 | ***0.001*** | |
| | Eukaryotic initiation factor 4A-III | ***0.002*** | |
| | Polymeric immunoglobulin receptor | ***0.006*** | |
| | Keratin, type II cytoskeletal 8 | ***0.014*** | |
| | Pancreatic triacylglycerol lipase | ***0.007*** | |
| | Immunoglobulin kappa variable 1-6 | ***0.008*** | |
| | Immunoglobulin kappa constant | ***0.014*** | |
| | Ferritin light chain | ***0.024*** | |
| | Dihydrolipoyl dehydrogenase, mitochondrial | ***0.049*** | |

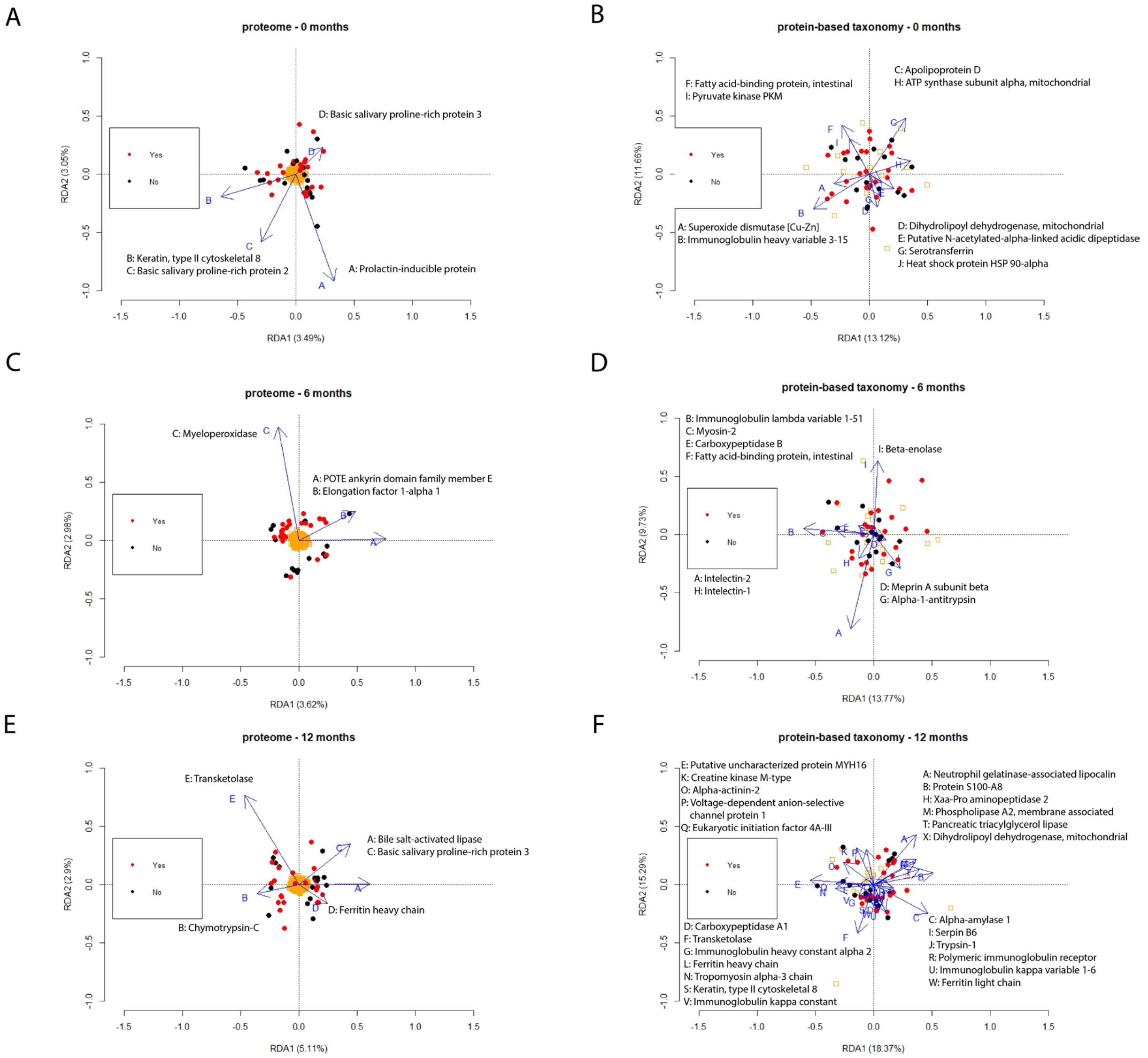

Figure S8. RDA using human proteins as explanatory variables, and coloured by outgrowth of CMA. Red dots: Yes = outgrowth of CMA; black dots: No = no outgrowth of CMA; orange squares: microbial variables (each square represents a microbial protein group (left) or family (right)). (A) RDA of proteome profiles at baseline visit; (B) RDA of protein-based taxonomy profiles (family level) at baseline visit; (C) RDA of proteome profiles at visit 6 months; (D) RDA of protein-based taxonomy profiles (family level) at visit 6 months; (E) RDA of proteome profiles at visit 12 months; (F) RDA of protein-based taxonomy profiles (family level) at visit 12 months.

Table S32. Protein functional classes related to metabolic pathways increased between visits in the group with outgrowth of CMA, but not in the other group as determined by LEfSe analysis. Second column: displays from which families these proteins originate. Third column: displays between which visits the increase was observed. Fourth column: families that show the same behaviour over visits than the protein classes in the first column. Families that increase in the same way as the protein classes (Figure 5) are indicated in bold.

| **Protein functional class** | **Families in data set** | **Increase between visits … (Figure 6)** |
|---|---|---|
| Selenocompound metabolism | *Bifidobacteriaceae, Lachnospiraceae, Ruminococcaceae* | Baseline and 6 months<br>Baseline and 12 months |
| Aminoacyl-tRNA biosynthesis | *Bifidobacteriaceae, Lachnospiraceae* | Baseline and 6 months<br>Baseline and 12 months |
| Amino sugar and nucleotide sugar metabolism | *Bifidobacteriaceae, Lachnospiraceae, Enterococcaceae* | Baseline and 6 months<br>6 months and 12 months |
| Cysteine and methionine metabolism | *Bacteroidaceae, Bifidobacteriaceae, Ruminococcaceae, Lachnospiraceae, Clostridiaceae* | Baseline and 6 months<br>Baseline and 12 months |
| Galactose metabolism | *Bifidobacteriaceae, Lachnospiraceae* | Baseline and 6 months |
| Pentose and glucuronate interconversions | *Bacteroidaceae, Clostridiaceae, Lachnospiraceae, Bifidobacteriaceae* | Baseline and 12 months |
| Pyruvate metabolism | *Bacteroidaceae, Enterobacteriaceae, Enterococcaceae, Clostridiaceae, Oscillospiraceae, Lachnospiraceae, Prevoltellaceae, Bifidobacteriaceae, Ruminococcaceae, Veillonellaceae,* ***Rikenellaceae*** | Baseline and 12 months |
| Porphyrin metabolism | *Bifidobacteriaceae, Lachnospiraceae* | Baseline and 12 months |
| Nitrogen metabolism | *Lachnospiraceae, Bacteroidaceae, Clostridiaceae, Ruminococcaceae, Bifidobacteriaceae, Veillonellaceae* | Baseline and 12 months |
| Fructose and mannose metabolism | *Bacteroidaceae, Streptococcaceae, Bifidobacteriaceae, Lachnospiraceae,* ***Coriobacteriaceae****, Enterococcaceae, Ruminococcaceae, Clostridiaceae* | Baseline and 12 months |
| Beta-alanine metabolism | *Lachnospiraceae, Veillonellaceae* | Baseline and 12 months |
| Fatty acid degradation | *Clostridiaceae, Bifidobacteriaceae, Lachnospiraceae, Oscillospiraceae, Veillonellaceae* | Baseline and 12 months |
| Propanoate metabolism | *Enterobacteriaceae, Clostridiaceae, Lachnospiraceae, Oscillospiraceae, Enterococcaceae, Bacteroidaceae, Bifidobacteriaceae, Veillonellaceae* | Baseline and 12 months |
| Glycerolipid metabolism | *Lachnospiraceae* | Baseline and 12 months |
| Streptomycin metabolism | *Bifidobacteriaceae, Lachnospiraceae* | Baseline and 12 months |
